# Diel cycle of lanthanide-dependent methylotrophy by TMED127/Methylaequorales bacteria in oligotrophic surface seawater

**DOI:** 10.1101/2025.06.11.659106

**Authors:** Jennifer B. Glass, Leilani N. Warters, Abdulaziz M. Alajlan

## Abstract

Methanol, the simplest alcohol, has long been recognized as a key energy and carbon source for soil and plant-associated bacteria and fungi, and is increasingly recognized as an important substrate for marine bacteria. Lanthanide-dependent methanol dehydrogenases (encoded by the gene *xoxF*) are now recognized as the key catalysts for methylotrophy in many environments, yet the identity of the most transcriptionally active methylotrophs in open ocean waters (“Clade X”) has remained elusive. Here we show that “Clade X” methylotrophs belong to the deep-branching alphaproteobacterial order TMED127, which we propose be renamed ‘Methylaequorales’; ‘methyl’ for ‘methylotrophic metabolism’ and ‘aequor’ for ‘ocean surface’, as these bacteria are most transcriptionally active near the sea surface. TMED127/Methylaequorales are present in surface waters of tropical and subtropical oceans throughout the global ocean. They have small, streamlined genomes (∼1.5 Mb), and appear to be obligate methylotrophs that use the serine cycle for carbon assimilation. They display a diel pattern of *xoxF5* and glucose dehydrogenase (*gdh*) transcription, peaking in the late afternoon, in oligotrophic surface water of the Sargasso Sea. Several other highly transcribed genes of unknown function had no homologs outside of TMED127/Methylaequorales genomes. Our findings illuminate an overlooked marine methylotrophic bacterium and predict a diel cycle of methanol production in surface seawater by an unknown pathway.

**Importance:** Methanol metabolism is increasingly recognized as an important process in the marine carbon cycle, yet the identity and metabolism of the microorganisms mediating methylotrophy in the open ocean have remained unknown. This study reveals that bacteria in the TMED127 order of Alphaproteobacteria, proposed new name of “Methylaequorales”, abundantly transcribe the key gene for lanthanide-dependent methylotrophy in oligotrophic surface waters of the world’s oceans. TMED127/Methylaequorales likely require methanol as a carbon and energy source and display a diel pattern of transcription of key genes for methylotrophy that peaks in the late afternoon. These findings motivate future studies on the mechanisms of methanol production in surface seawater.

## Introduction

Methanol (CH_3_OH) is a simple alcohol that is used by methylotrophic bacteria as an energy and carbon source (1, 2). The first step of bacterial methylotrophy, oxidation of methanol to formaldehyde, is mediated the periplasmic enzyme pyrrolo-quinoline quinone (PQQ) methanol dehydrogenase, which requires either calcium or a light lanthanide elements (lanthanum, cerium, praseodymium, and neodymium) as its cofactor (3). Lanthanide-dependent methanol dehydrogenases are encoded by the gene *xoxF* and calcium-dependent methanol dehydrogenases are encoded by the gene *mxaF*. Formaldehyde produced by methanol dehydrogenase is then further oxidized to yield energy for cellular growth and/or assimilated into cellular carbon via the ribulose monophosphate cycle or the serine cycle (4).

Methylotrophy appears to be widespread in terrestrial and marine ecosystems. Plants make methanol from cell wall pectins using pectin methylesterase (5), and terrestrial plant-associated methylotrophic bacteria have been extensively studied for their agricultural importance (6). Methanol is also produced by marine phytoplankton via unknown mechanism(s) (7, 8) and can also be produced abiotically by photodegradation of methoxy aromatic groups in lignin (9) or reaction of methyl peroxy radical with hydroxyl radicals (10). Some marine bacteria secrete pectin methyltransferases, possibly producing methanol as a byproduct (11–13). Most methanol emitted to the atmosphere is rapidly deposited back to the ocean surface (14–16), where it is rapidly consumed by bacteria for energy and, to a lesser degree, for carbon assimilation (17–21).

Although calcium is nine orders of magnitude more abundant than lanthanides in seawater, marine microbial methanol oxidation is primarily lanthanide dependent, based on the prevalence and transcriptional activity of *xoxF* compared to *mxaF* (22). The dominant types of *xoxF* in most natural environments are *xoxF5*, found in many orders of Alpha- and Gammaproteobacteria, and *xoxF4*, found in Methylophilaceae (23, 24). Selective drawdown of light lanthanides relative to heavy lanthanides in seawater is associated with biological activity (25–27). In coastal waters, methylotrophic bacteria are dominated by Methylophilaceae (also known as OM43; 28, 29-34), which possess *xoxF4,* and Methylophagaceae, which possess both *xoxF5* and *mxaF* (21, 35–37). Omics studies have revealed a much broader diversity of marine bacteria with the capacity for lanthanide-based methylotrophy than previously realized (24, 38–41). A recent study reported an unidentified alphaproteobacterial “Clade X” of *xoxF5* genes as having highest transcriptional activity in Tara Oceans seawater samples (22). Here we investigated the identity, predicted metabolism, and transcriptional activity of methylotrophic bacteria in oligotrophic waters representative of the vast majority of the ocean’s surface to resolve the identity and metabolism of the elusive “Clade X” methylotroph.

## Results

### Most highly transcribed xoxF5 genes in oligotrophic surface seawater belong to the alphaproteobacterial order TMED127

RNA-Seq datasets in the Department of Energy (DOE) Joint Genome Institute Integrated Microbial Genomes and Microbiomes (JGI IMG/M) database (42) were searched for PQQ-dependent methanol dehydrogenases (see Materials and Methods for details). We found extremely high transcription of lanthanide-dependent methanol dehydrogenase *xoxF5* genes (up to 2056 RPKM) in oligotrophic surface seawater from three different ocean basins (North Atlantic, South Atlantic, and Indian Ocean (39, 43–45); **Table S1)**. In a dataset from the Bermuda Atlantic Time-series Study (BATS; Sargasso Sea, North Atlantic Ocean) with free-living (SX) and particulate (LSF) size fractions, *xoxF5* transcripts were almost exclusively present in free-living fractions (**Table S1)**, suggesting that active methylotrophs are free living rather than particle associated. The most highly transcribed *xoxF5* genes had high identity (93-100%) to genes from genus GCA-002690875 and GCA-2691245 of the uncultured marine alphaproteobacterial order TMED127 (GTDB taxonomy; 46), previously known as “Ricksettsiales” in NCBI taxonomy **(Fig. 1A; Table S1)**. Multiple identical full-length sequences in TMED127 genus GCA-2691245 were recovered **(Table S1)**, motivating further analysis of this genus (see below).

**Figure 1.**
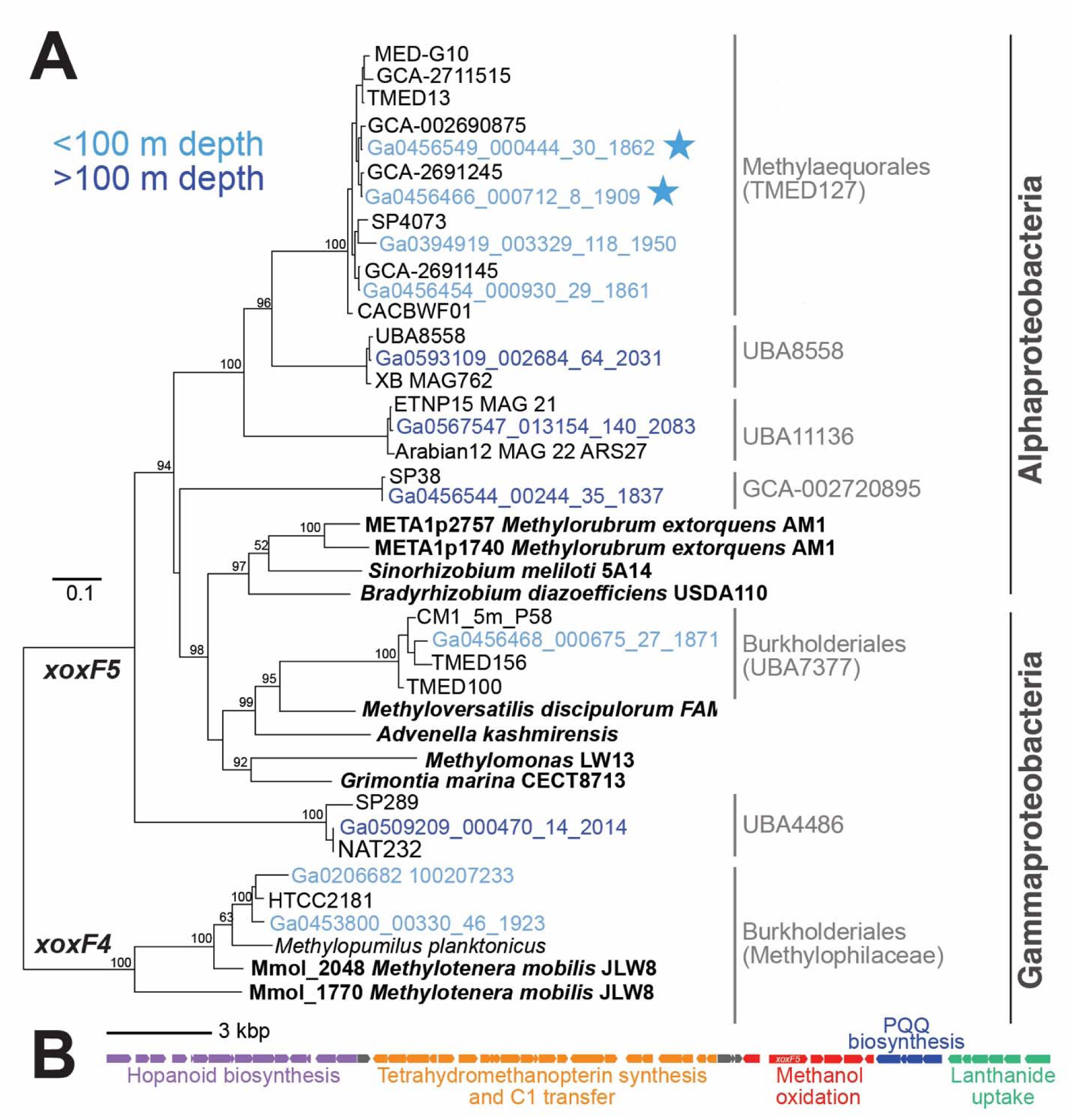
Lanthanide-dependent methanol dehydrogenase (XoxF) phylogeny and “methylotrophy island” in TMED127 genomes. (A) Maximum likelihood phylogenetic tree of PQQ-methanol dehydrogenases XoxF5 sequences, with XoxF4 sequences as outgroup. Sample depths are colored (light blue: 0-100 m depth; dark blue: >100 m depth). All JGI IMG sequences (beginning with “Ga”) are available in **Table S1**, and those with >100 RPKM in any sample JGI IMG are on the phylogeny. Classes are in dark gray. Orders containing highly transcribed marine JGI IMG samples are labeled in light gray according to GTDB taxonomy. TMED127 nodes are labeled by genera; all other nodes are labeled by SAG/MAG isolate name or species name. The stars mark the JGI IMG seawater samples with highest transcription from the Sargasso Sea. Sequences encoding biochemically characterized enzymes (bold) are from refs. (23, 24, 49). The two Burkholderiales clades are designated by family (Methylophilaceae) and genus (UBA7377). (B) “Methylotrophy island” (∼45 kbp) in TMED127 genomes (drawn based on isolate CM1_5m.P40), with the following modules: hopanoid biosynthesis (CM15mP40_03190-03050), tetrahydromethanopterin synthesis and C1 transfer (CM15mP40_03030-02810), methanol oxidation (CM15mP40_02770-02720), PQQ biosynthesis (CM15mP40_02710-02680), and lanthanide uptake (CM15mP40_02670-02620). Gray indicates hypothetical proteins.

TMED127 *xoxF5* genes form a monophyletic clade with other Alphaproteobacteria *xoxF5* genes from the orders UBA8558 and UBA11136 that are transcribed in deeper seawater samples (10-150 RPKM; **Fig. 1A; Table S1**). BLAST searches of metatranscriptomes from the Eastern Tropical North Pacific oxygen deficient zone (47) revealed that UBA11136 *xoxF5* transcripts are abundantly transcribed (up to 550 RPKM) in anoxic waters **(Table S2)**, supporting the hypothesis that UBA11136 oxidizes methanol in oxygen deficient zones (48). Other Alphaproteobacteria (GCA-002720895) and Gammaproteobacteria (Burkholderiales (genus UBA7377) and UBA4486) also transcribed *xoxF5* genes in globally distributed samples and across a range of depths (10-500 RPKM; **Fig. 1A; Table S1**), consistent with previous reports (38, 41). As predicted, Methylophilaceae/OM43 *xoxF4* transcripts (1-350 RPKM) were present in coastal waters **(Fig. 1A; Table S1)**.

### TMED127 genomes are small and contain “methylotrophy island”

There are 64 TMED127 SAG/MAGs in GTDB (RS207 and RS226), originating from seawater samples of the Atlantic Ocean, Pacific Ocean, Mediterranean Sea, Red Sea, and Brisbane River **(Table S3)**. These include three MAGs comprising “Clade X” in ref. (22): MED715, MED652, and MED657 (**Table S3)**. TMED127 SAG/MAGs are small (∼1.5 Mb estimated complete size) and contain a ∼45 kbp “methylotrophy island” with genes for methanol oxidation, tetrahydromethanopterin synthesis and C1 transfer, lanthanide uptake (*lutABEFCG* (50)), and pyrroloquinoline quinone (PQQ) biosynthesis (**Fig. 1B).** Upstream of the “methylotrophy island” is a gene cluster encoding the hopanoid biosynthetic pathway implicated in structural ordering of the outer membrane (51, 52). We identified the “methylotrophy island” in 44 of the 64 TMED127 SAG/MAGs, spanning eight genera **(Table S3)**, including the three MAGs previously shown to comprise “Clade X”, suggesting that the “methylotrophy island” is present throughout the TMED127 order. Other TMED127 SAG/MAGs appeared to have the same gene synteny but had truncated *xoxF5*-containing contigs.

### Diel transcriptional rhythm of methanol and glucose dehydrogenases in TMED127

A previous metatranscriptomic time series at BATS found that ∼10% of all genes had diel transcriptional patterns (44). To assess transcription of TMED127 genes, we mapped transcripts from surface seawater (5 m depth) from the previous study, sampled every 4 hours over 5 days **(Table S4)**, to most complete TMED127 SAG from genus GCA-2691245 (AG-892-F10; estimated 90% completeness; 0.01% contamination (53); **Fig. 1A; Table S1**). Throughout the time series, *xoxF5* was consistently the most abundant transcript, comprising 16 to 57% (1.6 to 5.7 × 10^5^ TPM) of AG-892-F10 transcripts **(Table S5)**. The *xoxF5* gene displayed a diel transcriptional cycle, consistently peaking in the late afternoon (16:00-20:00; **Fig. 2**). A PQQ-dependent glucose dehydrogenase (*gdh*), which likely produces the lanthanide chelator D-glucono-1,5-lactone, showed the same transcriptional pattern at roughly an order of magnitude lower abundance **(Fig. 2; Table S5)**. Five other genes were transcribed at >10^4^ TPM but not did not show a temporal pattern; these included proteorhodopsin, two hypothetical proteins (AG-892-F10.CDS.256 and AG-892-F10.CDS.257), the phosphate ABC transport substrate-binding protein (*pstS*), and a DoxX-like family protein (**Table S5)**. Homologs of AG-892-F10.CDS.256 and AG-892-F10.CDS.257 (34% lysine), were only present in other TMED127 SAGs/MAGs, whereas the DoxX-like integral membrane family protein had homologs in other marine Alphaproteobacteria and Methylophilacaeae/OM43.

**Figure 2.**
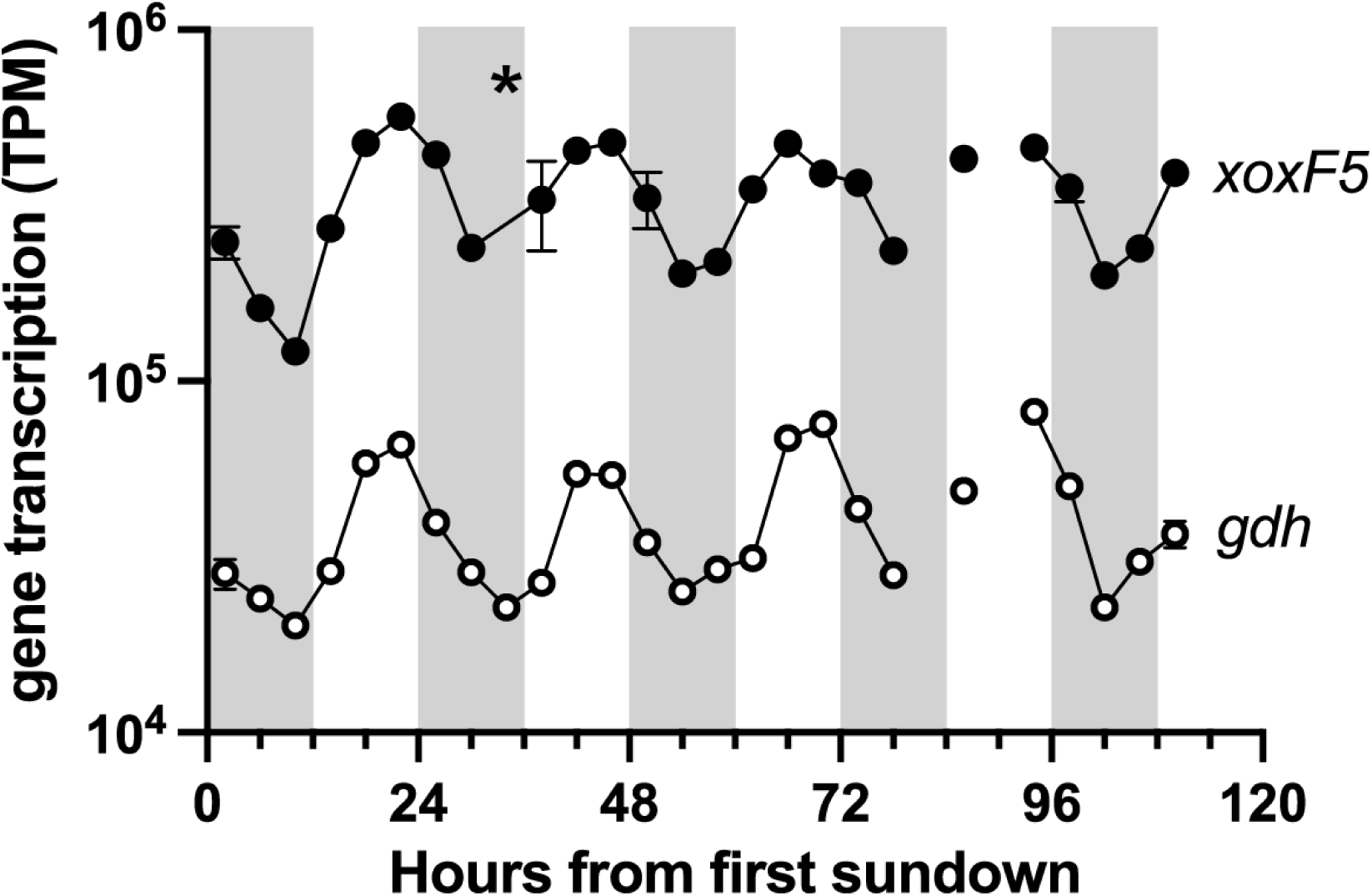
Diel transcription of TMED127 lanthanide-dependent methanol dehydrogenase (*xoxF5*) and glucose dehydrogenase (*gdh*) at Bermuda Atlantic Time-series Study (BATS) site, Sargasso Sea, North Atlantic Ocean. The asterisk represents a single anonymously high sample. Error bars (standard deviation) are shown for the nine time points with multiple metatranscriptomes. Missing lines are due to lack of samples for two timepoints.

### TMED127 are likely obligate methylotrophs

Genomic analysis suggests that TMED127 are obligate methylotrophs. TMED127’s core methylotrophic modules match those of the well-studied plant-associated facultative methylotroph *Methylorubrum extorquens* AM1 (54, 55); roughly two-thirds (956/1535) of AG-892-F10’s genes have homologs in AM1 **(Tables S5)**. Based on extensive previous studies of AM1 metabolism (56, 57), we predict that TMED127 uses the same methylotrophy pathways: after methanol oxidation via XoxF, formaldehyde is oxidized to formate via the tetrahydromethanopterin C1 transfer pathway, which is then either oxidized to CO_2_ by tungsten-containing formate dehydrogenase for regeneration of NAD(P)H or assimilated into biomass via the serine and ethylmalonyl-CoA pathways **(Fig. 3; Table S5)**. TMED127 genes without AM1 homologs include several genes common in marine proteobacteria (e.g., proteorhodopsin for light-driven proton pumping and nickel-dependent superoxide dismutase **(Fig. 3))**. While TMED127’s small (∼1.5 Mb) genome has only one copy of most key functional genes, AM1’s large (6.9 Mb) genome has multiple copies of many key functional genes. AM1 also has a full TCA cycle, enabling its growth on multiple-carbon compounds. Although we cannot be certain of gene absence in incomplete genomes, none of the 64 SAGs/MAGs was found to possess the 2-oxoglutarate dehydrogenase complex, suggesting that they have incomplete TCA cycles characteristic of obligate methylotrophs (58).

**Figure 3.**
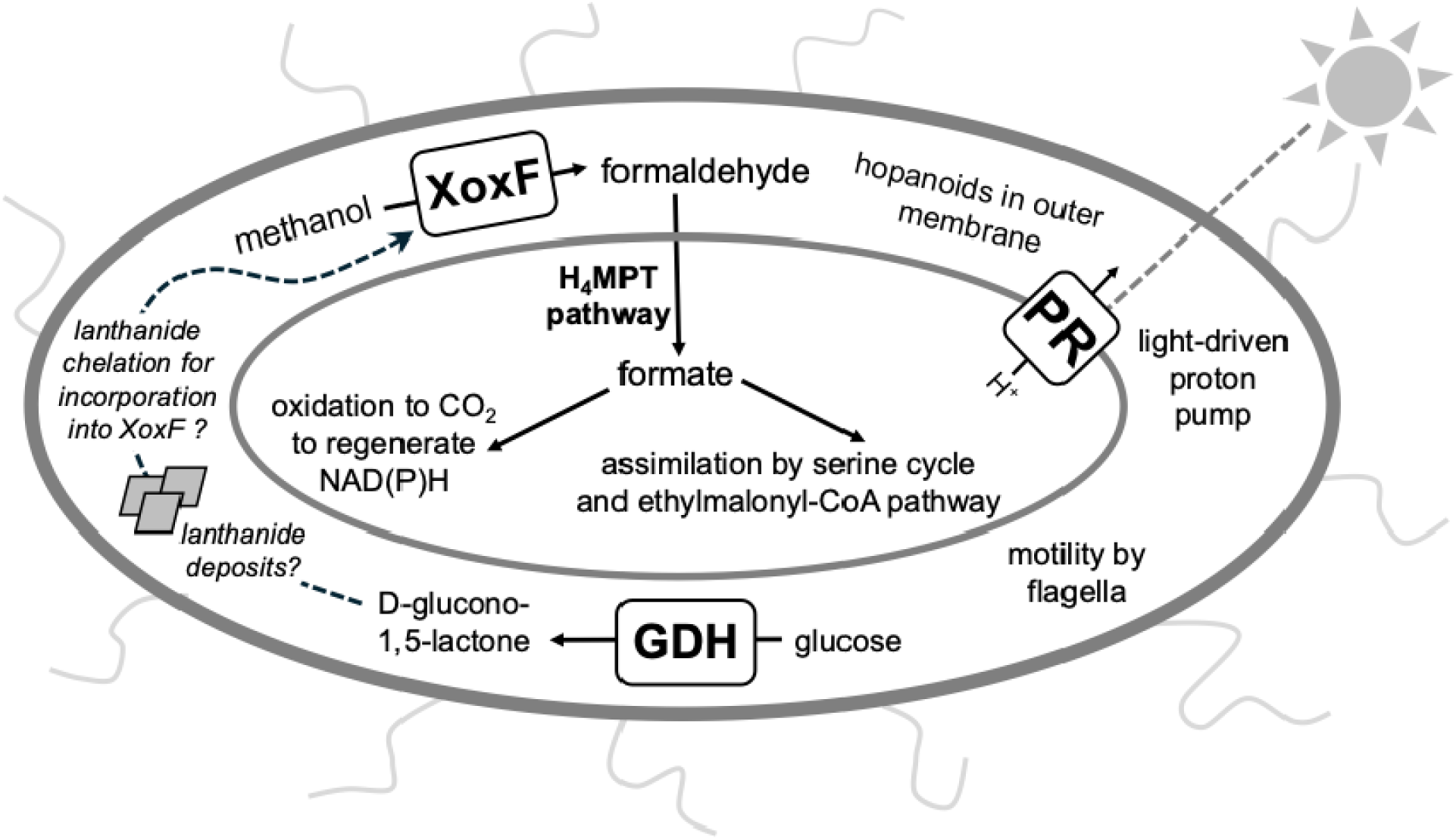
Schematic of key metabolisms and functions in TMED127/Methylaequorales based on genomic reconstruction. Three of the most highly transcribed proteins are shown (PQQ-dependent methanol dehydrogenase (XoxF), PQQ-dependent glucose dehydrogenase (GDH), and proteorhopsin (PR)). Question marks represent hypothetical lanthanide deposits and lanthanide chelation by D-glucono-1,5-lactone, which remains unconfirmed.

### Description of proposed TMED127 names

We propose that the species studied in the present study (AG-892-F10, genus GCA-2691245) be named *Methylaequorum cenaphilum* gen. nov., sp. nov. : Me.thy.lae.quo’rum N.L. neut. n. *methyl*, pertaining to the methyl group; L. neut. n. *aequor*, surface of the sea; N.L. neut. n. *Methylaequorum*, methyl (group oxidizing) organism [of the] sea surface; ce.na.phi’lum L. fem. n. *cena*, the principal meal of the day in ancient Roman culture, originally taken in the afternoon; N.L. masc. adj. suff. *-philus*, loving; N.L. neut. adj. *cenaphilum*, late afternoon meal loving. At the family level, Me.thy.lae.quo.ra’ce.ae. N.L. neut. n. *Methylaequorum*, referring to the type genus Methylaequorum; *-aceae*, ending to denote a family; N.L. fem. pl. n. *Methylaequoraceae*, the Methylaequorum family. At the order level, Me.thy.lae.quo.ra’les N.L. neut. n. *Methylaequorum*, referring to the type genus Methylaequorum; *-ales*, ending to denote an order; N.L. fem. pl. n. *Methylaequorales*, the Methylaequorum order.

### TMED127/Methylaequorales are a deep branching alphaproteobacterial order that are widespread in the global oceans

We constructed a phylogeny using the conserved protein NADH ubiquinone oxidoreductase subunit L (NuoL) as in Cevallos and Degli Esposti (59). As previously reported (60), the TMED127 order is deep-branching, along with the orders Puniceispirillales, TMED109, and Rickettsiales **(Fig. 4A)**. To date, the TMED127/Methylaequorales order contains only one family (TMED127/Methylaequoraceae) and twelve genera **(Fig. 4A)**. The three genera with highest relative abundance in marine metagenomes (up to ∼4% relative abundance each) were TMED13 (0-48° N/S latitude; **Fig. 4B; Table S6)**, GCA-2691245 (*Methylaequorum*; 0-45° N/S latitude; **Fig. 4C; Table S7)**, and SP4073 (16-46° N/S latitude; **Fig. 4D; Table S8).** TMED127/Methylaequorales were not observed at relative abundances of >0.1% above 50° N/S latitude.

**Figure 4.**
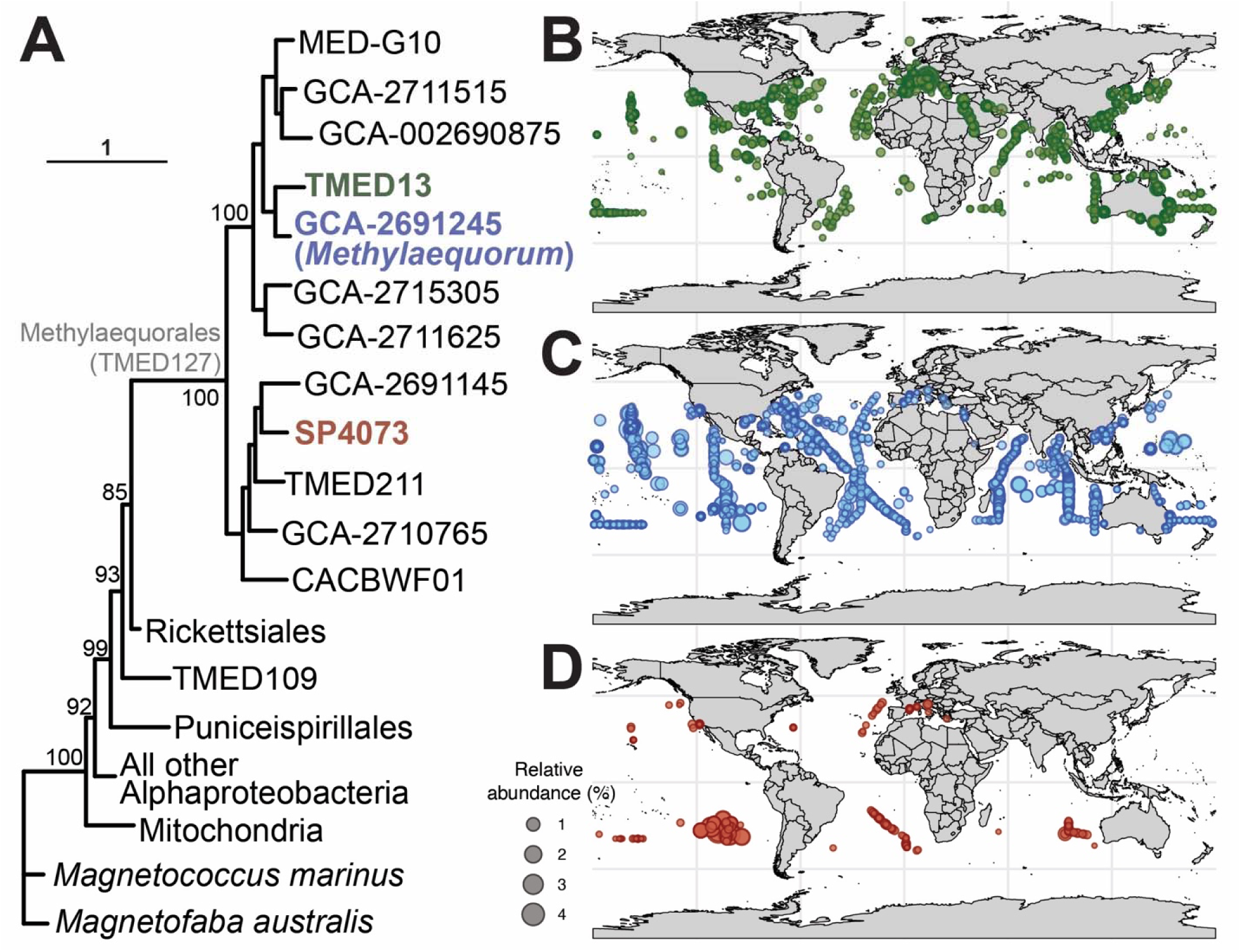
Alphaproteobacterial phylogeny, geographic distribution, and relative abundance of TMED127/Methylaequorales genera. (A) Phylogeny was constructed using the alphaproteobacterial phylogenetic marker NADH ubiquinone oxidoreductase subunit L (NuoL). TMED127/Methylaequorales genera are labelled. Scale bar represents substitutions per site. Colors of genera on phylogeny correspond to colors on B-D panels. (B) Relative abundance bubble plot of genus TMED13 overlaid on global map. (C) Relative abundance bubble plot of genus GCA-2691245 (*Methylaequorum*) overlaid on global map. (D) Relative abundance bubble plot of genus SP4073 overlaid on global map.

## Discussion

This study illuminates the identity, metabolism, and diel rhythms of TMED127 (proposed name: Methylaequorales (order); Methylaequoraceae (family)), a deep-branching order of marine alphaproteobacteria (60) that can comprise up to ∼10% of metagenomic reads in surface seawater. Although confirmation of their physiology and metabolism will require isolation and culture experiments, TMED127/Methylaequorales appear to be free-living obligate methylotrophs that are widespread in global oceans, particularly in tropical and subtropical latitudes. This order is highly transcriptionally active, with a diel cycle of *xoxF5* transcription peaking in the late afternoon in surface water of the Sargasso Sea.

The source of methanol to TMED127/Methylaequorales is unknown. It may be produced as a waste product by eukaryotic phytoplankton and/or cyanobacteria, as is known to occur in culture experiments (7, 8), and via photochemical or enzymatic degradation of other organic molecules produced by phytoplankton in surface seawater (9–12). Peak *xoxF5* transcription in the late afternoon suggests that methylotrophy by TMED127/Methylaequorales is subject to temporal niche partitioning (61), and that methanol supply may be highest in the late afternoon. Seawater carbohydrate concentrations are highest in the afternoon, likely due to passive leakage of uncharged molecules during “photosynthetic overflow” from phytoplankton (62). It is possible that similar methanol leakage from phytoplankton also peaks in the afternoon, but further study of methanol production mechanisms in seawater is required.

PQQ-dependent glucose dehydrogenase (GDH), a periplasmic protein that oxidizes D-glucose to D-glucono-1,5-lactone using calcium as a cofactor (63), showed a synchronous transcriptional pattern to *xoxF5* in TMED127/Methylaequorales, rising throughout the day and falling at night. D-glucono-1,5-lactone is a strong metal chelator that can solubilize calcium phosphate minerals (64) and may function as a lanthanide chelator in methylotrophic bacteria (65). Crystalline accumulations of lanthanides are present in the periplasm (65, 66) and cytoplasm (50) of terrestrial alphaproteobacterial methylotrophs. We hypothesize that D-glucono-1,5-lactone production by GDH releases lanthanides from intracellular storage deposits for incorporation into XoxF **(Fig. 3)**; this hypothesis will require future testing once bacterial isolates are available.

The full complement of lanthanide acquisition and storage pathways in TMED127/Methylaequorales also requires more study. We identified genes encoding the high affinity lanthanide uptake transport system (LutABEFCG; 50), but did not find homologs of genes for “lanthanophore” ligands (67) nor lanthanum storage proteins such as lanmodulin (68) and lanpepsy (69). Given the extremely low (picomolar) lanthanide concentrations in seawater (70, 71), it seems likely that these bacteria have additional lanthanide scavenging strategies, possibly involving one or more of the highly transcribed genes of unknown function.

Our findings suggest that TMED127/Methylaequorales may be the key player in lanthanide-dependent methylotrophy in oligotrophic seawater. More studies are needed to determine whether a diel cycle of methanol production occurs in surface seawater and the quantitative importance of TMED127/Methylaequorales in the global carbon and lanthanide cycles.

## Materials and Methods

### Identifying and quantifying *xoxF* gene transcription in seawater samples

A python code (GeneSweeper; available at https://github.com/GlassLabGT/Python-scripts) was used to search for all possible product names for the target protein, PQQ-dependent methanol dehydrogenase: (“PQQ-dependent dehydrogenase (methanol/ethanol family)”, “methanol dehydrogenase (cytochrome c) subunit 1”, “methanol dehydrogenase (cytochrome)”, “alcohol dehydrogenase (cytochrome c)”, “glucose dehydrogenase”, “quinohemoprotein ethanol dehydrogenase”, and “quinoprotein glucose dehydrogenase”) in CSV files (including gene ID, product name, scaffold ID, gene sequence length, and read count) for all coastal and marine datasets in Joint Genome Institute’s (JGI) Integrated Microbial Genome (IMG) RNASeq Studies database as of May 2025. Only unamended seawater samples were processed; enrichments or laboratory cultures were removed. The total read count of each metatranscriptome was obtained from the JGI IMG RNASeq Studies database. Gene transcription was then converted to reads per kilobase per million mapped reads (RPKM). Amino acid sequences were obtained from reference metagenomes using the JGI IMG “Find Genes” function with the Gene ID as input. Each amino acid sequence was then queried against the NCBI nr database by BLASTP search (May 2025). Only full-length gene sequences were kept for further processing. The top NCBI hit was matched to its GTDB taxonomy (release 10-RS226) by searching GTDB (46) for the GCA assembly of the SAG/MAG.

### XoxF phylogeny

Sequences (*n* = 50; 550 gap-free sites) were aligned using the MAFFT online server with the L-INS-i method (72). A maximum likelihood phylogeny with 1000 bootstraps was constructed in W-IQ-tree (73, 74) using the LG+F+I+G4 model based on ModelFinder results (75) with ultrafast 1000 bootstraps (73). The phylogeny was visualized in iTOL (76). The alignment fasta file (XoxF_alignment.fasta) is available as a supplemental data set.

### Transcription of *xoxF* in Eastern Tropical North Pacific oxygen minimum zone

Magic-Basic Local Alignment Search Tool (Magic-BLAST) (77) was used with default parameters to search ETNP ODZ metatranscriptomes (PRJNA727903; 47) for *xoxF* genes from UBA11136 MAGs using DNA sequence of *xoxF5* from Alphaproteobacteria bacterium isolate ETNP15_MAG_21 (MDP6304722) as the query. Read hits were normalized to reads per kilobase million (RPKM).

### TMED127 SAG/MAG analysis and mapping to metatranscriptomes

All SAG/MAGs in the Alphaproteobacteria order TMED127 in GTDB release 10-RS226 were collected into a BV-BRC genome group, which was used to search for product names via “Proteins” tab and for metabolic pathways via “Pathways” tab (78). The TMED127 SAG AG-892-F10 (estimated 90% completeness; 0.01% contamination; 53) was chosen for further study because it was the most complete SAG/MAG from the genus GCA-2691245, which contained the most abundant *xoxF5* transcripts in surface seawater (see text). Additional analysis was performed in KBase (79). SAG AG-892-F10 (NCBI WGS Project CACMWI01; genome assembly GCA_902617375.1) and *Methylorubrum extorquens* AM1 (NCBI Reference Sequence NC_012808.1) were imported into KBase, and the KBase program “Compare Two Proteomes” was used to find homologs. RNA-Seq data (JGI GOLD Study ID Gs0161320; **Table S4**) were imported into KBase and aligned to SAG AG-892-F10 using the Bowtie2 app (80). The aligned RNA-Seq data was then assembled into a set of transcripts quantified by transcripts per million (TPM) using the StringTie v.2.1.5 (81) app.

### Alphaproteobacteria NuoL phylogeny

A phylogeny of alphaproteobacterial NADH ubiquinone oxidoreductase subunit L (NuoL) and mitochondrial ND5 marker proteins (*n* = 146; 510 gap-free sites) were aligned with MAFFT online server with the L-INS-i method (72). Sequences from two species of c Magnetococcia were used as the outgroup. A maximum likelihood phylogeny with 1000 bootstraps was constructed in W-IQ-tree (73, 74) using the LG+F+I+G4 model based on ModelFinder results (75) with ultrafast 1000 bootstraps (73). The phylogeny was visualized in iTOL (76). The alignment fasta file (NuoL _alignment) is available as a supplemental data set.

### Gene neighborhood

The methylotrophy gene neighborhood was generated using the EFI Gene Neighborhood Tool (48) with single sequence BLAST of the UniProt database with NCBI accession GIR25668 as the query.

### Distribution and relative abundance of TMED127 genera

The twelve TMED127 genera were individually searched for using Sandpiper (82) was searched for used to obtain csv files with relative abundance, latitude, and longitude data. Samples with <0.1% relative abundance of each TMED127 genus were removed. Global maps were overlaid with relative abundance data for the three most abundant genera in R using gglot2 (83).

## Supporting information

XoxF_alignment

NuoL_alignment

## Data Availability

The KBase bioinformatic pipeline is available at https://narrative.kbase.us/narrative/214786. SAG AG-892-F10 is available as NCBI BioSample SAMEA6069714 (SRA: ERS3870829; GCA_902617375.1). RNA-Seq data for the BATS timeseries is available as JGI GOLD Study ID Gs0161320. The TMED127 BV-BRC genome group is available at https://www.bv-brc.org/workspace/jbglass1@patricbrc.org/TMED127%20genomes. The newly proposed names have been submitted to the SeqCode Registry as accession seqco.de/r:_v7atfaf.

## Acknowledgments

We thank Naomi Gilbert, Steven Wilhelm, Matthew Sullivan, and David Walsh for providing access to sequences and metadata. We thank Ann Pearson, Paul Carini, and Steven Wilhelm for helpful discussions. We thank William Whitman and Luis-Miguel Rodríguez Rojas for curation assistance with submission to SeqCode Registry.

