## Supplementary material for "Diel cycle of lanthanide-dependent methylotrophy by TMED127/Methylaequorales bacteria in oligotrophic surface seawater": XoxF_alignment

```
>MAI84053_MED657_o_f_TMED127_g_GCA-2691145
------M-SI-LKKTLIASA----ALALPF---SVMANSKLEKMMGDPKQWVQQSGDYAG
HRYSELDQINKSNVGSLKVAWTFSTGVLRGHEGGPLVI--------GD-VMYVHTAIPNI
VYALDLNDNQRILWKYNPVKGGKYPGGETMDRV-ISVMCCDNVNRGVNYHPNGTIVLHAA
DTSVIALDAKTGKEKWVTTNL-----HAGTGK----------YGVSASTGSGQPFIIKDM
VGVGCSGAEFGVRCNWTQYDIKSGKRLWRAYSMGPDKDMLVDPN--KTTSML--------
---------------------KPIGKDSSLKTW------------------PGDAWKIGG
GSIWGFYGYDSKLNTMYYGTGNPSTWNPVVRPGDNKWSMTINARNPDNGMAKWLYQMTPY
DEWDYDGVNEMILVDMK-VKGKNRKALVHFDRNGFAYTMDRVSGELLVAEKYDPAVNWAT
HVDMKT-----GRPQVVDRYSTAHQGE--DVNTTNICPAALGTKDQQPAAYDRLSGLFMV
PTNHVCMDYEPFKV-----DYVAGNAYVGATLSMYPA--------PGG----THLGNFIG
WDADKGKIAWSNPEPFSVWSGAMATAGGITFYGTLEGYAKAVES-KTGKELFSF-KTPSG
IIGNFMAYQHKGKQYVGILSGVGGWAGI-----GLAAGLTD----PT-----AGLGAVGA
YSALSNHTNL--GGVLTVFALP---------------
>Ga0456454_000930_29_1861_BATS_5m_118_RPKM
------M-SI-LKKTLIASA----ALALPF---SVMANSKLEKMMGDPKQWVQQSGDYAG
HRYSELDQINKSNVGSLKVAWTFSTGVLRGHEGGPLVI--------GD-VMYVHTAIPNI
VYALDLNDNQRILWKYNPVKGGKYPGGETMDRV-ISVMCCDNVNRGVNYHPNGTIVLHAA
DTSVIALDAKTGKEKWVTTNL-----HAGTGK----------YGVSASTGSGQPFIIKDM
VGVGCSGAEFGVRCNWTQYDIKSGKRLWRAYSMGPDKDMLVDPN--KTTSML--------
---------------------KPIGKDSSLKTW------------------PGDAWKIGG
GSIWGFYGYDSKLNTMYYGTGNPSTWNPVVRPGDNKWSMTINARNPDNGMAKWLYQMTPY
DEWDYDGVNEMILVDMK-VKGKNRKALVHFDRNGFAYTMDRVSGELLVAEKYDPAVNWAT
HVDMKT-----GRPQVVDRYSTAHQGE--DVNTTNICPAALGTKDQQPAAYDRLSGLFMV
PTNHVCMDYEPFKV-----DYVAGNAYVGATLSMYPA--------PGG----THLGNFIG
WDADKGKIAWSNPEPFSVWSGAMATAGGITFYGTLEGYAKAVES-KTGKELFSF-KTPSG
IIGNFMAYSHKGKQYVGILSGVGGWAGI-----GLAAGLTD----PT-----AGLGAVGA
YSALSNHTAL--GGVLTVFALP---------------
>AG-893-J11_g_s_GCA-002690875
------M-SI-LKKSIIASA----ALALPL---AVSANSKIERMMKDPSQWVQQSGDYAG
HRYSALDQINKSNVGSLKVAWTFSTGVLRGHEGGPLVI--------GD-TMYVHTAIPNI
VYALDLNDNQRIMWKYNPVKGGKYPGGETMDRV-ISVMCCDNVNRGVNYHPNGTIVLHAA
DTSVIALDAKTGKEKWVTTNL-----HAGTGK----------YGVSASTGSGQPFIIKDM
VGVGCSGAEFGVRCNWTQYDINSGKRLWRAYSMGPDKDMLVDPN--KTTSML--------
---------------------KPIGKDSSLKTW------------------PGDAWKIGG
GSIWGFYGYDSSLNTLYYGTGNPSTWNPVVRPGDNKWSMTINARNPDNGMAKWLYQMTPY
DEWDYDGVNEMILVDMK-VKGKNRQALVHFDRNGFAYTMDRASGELLVAEKYDPAVNWAT
HVDMKT-----GRPQVVDRYSTAHQGE--DVNTTNICPAALGSKDQQPAAYDRLSGLFMV
PTNHVCMDYEPFKV-----DYVAGNAYVGATLSMYPA--------PGG----THLGNFIA
WDADKGKIVWSNPEPFSVWSGALATAGGITFYGTLEGYAKAVES-KTGKELFKF-KTPSG
IIGNFMAYSHKGKQYVGILSGVGGWAGI-----GLAAGLTD----PT-----AGLGAVGA
YSALSNHTAL--GGVLTVFALP---------------
>AG-899-D02_g_s_GCA-002690875
------M-SI-LKKSIIASA----ALALPL---AVSANSKIERMMKDPSQWVQQSGDYAG
HRYSALDQINKSNVGSLKVAWTFSTGVLRGHEGGPLVI--------GD-TMYVHTAIPNI
VYALDLNDNQRIMWKYNPVKGGKYPGGETMDRV-ISVMCCDNVNRGVNYHPNGTIVLHAA
DTSVIALDAKTGKEKWVTTNL-----HAGTGK----------YGVSASTGSGQPFIIKDM
VGVGCSGAEFGVRCNWTQYDINSGKRLWRAYSMGPDKDMLVDPN--KTTSML--------
```

```
---------------------KPIGKDSSLKTW------------------PGDAWKIGG
GSIWGFYGYDSSLNTLYYGTGNPSTWNPVVRPGDNKWSMTINARNPDNGMAKWLYQMTPY
DEWDYDGVNEMILVDMK-VKGKNRQALVHFDRNGFAYTMDRASGELLVAEKYDPAVNWAT
HVDMKT-----GRPQVVDRYSTAHQGE--DVNTTNICPAALGSKDQQPAAYDRLSGLFMV
PTNHVCMDYEPFKV-----DYVAGNAYVGATLSMYPA--------PGG----THLGNFIA
WDADKGKIVWSNPEPFSVWSGALATAGGITFYGTLEGYAKAVES-KTGKELFKF-KTPSG
IIGNFMAYSHKGKQYVGILSGVGGWAGI-----GLAAGLTD----PT-----AGLGAVGA
YSALSNHTAL--GGVLTVFALP---------------
>AG-892-K07_g_s_GCA-002690875
------M-SI-LKKSIIASA----ALALPL---AVSANSKIERMMKDPSQWVQQSGDYAG
HRYSALDQINKSNVGSLKVAWTFSTGVLRGHEGGPLVI--------GD-TMYVHTAIPNI
VYALDLNDNQRIMWKYNPVKGGKYPGGETMDRV-ISVMCCDNVNRGVNYHPNGTIVLHAA
DTSVIALDAKTGKEKWVTTNL-----HAGTGK----------YGVSASTGSGQPFIIKDM
VGVGCSGAEFGVRCNWTQYDINSGKRLWRAYSMGPDKDMLVDPN--KTTSML--------
---------------------KPIGKDSSLKTW------------------PGDAWKIGG
GSIWGFYGYDSSLNTLYYGTGNPSTWNPVVRPGDNKWSMTINARNPDNGMAKWLYQMTPY
DEWDYDGVNEMILVDMK-VKGKNRQALVHFDRNGFAYTMDRASGELLVAEKYDPAVNWAT
HVDMKT-----GRPQVVDRYSTAHQGE--DVNTTNICPAALGSKDQQPAAYDRLSGLFMV
PTNHVCMDYEPFKV-----DYVAGNAYVGATLSMYPA--------PGG----THLGNFIA
WDADKGKIVWSNPEPFSVWSGALATAGGITFYGTLEGYAKAVES-KTGKELFKF-KTPSG
IIGNFMAYSHKGKQYVGILSGVGGWAGI-----GLAAGLTD----PT-----AGLGAVGA
YSALSNHTAL--GGVLTVFALP---------------
>GOS2235_10351912_Colon_Panama
------M-SI-LKKSIIASA----ALALPL---AVSANSKIERMMKDPSQWVQQSGDYAG
HRYSALDQINKSNVGSLKVAWTFSTGVLRGHEGGPLVI--------GD-TMYVHTAIPNI
VYALDLNDNQRIMWKYNPVKGGKYPGGETMDRV-ISVMCCDNVNRGVNYHPNGTIVLHAA
DTSVIALDAKTGKEKWVTTNL-----HAGTGK----------YGVSASTGSGQPFIIKDM
VGVGCSGAEFGVRCNWTQYDINSGKRLWRAYSMGPDKDMLVDPN--KTTSML--------
---------------------KPIGKDSSLKTW------------------PGDAWKIGG
GSIWGFYGYDSSLNTLYYGTGNPSTWNPVVRPGDNKWSMTINARNPDNGMAKWLYQMTPY
DEWDYDGVNEMILVDMK-VKGKNRQALVHFDRNGFAYTMDRASGELLVAEKYDPAVNWAT
HVDMKT-----GRPQVVDRYSTAHQGE--DVNTTNICPAALGSKDQQPAAYDRLSGLFMV
PTNHVCMDYEPFKV-----DYVAGNAYVGATLSMYPA--------PGG----THLGNFIA
WDADKGKIVWSNPEPFSVWSGALATAGGITFYGTLEGYAKAVES-KTGKELFKF-KTPSG
IIGNFMAYSHKGKQYVGILSGVGGWAGI-----GLAAGLTD----PT-----AGLGAVGA
YSALSNHTAL--GGVLTVFALP---------------
>AG-915-P07_g_s_GCA-002690875
------M-SI-LKKSIIASA----ALALPL---AVSANSKIERMMKDPSQWVQQSGDYAG
HRYSALDQINKSNVGSLKVAWTFSTGVLRGHEGGPLVI--------GD-TMYVHTAIPNI
VYALDLNDNQRIMWKYNPVKGGKYPGGETMDRV-ISVMCCDNVNRGVNYHPNGTIVLHAA
DTSVIALDAKTGKEKWVTTNL-----HSGTGK----------YGVSASTGSGQPFIIKDM
VGVGCSGAEFGVRCNWTQYDINSGKRLWRAYSMGPDKDMLVDPN--KTTSML--------
---------------------KPIGKDSSLKTW------------------PGDAWKIGG
GSIWGFYGYDSSLNTLYYGTGNPSTWNPVVRPGDNKWSMTINARNPDNGMAKWLYQMTPY
DEWDYDGVNEMILVDMK-VKGKNRQALVHFDRNGFAYTMDRASGELLVAEKYDPAVNWAT
HVDMKT-----GRPQVVDRYSTAHQGE--DVNTTNICPAALGSKDQQPAAYDRLSGLFMV
PTNHVCMDYEPFKV-----DYVAGNAYVGATLSMYPA--------PGG----THLGNFIA
WDADKGKIVWSNPEPFSVWSGALATAGGITFYGTLEGYAKAVES-KTGKELFKF-KTPSG
```

```
IIGNFMAYSHKGKQYVGILSGVGGWAGI-----GLAAGLTD----PT-----AGLGAVGA
YSALSNHTAL--GGVLTVFALP---------------
>Ga0456549_000444_30_1862_BATS_5m_2056_RPKM
------M-SI-LKKSIIASA----ALALPL---AVSANSKIEKMMKDPSQWVQQSGDYAG
HRYSALDQINKSNVGSLKVAWTFSTGVLRGHEGGPLVI--------GD-TMYVHTAIPNI
VYALDLNDNQRIMWKYNPVKGGKYPGGETMDRV-ISVMCCDNVNRGVNYHPNGTIVLHAA
DTSVIALDAKTGKEKWVTTNL-----HAGTGK----------YGVSASTGSGQPFIIKDM
VGVGCSGAEFGVRCNWTQYDINSGKRLWRAYSMGPDKDMLVDPN--KTTSML--------
---------------------KPIGKDSSLKTW------------------PGDAWKIGG
GSIWGFYGYDSKLNTLYYGTGNPSTWNPVVRPGDNKWSMTIMARNPDNGMAKWLYQMTPY
DEWDYDGVNEMILVDMK-VKGKNRQALVHFDRNGFAYTMDRASGELLVAEKYDPAVNWAT
HVDMKT-----GRPQVVDRYSTAHQGE--DVNTTNICPAALGTKDQQPAAYDRLSGLFMV
PTNHVCMDYEPFKV-----DYVAGNAYVGATLSMYPA--------PGG----THLGNFIA
WDADKGKIAWSNPEPFSVWSGALATAGGITFYGTLEGYAKAVES-KTGKELFSF-KTPSG
IIGNFMAYSHKGKQYVGILSGVGGWAGI-----GLAAGLTD----PT-----AGLGAVGA
YSALSNHTAL--GGVLTVFALP---------------
>Ga0456466_000712_8_1909_5m_BATS_1763_RPKM
------M-SI-LKKSIIASA----ALALPF---AVSANSKLEKMMKDPGQWVQQSGDYAG
HRFSNLDQINKSNVGSLKVAWTFSTGVLRGHEGGPLVI--------GD-VMYVHTAIPNI
VYALDLNDNQRIMWKYNPVKGGKYPGGETMDRV-ISVMCCDNVNRGVNYHPNGTIVLHAA
DTSVIALDAKTGKEKWVTTNL-----HAGTGK----------YGVSASTGSGQPFIIKDM
VGVGCSGAEFGVRCNWTQYDINSGKRLWRAYSMGPDKDMLVDPN--KTTSML--------
---------------------KPIGKDSSLKTW------------------PGDAWKIGG
GSIWGFYGYDSSLNTLYYGTGNPSTWNPVVRPGDNKWSMTINARNPDNGMAKWLYQMTPY
DEWDYDGVNEMILVDMK-VKGKNRQALVHFDRNGFAYTMDRASGELLVAEKYDPAVNWAT
HVDMKT-----GRPQVVDRYSTAHQGE--DVNTTNICPAALGTKDQQPAAYDRLSGLFMV
PTNHVCMDYEPFKV-----DYVAGNAYVGATLSMYPA--------PGG----THLGNFIA
WDADKGKIAWSNPEPFSVWSGALATAGGITFYGTLEGYAKAVES-KTGKELFSF-KTPSG
IIGNFMAYSHKGKQYVGILSGVGGWAGI-----GLAAGLTD----PT-----AGLGAVGA
YSALSNHTAL--GGVLTVFALP---------------
>GIR25668_CM1_5m_P40_g_GCA-2691245
------M-SI-LKKSIIASA----ALALPF---AVSANSKLEKMMKDPGQWVQQSGDYAG
HRFSNLDQINKSNVGSLKVAWTFSTGVLRGHEGGPLVI--------GD-VMYVHTAIPNI
VYALDLNDNQRIMWKYNPVKGGKYPGGETMDRV-ISVMCCDNVNRGVNYHPNGTIVLHAA
DTSVIALDAKTGKEKWITTNL-----HSGTGK----------YGVSASTGSGQPFIIKDM
VGVGCSGAEFGVRCNWTQYDINSGKRLWRAYSMGPDKDMLVDPN--KTTSML--------
---------------------KPIGKDSSLKTW------------------PGDAWKIGG
GSIWGFYGYDSKLNTLYYGTGNPSTWNPVVRPGDNKWSMTIMARNPDNGMAKWLYQMTPY
DEWDYDGVNEMILVDMK-VKGKNRQALVHFDRNGFAYTMDRASGELLVAEKYDPAVNWAT
HVDMKT-----GRPQVVDRYSTAHQGE--DVNTTNICPAALGTKDQQPAAYDRLSGLFMV
PTNHVCMDYEPFKV-----DYVAGNAYVGATLSMYPA--------PGG----THLGNFIA
WDADKGKIAWSNPEPFSVWSGALATAGGITFYGTLEGYAKAVES-KTGKELFSF-KTPSG
IIGNFMAYSHKGKQYVGILSGVGGWAGI-----GLAAGLTD----PT-----AGLGAVGA
YSALSNHTAL--GGVLTVFALP---------------
>AG-892-F10_g_s_GCA-2691245
------M-SI-LKKSIIASA----ALALPF---AVSANSKLEKMMKDPGQWVQQSGDYAG
HRFSNLDQINKSNVGSLKVAWTFSTGVLRGHEGGPLVI--------GD-VMYVHTAIPNI
VYALDLNDNQRIMWKYNPVKGGKYPGGETMDRV-ISVMCCDNVNRGVNYHPNGTIVLHAA
```

```
DTSVIALDAKTGKEKWITTNL-----HSGTGK----------YGVSASTGSGQPFIIKDM
VGVGCSGAEFGVRCNWTQYDINSGKRLWRAYSMGPDKDMLVDPN--KTTSML--------
---------------------KPIGKDSSLKTW------------------PGDAWKIGG
GSIWGFYGYDSKLNTLYYGTGNPSTWNPVVRPGDNKWSMTIMARNPDNGMAKWLYQMTPY
DEWDYDGVNEMILVDMK-VKGKNRQALVHFDRNGFAYTMDRASGELLVAEKYDPAVNWAT
HVDMKT-----GRPQVVDRYSTAHQGE--DVNTTNICPAALGTKDQQPAAYDRLSGLFMV
PTNHVCMDYEPFKV-----DYVAGNAYVGATLSMYPA--------PGG----THLGNFIA
WDADKGKIAWSNPEPFSVWSGALATAGGITFYGTLEGYAKAVES-KTGKELFSF-KTPSG
IIGNFMAYSHKGKQYVGILSGVGGWAGI-----GLAAGLTD----PT-----AGLGAVGA
YSALSNHTAL--GGVLTVFALP---------------
>PDH56613_MED-G10_g_s_MED-G10
------M-SI-LKKSIIASA----ALALPL---AVSANSKLEKMMKDPSQWVQQSGDYAG
HRYSSLDQINKSNVSSLKVAWTFSTGVLRGHEGGPLVI--------GD-TMYVHTAIPNI
VYALDLNDNQRIIWKYNPVKGGKYPGGETMDRV-ISVMCCDNVNRGVNYHPNGTIVLHAA
DTSVIALDAKTGKEKWITTNL-----HAGTGK----------YGVSASTGSGQPFIIKDT
VGVGCSGAEFGVRCNWTSYDIKNGKKLWRAYSMGPDKDMLVDPN--KTTSML--------
---------------------KPIGKDSSLKTW------------------PGDAWKIGG
GSIWGFYGYDSKLNTLYYGTGNPSTWNPVVRPGDNKWSMTINARNPDNGMAKWLYQMTPY
DEWDYDGVNEMILVDMK-VKGKNRQALVHFDRNGFAYTMDRASGELLVAEKYDPAVNWAT
HVDMKT-----GRPQVVDRYSTAHQGE--DVNTTNICPAALGTKDQQPAAYDRLSGLFMV
PTNHVCMDYEPFKV-----DYVAGNAYVGATLSMYPA--------PGG----THLGNFIG
WDADKGKIVWSNPEPFSVWSGAMATAGGITFYGTLEGYAKAVES-KTGKELFSF-KTPSG
IIGNFMSYTHKGKQYVGILSGVGGWAGI-----GLAAGLTD----PT-----AGLGAVGA
YSALADHTAL--GGVLTVFALP---------------
>SD3109_NP_metabat_bins_2.tsv.358_g_s_TMED13
------M-SI-LKKSVIASA----ALALPF---AVSANSKLEKMMKDPNQWVQQSGDYAG
HRYSELDQINKSNVGSLKVAWTFSTGVLRGHEGGPLVI--------GD-TMYVHTAIPNI
VYALDLNDNQRIIWKYNPVKGGKYPGGETMDRV-ISVMCCDNVNRGVNYHPNGTIVLHAA
DTSVIALDAKTGKEKWITTNL-----HAGTGK----------YGVSASTGSGQPFIIKDT
VGVGCSGAEFGVRCNWTSYDIKSGKRLWRAYSMGPDKDMLVDPN--KTTSML--------
---------------------KPIGKDSSLKTW------------------PGDAWKIGG
GSIWGFYGYDSKLNTLYYGTGNPSTWNPVVRPGDNKWSMTINARNPDNGMAKWLYQMTPY
DEWDYDGVNEMILVDMK-VKGKNRQALVHFDRNGFAYTMDRASGELLVAEKYDPAVNWAT
HVDMKT-----GRPQVVDRYSTAHQGE--DVNTTNICPAALGTKDQQPAAYDRMSGLFMV
PTNHVCMDYEPFKV-----DYVAGNAYVGATLSMYPA--------PGG----THLGNFIA
WDADKGKIVWSNPEPFSVWSGAMATAGGITFYGTLEGYAKAVES-KTGKELFSF-KTPSG
IIGNFMSYTHKGKQYVGILSGVGGWAGI-----GLAAGLTD----PT-----AGLGAVGA
YSALSNHTAL--GGVLTVFALP---------------
>MEL0124461_139539_COM_metabat_bins_2.tsv.238_g_s_TMED13
------M-SI-LKKSVIASA----ALALPF---AVSANSKLEKMMKDPNQWVQQSGDYAG
HRYSELDQINKSNVGSLKVAWTFSTGVLRGHEGGPLVI--------GD-TMYVHTAIPNI
VYALDLNDNQRIIWKYNPVKGGKYPGGETMDRV-ISVMCCDNVNRGVNYHPNGTIVLHAA
DTSVIALDAKTGKEKWITTNL-----HSGTGK----------YGVSASTGSGQPFIIKDT
VGVGCSGAEFGVRCNWTSYDIKSGKRLWRAYSMGPDKDMLVDPN--KTTSML--------
---------------------KPIGKDSSLKTW------------------PGDAWKIGG
GSIWGFYGYDSKLNTLYYGTGNPSTWNPVVRPGDNKWSMTINARNPDNGMAKWLYQMTPY
DEWDYDGVNEMILVDMK-VKGKNRQALVHFDRNGFAYTMDRASGELLVAEKYDPAVNWAT
HVDMKT-----GRPQVVDRYSTAHQGE--DVNTTNICPAALGSKDQQPAAYDRLSGLFMV
```

```
PTNHVCMDYEPFKV-----DYVAGNAYVGATLSMYPA--------PGG----THLGNFIA
WDADKGKIVWSNPEPFSVWSGAMATAGGITFYGTLEGYAKAVES-KTGKELFSF-KTPSG
IIGNFMSYTHKGKQYVGILSGVGGWAGI-----GLAAGLTD----PT-----AGLGAVGA
YSALSNHTAL--GGVLTVFALP---------------
>AG-915-O07_g_s_TMED13
------M-SI-LKKSIVASA----ALALPF---AVSANSKLEKMMKDPNQWVQQSGDYAG
HRYSELDQINKSNVGSLKVAWTFSTGVLRGHEGGPLVI--------GD-TMYVHTAIPNI
VYALDLNDNQRIIWKYNPVKGGKYPGGETMDRV-ISVMCCDNVNRGVNYHPNGTIVLHAA
DTSVIALDAKTGKEKWITTNL-----HAGTGK----------YGVSASTGSGQPFIIKDT
VGVGCSGAEFGVRCNWTSYDIKSGKRLWRAYSMGPDKDMLVDPN--KTTSML--------
---------------------KPIGKDSSLKTW------------------PGDSWKIGG
GSIWGFYGYDSKLNTLYYGTGNPSTWNPVVRPGDNKWSMTINARNPDNGMAKWLYQMTPY
DEWDYDGVNEMILVDMK-VKGKNRQALVHFDRNGFAYTMDRASGELLVAEKYDPAVNWAT
HVDMKT-----GRPQVVDRYSTAHQGE--DVNTTNICPAALGTKDQQPAAYDRMSGLFMV
PTNHVCMDYEPFKV-----DYVAGNAYVGATLSMYPA--------PGG----THLGNFIG
WDADKGKIVWSNPEPFSVWSGAMATAGGITFYGTLEGYAKAVES-KTGKELFSF-KTPSG
IIGNFMSYTHKGKQYVGILSGVGGWAGI-----GLAAGLTD----PT-----AGLGAVGA
YSALSSHTAL--GGVLTVFALP---------------
>AG-404-M02_g_s_GCA-2711515
------M-SI-LKKSIIASA----ALALPL---AVSANSKLEKMMKDPSQWVQQSGDYAG
HRYTTLDQINKSNVGSLKVAWTFSTGVLRGHEGGPLVI--------GD-TMYVHTAIPNI
VYALDLNDDQRIIWKYNPVKGGKYPGGETMDRV-ISVMCCDNVNRGVNYHPNGTIVLHAA
DTSVIALDAKTGKEKWITTNL-----HAGTGK----------YGVSASTGSGQPFIIKDT
VGVGCSGAEFGVRCNWTSYDINSGKRLWRAYSMGPDKDMLVDPN--KTTSML--------
---------------------KPIGKDSSLKTW------------------PGDAWKIGG
GSIWGFYGYDSKLNTLYYGTGNPSTWNPVVRPGDNKWSMTINARNPDNGMAKWLYQMTPY
DEWDYDGVNEMILVDMK-VKGKNRQALVHFDRNGFAYTMDRASGELLVAEKYDPAVNWAT
HVDMKT-----GRPQVQDRYSTAHQGE--DVNTTNICPAALGSKDQQPAAYDRLSGLFMV
PTNHVCMDYEPFKV-----DYVAGNAYVGATLSMYPA--------PGG----THLGNMIA
WDADKGKIVWSNPEPFSVWSGAMATAGGITFYGTLEGYAKAVES-KTGKELFKF-KTPSG
IIGNFMSYKHKGKQYVGILSGVGGWAGI-----GLAAGLTD----PT-----AGLGAVGA
YSALHNHTAL--GGVLTVFSLP---------------
>AG-337-B03_g_s_SP4073
------M-SI-LKKTLIASA----ALALPF---SVSANSKLEKMMKDSSQWVQQSGNYAG
ERYTELDQINKSNVGSMKVAWTFSTGVLRGHEGGPLVI--------GD-VMYVHTAIPNI
VYALDLNDNQRILWKYNPVKGGKYPGGETMDRV-ISVMCCDNVNRGVNYHPNGTIVLHAA
DTSVIALDAKTGKEKWITTNL-----HAGTGK----------YGVSASTGSGQPFIIKDM
VGVGCSGAEFGVRCNWTQYDIKSGKRLWRAYSMGPDKDMLVDPA--KTTSML--------
---------------------KPIGKDSSLKTW------------------PGDAWKIGG
GSIWGFYGYDSKLNTLYYGTGNPSTWNPVVRPGDNKWSMTINARNPDNGMAKWLYQMTPY
DEWDYDGVNEMILVDMK-VKGKNRKALVHFDRNGFAYTMDRASGELLVAEKYDPAVNWAT
HVDMKT-----GRPQVVDKYSTAHQGE--DVNTTNICPAALGTKDQQPASYDRLSGLFMV
PTNHVCMDYEPFKV-----DYVAGNAYVGATLSMYPA--------PGG----THLGNFIG
WDADKGEIAWSNPEPFSVWSGALSTAGGITFYGTLEGYAKAVES-KTGKELFSF-KTPSG
IIGNFMAYEHKGKQYVGILSGVGGWAGI-----GLAAGLTD----PT-----AGLGAVGA
YAALSNHTNL--GGVLTVFALP---------------
>AG-337-E20_g_s_SP4073
------M-SI-LKKTLIASA----ALALPF---SVSANSKLEKMMKDSSQWVQQSGNYAG
```

```
ERYTELDQINKSNVGSMKVAWTFSTGVLRGHEGGPLVI--------GD-VMYVHTAIPNI
VYALDLNDNQRILWKYNPVKGGKYPGGETMDRV-ISVMCCDNVNRGVNYHPNGTIVLHAA
DTSVIALDAKTGKEKWITTNL-----HAGTGK----------YGVSASTGSGQPFIIKDM
VGVGCSGAEFGVRCNWTQYDIKSGKRLWRAYSMGPDKDMLVDPA--KTTSML--------
---------------------KPIGKDSSLKTW------------------PGDAWKIGG
GSIWGFYGYDSKLNTLYYGTGNPSTWNPVVRPGDNKWSMTINARNPDNGMAKWLYQMTPY
DEWDYDGVNEMILVDMK-VKGKNRKALVHFDRNGFAYTMDRASGELLVAEKYDPAVNWAT
HVDMKT-----GRPQVVDKYSTAHQGE--DVNTTNICPAALGTKDQQPASYDRLSGLFMV
PTNHVCMDYEPFKV-----DYVAGNAYVGATLSMYPA--------PGG----THLGNFIA
WDADKGEIVWSNPEPFSVWSGALSTAGGITFYGTLEGYAKAVES-KTGKELFSF-KTPSG
IIGNFMAYEHKGKQYVGILSGVGGWAGI-----GLAAGLTD----PT-----AGLGAVGA
YAALSNHTNL--GGVLTVFALP---------------
>MDC3176690_AH-716-D07_g_s_CACBWF01
------M-SI-LKKTLIASA----ALALPG---LVSANAKLEKMMKDPSQWVQQSGDYAG
HRYSELDQINKSNVGSLKVAWTFSTGVLRGHEGGPLVI--------GD-VMYVHTAIPNI
VYALDLNDNQRIMWKYNPVKGGKYPGGETMDRV-ISVMCCDNVNRGVNYHPNGTIVLHAA
DTSVIALDAKTGKEKWVTTNL-----HAGTGK----------YGVSASTGSGQPFIIKDM
VGVGCSGAEFGVRCNWTQYDIKTGKRLWRAYSMGPDADMLVDPS--KTTSML--------
---------------------KPVGKDSSLKTW------------------PGDAWKIGG
GSIWGFYGYDSKLNTLYYGTGNPSTWNPVVRPGDNKWSMTINARNPDNGMAKWLYQMTPY
DEWDYDGVNEMILVDMK-VKGKNRKTLVHFDRNGFAYTMDRASGELLVAEKYDPAVNWAT
HVDMKT-----GRPQVVDRYSTAHQGE--DVNTTNICPAALGTKDQQPAAYDRLSGLFMV
PTNHVCMDYEPFKV-----DYVAGNAYVGATLSMYPA--------PGG----THLGNFIA
WDADKGKIVWSNPEPFSVWSGAMATAGGITFYGTLEGYAKAVDS-KTGKELFSF-KTPSG
IIGNFMAYEHKGKQYVGILSGVGGWAGI-----GLAAGLTD----PT-----AGLGAVGA
YSALHNHTNL--GGVLTVFALP---------------
>AG-896-A16_g_s_CACBWF01
------M-SI-LKKTLIASA----ALALPG---LVSANAKLEKMMKDPSQWVQQSGDYAG
HRYSELDQINKSNVGSLKVAWTFSTGVLRGHEGGPLVI--------GD-VMYVHTAIPNI
VYALDLNDNQRIMWKYNPVKGGKYPGGETMDRV-ISVMCCDNVNRGVNYHPNGTIVLHAA
DTSVIALDAKTGKEKWVTTNL-----HSGTGK----------YGVSASTGSGQPFIIKDM
VGVGCSGAEFGVRCNWTQYDIKTGKRLWRAYSMGPDADMLVDPS--KTTSML--------
---------------------KPVGKDSSLKTW------------------PGDAWKIGG
GSIWGFYGYDSKLNTLYYGTGNPSTWNPVVRPGDNKWSMTINARNPDNGMAKWLYQMTPY
DEWDYDGVNEMILVDMK-VKGKNRKALVHFDRNGFAYTMDRASGELLVAEKYDPAVNWAT
HVDMKT-----GRPQVVDRYSTAHQGE--DVNTTNICPAALGTKDQQPAAYDRLSGLFMV
PTNHVCMDYEPFKV-----DYVAGNAYVGATLSMYPA--------PGG----THLGNFIA
WDADKGKIVWSNPEPFSVWSGAMATAGGITFYGTLEGYAKAVDS-KTGKELFSF-KTPSG
IIGNFMAYEHKGKQYVGILSGVGGWAGI-----GLAAGLTD----PT-----AGLGAVGA
YSALHNHTNL--GGVLTVFALP---------------
>Ga0394919_003329_118_1950_IndianOcean_Subantarctic_50m_233_RPKM
------M-SI-LKKTLIASA----ALALPF---SVSANAKLEKMMSDPNQWVQQSGDYAG
HRYSELDSINKSNVGSLKVAWSFSTGVLRGHEGSPLVI--------GD-VMYVHTAIPNI
VYALDLNDDQKILWKYNPVKGGKYPGGETMDRV-ISVMCCDNVNRGVNYHPNGTIVLHAA
DTSVIALDAKTGKEKWVTTNL-----HSGTGK----------YGVSASTGSGQPFIIKDM
VGVGCSGAEFGVRCNWTQYDIKSGKRLWRAYSMGPDKDMLVDPS--KTTSML--------
---------------------KPIGKDSSLKTW------------------PGDAWKIGG
GSIWGFYGYDSKLNTMYYGTGNPSTWNPVVRPGDNKWSMTIMARNPDNGMAKWLFQMTPY
```

```
DEWDYDGVNEMILVDMK-VKGKNRKALVHFDRNGFAYTMDRASGELLVAEKYDPAVNWAT
HVDYKT-----GRPQVVDQYSTAHQGE--DVNSTNICPAALGTKDQQPASYDRLSGLFMV
PTNHVCMDYEPFKV-----DYVAGNAYVGATLSMYPA--------PGG----THLGAFIG
WDADKGKIAWSNPEPFSVWSGAMSTAGGVTFYGTLEGYAKAVES-KTGKELFKF-KTPSG
IIGNFMSYSHKGKQYVGILSGVGGWAGI-----GLAAGLTD----PS-----AGLGAVGA
YAALSNHTNL--GGVLTVFSLP---------------
>HAE75578_UBA8558_o_UBA8558
------MKSV-LKAAAVVTA----LVVGAG---IASANDELKKLSADENNWAMPTKDYAA
TRYSKLDQINKGNVGNMQVAWTFSTGVLRGHEGGPLVI--------GDGFLYIHTAIPNK
VFKLDLNNNAAIAWAYNPVKGGKYPSGQAMDRV-ISVMCCDNVNRGVAY-ADGKIFLHAA
DTSLIALDKNSGKEVWVVENM-----HAGKGK----------YAVSASTGTGVPFIVKNL
VMVGCSGAEFGVRCNWTAYNLKNGKRVWRAYSMGSDKDIIADPK--KTTHML--------
---------------------KPVGKNSSTKTW------------------TKDQWKTGG
GSIWGWYAWDESLDLIYYGTGNPSTWNPVVRPGDNKWSMTIMARDPDNGKAAWFYQMTPY
DEWDYDGVNEMILADV-----AGRKALVHFDRNGFAYTMDRATGELLVAEFYDPASNWAT
HVDMKT-----GRPQVIPAKSTATQGE--NVNTKDICPAALGSKDQQPAAYSPLTGLFYV
PTNHVCMDYEPFKV-----DYVAGNAYVGATLSMFPA--------PGG----THLGAFIA
WDAAKGKIVWSNAEPFSVWSGVLTTAGGIACVGTLEGYLKCVDQ-KDGKELYNF-KTPSG
IIGNVNTWAHNGKQYIGVLSGVGGWAGI-----GLAAGLTD----PT-----AGLGAVGA
YAALSNFTQL--GGVLTVFHLP---------------
>Ga0593109_002684_64_2031_BATS_2000m_136_RPKM
------MKSV-LKAAAVVTA----LVVGAG---IASANDELKKLSADENNWAMPTKDYAA
TRYSKLDQINKGNVGNMQVAWTFSTGVLRGHEGGPLVI--------GDGFLYIHTAIPNK
VFKLDLNNNAAIAWAYNPVKGGKYPSGIAMDRV-ISVMCCDNVNRGVAY-ADGKIFLHAA
DTSLIALDKNNGKEVWVVENM-----HAGKGK----------YAVSASTGTGVPFIVKNM
VMVGCSGAEFGVRCNWTAYNLKNGKRVWRAYSMGSDKDIIANPN--KTTHML--------
---------------------KPVGKNSSTKTW------------------TKDQWKTGG
GSIWGWYAWDESLDLIYYGTGNPSTWNPVVRPGDNKWSMTIMARDPDNGMAAWFYQMTPY
DEWDYDGVNEMILADV-----AGRKALVHFDRNGFAYTMDRATGELLVAEFYDPASNWAT
HVDMKT-----GRPQVIPAKSTATQGE--NVNTKDICPAALGSKDQQPAAYSPLTGLFYV
PTNHVCMDYEPFKV-----DYVAGNAYVGATLSMFPA--------PGG----THLGAFIA
WDAAKGKIVWSNAEPFSVWSGVLTTAGGIACVGTLEGYLKCVDQ-KDGKELYNF-KTPSG
IIGNVNTWAHNGKQYIGVLSGVGGWAGI-----GLAAGLTD----PT-----AGLGAVGA
YAALSNFTQL--GGVLTVFHLP---------------
>MEC8875952_Alphaproteobacteria_XB_MAG762_o__UBA8558
------MKSV-LKAAAVVSA----LVVGAG---VANANDELKRLIADENNWATPTKDYAA
TRYSKLDQINKGNVGNMQVAWTFSTGVLRGHEGGPLVI--------GDGFLYIHTAIPNK
VFKLDLNNNAAIAWAYNPVKGGKYPSGQAMDRV-ISVMCCDNVNRGVAY-ADGKIFLHAA
DTSLIALDKNSGKEVWVVENM-----HAGKGK----------YAVSASTGTGVPFIVKNM
VMVGCSGAEFGVRCNWTAYNLKNGKRVWRAYSMGSDKDIIANPN--KTTHML--------
---------------------KPVGKNSSTKTW------------------TKDQWKTGG
GSIWGWYAWDESLDLIYYGTGNPSTWNPVVRPGDNKWSMTIMARDPDNGMAAWFYQMTPY
DEWDYDGVNEMILADV-----AGRKALVHFDRNGFAYTMDRATGELLVAEFYDPASNWAT
HVDMKT-----GRPQVIPAKSTATQGE--NVNTKDICPAALGSKDQQPAAYSPLTGLFYV
PTNHVCMDYEPFKV-----DYVAGNAYVGATLSMFPA--------PGG----THLGAFIA
WDAAKGKIVWSNDEPFSVWSGVLTTAGGIACVGTLEGYLKCVDQ-KDGKELYKF-KTPSG
IIGNVNTWAHNGKQYIGVLSGVGGWAGI-----GLAAGLTD----PT-----AGLGAVGA
YAALSNFTQL--GGVLTVFHLP---------------
```

```
>MDP6304722_Alphaproteobacteria_ETNP15_MAG_21_o_UBA11136
------MKSI-FKTTLLASA----VFALPG---LAQANDELLRLSADPANWAIPSGNYSG
TRYSELDQVNKSNVGSLKVAWQFSTGVLRGHEGGPLVI--------GD-VMYVVTPIPNI
VFALDLNDEGRIIWKHIPVQGGKYPSGETQARV-TSVMCCDNVIRGVSY-AGGKIFVGLA
DSTLQALDAKTGKLVWQAHNM-----SAGKAGESEATGIKPVRGNAASTNTSVPFVVKDN
VFIGCSGAEFGVRCNMTGYNINSGKRAWRAYSMGPDGDILVDPN--KTTHML--------
---------------------KPVGKNSSLKTW------------------KGDAWQIGG
GSLWGFWAWDPDLDLMYYGTGNPSTWNPVARPGDNRWSMTIMARDPDDGMAKWLYQMTPH
DEWDYDGINEMILVDQE-VKGKNRKALVHFDRNGFNYTLDRATGELLVAEKHDPMVNWAT
HIDMKT-----GRPALVDKYGTEHGGKGLDYLTKGICPAALGTKDQQPAAYSPRTKLFYI
PANHVCMNYEPVHV-----EYVVGDAYVGANLDMMPG--------PGG-----NMGEFVA
WDAGKGKVVWEIAEPFSVWSGVLATAGGIVMYGTLEGFVKAVDE-TSGKLLYKF-NSPSG
IIGNINTWSHKGKQQIGVYSGVGGWAGY-----GVAGKLNDIAGNDR-----AALGAIGG
YRHLKNYTQA--GGVLTVFELP---------------
>Ga0567547_013154_140_2083_Gulf_of_California_150_RPKM
------MKSI-LKTTLLASA----VFALPG---LAQANDELLRLSADPANWAIPSGNYSG
TRYSELDQVNKSNVGSLKVAWQFSTGVLRGHEGGPLVI--------GD-VMYVVTPIPNI
VFALDLNDEGRIIWKHIPVQGGKYPSGETQARV-TSVMCCDNVIRGVSY-AGGKIFVGLA
DSTLQALDAKTGKLVWQAHNM-----SAGKAGESEATGIKPVRGNAASTNTSVPFVVKDN
VFIGCSGAEFGVRCNMTGYNIKSGKRAWRAYSMGPDSDILVDPS--KTTHML--------
---------------------KPVGKNSSLKTW------------------KGDAWQIGG
GSLWGFWAWDPDLDLMYYGTGNPSTWNPVARPGDNRWSMTIMARDPDDGMAKWLYQMTPH
DEWDYDGINEMILVDQE-VKGKNRKALVHFDRNGFNYTLDRATGELLVAEKHDPMVNWAT
HIDMKT-----GRPALVDKYGTEHGGKGLDYLTKGICPAALGTKDQQPAAYSPRTKLFYI
PANHVCMNYEPVHV-----EYVVGDAYVGANLDMMPG--------PGG-----NMGEFVA
WDAGKGKVVWEIAEPFSVWSGVLATAGGIVMYGTLEGFVKAVDE-TSGKLLYKF-NSPSG
IIGNINTWSHKGKQQIGVYSGVGGWAGY-----GVAGKLNDIAGNDR-----AALGAIGG
YRHLKNYTQA--GGVLTVFELP---------------
>MDP6486191_Alphaproteobacteria_Arabian12_MAG_22_ARS27_o_UBA11136
------MKSI-LKTTLLASA----VFALPG---LAQANDELLRLSADPANWAIPSGNYSG
TRYSELDQVNKSNVGSLKVAWQFSTGVLRGHEGGPLVI--------GD-VMYIVTPIPNY
VFALDLNNEGRIIWKHIPVQGGKYPSGETQARV-TSVMCCDNVIRGVSY-AGGKIFVGLA
DSTLQALDAKTGKLVWQAHNM-----SAGKAGESEATGIKPVRGNAASTNTSVPFVVKNN
VFIGCSGAEFGVRCNMTGYNINSGKRAWRAYSMGPDGDILVDPN--KTTHML--------
---------------------KPVGKNSSLKTW------------------KGDAWQIGG
GSLWGFWAWDPDLDLMYYGTGNPSTWNPVARPGDNRWSMTIMARDPDDGMAKWLYQMTPH
DEWDYDGINEMILVDQE-VKGKNRKALVHFDRNGFNYTLDRVTGELLVAEKHDPMVNWAT
HIDMKT-----GRPALVDKYGTEHGGKGLDYLTKGICPAALGTKDQQPAAYSPRTKLFYI
PANHVCMNYEPVHV-----EYVVGDAYVGANLDMMPG--------PGG-----NMGEFVA
WDAGTGKVVWEIAEPFSVWSGVLATAGGIVMYGTLEGFVKAVDE-TSGKELYRF-NSPSG
IIGNINTWLHKGKQQIGVFSGVGGWAGY-----GVAGKLNDIEGNDR-----AALGAIGG
YRHLKNYTQA--GGVLTVFELP---------------
>OUW02738_TMED156_o__Burkholderiales_f_Burkholderiaceae_g_UBA7377
------MKFK-LN-AICAAV--AVTVAAPA---VVHANSSLGKLMNNPKNWAAQSGDFFN
QRHTKLKQITKSNVNKLQMAWSFSTGVLRGHEGGPLVI--------GD-TLYVHSAFPNK
VFAINLADQ-TIKWKYEPKQD---------PSV-IPVMCCDTVNRGLAY-AEGKIFLQQA
DTTLVALNAKSGKVIWSQKNG-----NPKVGS----------------TNTNAPHVFKDK
VFTGISGGEFGVQGFVAAYDINSGKRQWKAYSVGPDDQIMFDPK--RTMAWNN-------
```

```
----------------GK--MRPVGKNSSTKTW------------------QGDQWKIGG
GTTWGWYSYDPKLNLMYYGSGNPSTWNPKQRPGDNKWSMTIFARDVDTGVAKWAYQMTPH
DEWDYDGINEVPLFDGKDKNGKKRGMLAHFDRNGFAYTLDRKTGELLVAEKFDPAVNWAT
HVDMNS-----GRPQVLAKYSTHIQGE--DVNSKGICPAALGSKDQQPVSFDPATGYFLV
PTNHVCMDYEPFRV-----EYTAGQPYVGATLSMFPA---------PNS---HGGMGNVIA
WNPIQGKIEWSIPEKFSVWAGTMTTATGVGFYATLDARLKAVDV-KSGKVLWTSPKLPSG
SIGNVHSWEHRGKQYIGILTGIGGWAGI-----GLAAGLEK----DT-----DGLGAVGG
YRELNRYTDL--GGTMMVFALPN--------------
>OUV31211_TMED100_o__Burkholderiales_f_Burkholderiaceae_g_UBA7377
------MKFK-LN-AICAAV--AVAVSAPA---VVHANSSLEKLMNNPKNWAAQSGDFFN
QRHTKLKQITKSNVNKLQMAWSFSTGVLRGHEGGPLVI--------GD-TLYVHSAFPNK
VFAINLADQ-TIKWKYEPKQD---------PSV-IPVMCCDTVNRGLAY-AEGKIFLQQA
DTTLVALNAKSGKVLWSTKNG-----NPKVGS----------------TNTNAPHIFKNK
VITGISGGEFGVQGFVAAYDINSGKRKWKAYSVGPDDQLKFDPK--KTMAWNN-------
----------------GK--MRPVGKDSSLKTW------------------QGDQWKIGG
GTTWGWYSYDPKLNLMYYGSGNPSTWNPKQRPGDNKWSMTIFARDVDTGVAKWAYQMTPH
DEWDYDGINEVPLFDGKGPKGKKRGMLAHFDRNGFAYTLDRKTGELLVAEKFDPAVNWAT
HVDMKS-----GRPQVLAKYSTHIQGE--DVNSKGICPAALGSKDQQPVSFDPATGLFLV
PTNHVCMDYEPFRV-----EYTAGQPYVGATLSMFPA---------PNS---HGGMGNVIA
WDPIKGKIEWSIPEKFSVWAGTMTTATGVGFYATLDAKLKAVDV-KSGKVLWTSPKLPSG
SIGNVHSWEHRGKQYIGILTGIGGWAGI-----GLAAGLEK----DT-----DGLGAVGG
YRELNKYTEL--GGTMMVFALPN--------------
>OUV02704_TMED82_o__Burkholderiales_f_Burkholderiaceae_g_UBA7377_Ga0456468_000675_27_1871_BATS_5m_478_RPKM
------MNFK-LN-ALCAAV--AVTVAAPA---VVHANSSLGKLMKNPANWAAPSGDFFN
QRHTKLKQITKSNVNKLQMAWSFSTGVLRGHEGGPLVI--------GD-TLYVHSAFPNK
VFAINLADQ-TIKWKYEPKQD---------PSV-IPVMCCDTVNRGLAY-AEGKIFLQQA
DTTLVALNAKNGKLIWSTKNG-----DPKLGQ----------------TNTNAPHIFKDK
VITGISGGEFGVQGFVAAYDINSGKKVWKAYSVGPDSDLKFDPK--KTMAWNN-------
----------------GK--MRPVGKDSSLKTW------------------QGDQWKIGG
GTTWGWYSYDPKLNLMYYGSGNPSTWNPKQRPGDNKWSMTIFARDVDTGVAAWAYQMTPH
DEWDYDGINEVPLFDGKDKNGKKRGMLAHFDRNGFGYTLDRKTGELLVAEKFDPAVNWAT
HVDMKS-----GRPQVLAKYSTHIQGE--DVNTKGICPAALGSKDQQPVSFDPATGYFLV
PTNHVCMDYEPFRV-----EYTAGQPYVGATLSMFPA---------PNS---HGGMGNVIA
WNPVQGKIEWSIPEKFSVWAGTMTTATGVGFYATLDARLKAVDI-KSGKVLWTSPKLPSG
SIGNVHSWEHRGKQYIGILTGIGGWAGI-----GLAAGLEK----DT-----DGLGAVGG
YRELSKYTDL--GGTMMVFALPN--------------
>GIR48069_CM1_5m_P58_o__Burkholderiales_f__Burkholderiaceae_g_UBA7377
------MKFK-LN-ALCAAV--AVSVAAPA---VVHANSSLNKLMNNSSNWAAQSGDFFN
QRHTKLKQINKSNVNKLQMAWSFSTGVLRGHEGGPLVI--------GD-TLYVHSAFPNK
VFAINLADQ-TIKWKYEPKQD---------PSV-IPVMCCDTVNRGLAY-AEGKIFLQQA
DTTLVALNAKNGKVMWSTKNG-----NPKVGQ----------------TNTNAPHVFKDK
VITGISGGEFGVQGFVAAYDINSGKKVWKAYSVGPDDQLMFDPK--KTMAWNN-------
----------------GK--MQPVGKNSSLKTW------------------EGDQWKIGG
GTTWGWYSYDPKNNMMYYGSGNPSTWNPKQRPGDNKWSMTIFARDVDTGMAKWAYQMTPH
DEWDYDGINEVPLFDGKDKNGKSRGMLAHFDRNGFAYTMDRNTGELLVAEKFDPAVNWAT
HVDMKS-----GRPQVLAKYSTHIQGE--DVNTKGICPAALGSKDQQPVSFDPATGYFLV
```

```
PTNHVCMDYEPFRV-----EYTAGQPYVGATLSMFPA--------PNS---HGGMGNVIA
WNPTQGKIEWSIPEKFSVWAGTLTTATGVGFYATLDARLKAVDV-KSGKILWTSPKLPSG
SIGNVHSWEHRGKQYVGILTGIGGWAGI-----GLAAGLEK----DT-----DGLGAVGG
YRELSKYTDL--GGTLMVFALPN--------------
>WP_024300827_Methyloversatilis_discipulorum_FAM1_c__Gammaproteobacteria_o__Burkholderiales_f__Rhodocyclaceae_characterized
------MSSK-MKQLPLWVA----ALAIPG---TALANADVAKLIKDPKNWAMQAGNFEN
QRYSTLNQINKSNVKNMKVAWTFSTGVLRGHEGGPLVI--------GD-TLFVHSPFPNK
VFAIDLETQ-KIKWKYEPKQD---------PSV-IPVMCCDTVNRGLAY-AEGKVFLQQA
DTMLVALDAKSGKKLWEVKNG-----DPKVGG----------------TNTNAPHVFKDK
VITGISGGEFGIRGYVAAYDINTGKQVWKGYSTGPDAEMLMDPE--KTMTWTN-------
----------------GK--MAPVGKDSSLKTW------------------QGDQWKIGG
GTTWGWYSYDPQENLMYYGSGNPSTWNPSQRPGDNKWSMTLWARDVDTGKARWVYQMTPH
DEWDFDGINEVILADVD-VKGKKRKVAVHFDRNGFAYTMDRVSGELLVAEKYDPKVNWAT
HVDMNT-----GRPQVVAKYSTRQNGE--DVNTKGVCPAALGTKDQQPAAFSPKNGLFYV
PTNHVCMDYEPFRV-----EYTAGQPYVGATLSMFPA--------PGS---HGGMGNFIT
WDAGKGKIVQSKPEKFSVWSGALVTAGGIACYGTLEGYLKCVDADDINKELFKF-KTPSG
IIGNVNTWEYKGKQYIGILSGIGGWAGI-----GLAAGLEK----PT-----DGLGAVGG
YAELASYTEL--GGTMTVFALP---------------
>WP_014751259_Advenella_kashmirensis_c__Gammaproteobacteria_o__Burkholderiales_f_Burkholderiaceae_characterized
----M-SFKITLKQMLCVAG----ALAITG---TAQADPQLQEAMKNPDNWAVQAGDFAN
QRYSKLDQINKDNVKNLQVAWTFSTGVLRGHEGGPLVI--------GD-VMYVHSPFPNK
VYAISLKDQ-SLLWKYEPTQD---------PNV-ITIMCCDTVNRGLAF-GDGKIILQQA
DTTMVALDAKTGKEVWKVKNG-----DFKVGE----------------SNTNAPHIFKDK
VITGISGGEFGVRGRLIAYNLKDGKEAWKAYSTGPDNEMLFDPE--KTMTWTD-------
----------------GK--MAAVGKDSSLKSW------------------KGDQWKIGG
GTTWGWFSYDPALNLVYYGTGNPGTWNPSQRPGDNKWSMSIIARDLDTGVAKWVYQMTPH
DEWDYDGVNENILADIK-VGGKDRKALVHFDRNGFGYTLDRESGELLVAEKYDPTINWAD
KIDMKS-----GRPVVNSKYSPAKGGE--DVNVKGICPSALGTKDQQPAAFSPKTGVFYV
PTNHVCMDYEPFAV-----DYVAGQPYVGATVSMFPT--------PNS---HGGMGNFIA
WDADKGKILWSIPERFSVWSGALATAGDVVFYGTLEGYIKAIDT--SGKELWKF-KTPSG
VIGNVTTWKYEGKQYIGVLSGIGGWAGI-----GLAAGLEK----ST-----DGLGAVGG
YKDLAKYTEL--GGALMVFALPDNVATAAAEKTTKTQ
>MexAM1_META1p1740_XoxF5_Me1_characterized
------MRAV---HLLALGA----GLAAAS---PALANESVLKGVANPAEQVLQTVDYAN
TRYSKLDQINASNVKNLQVAWTFSTGVLRGHEGSPLVV--------GN-IMYVHTPFPNI
VYALDLDQGAKIVWKYEPKQD---------PSV-IPVMCCDTVNRGLAY-ADGAILLHQA
DTTLVSLDAKSGKVNWSVKNG-----DPSKGE----------------TNTATVLPVKDK
VIVGISGGEFGVQCHVTAYDLKSGKKVWRGYSIGPDDQLIVDPE--KTTSLG--------
---------------------KPIGKDSSLKTW------------------EGDQWKTGG
GCTWGWFSYDPKLDLMYYGSGNPSTWNPKQRPGDNKWSMTIWARNPDTGMAKWVYQMTPH
DEWDFDGINEMILTDQK-FDGKDRPLLTHFDRNGFGYTLDRATGEVLVAEKFDPVVNWAT
KVDLDKGSKTYGRPLVVSKYSTEQNGE--DVNSKGICPAALGTKDQQPAAFSPKTGLFYV
PTNHVCMDYEPFRV-----TYTPGQPYVGATLSMYPA--------PGS---HGGMGNFIA
WDNLQGKIKWSNPEQFSAWGGALATAGDVVFYGTLEGFLKAVDS-KTGKELYKF-KTPSG
```

```
IIGNVMTYEHKGKQHVAVLSGVGGWAGI-----GLAAGLTD----PN-----AGLGAVGG
YAALSSYTNL--GGQLTVFSLPNN-------------
>MexAM1_META1p2757_XoxF5_Me2_characterized
------MRQA---HLLGLML----ALGTTG----ALANEDVLKRTQDPNQQVLQTLDYAN
TRYSKLDQINASNVKNLQVAWTFSTGVLRGHEGSPLVV--------GD-IMYVHTPFPNI
VYALDLNNDSKILWKYEPKQD---------PSV-IPVMCCDTVNRGLAY-ADGAIILHQA
DTTLVSLDAKTGKVNWSVKNG-----DSKVGE----------------TNTATVLPVKDK
IIVGISGAEYGIRGHMTAYDAKTGKRVWRAYSVGPDDEMLVDPE--KTTSLG--------
---------------------KPIGKDSSLKTW------------------EGDQWKTGG
GATWGWYSYDPKLDLFYYGTANPSTWNPKQRPGDNKWTMAIFARNPDTGQAKWIYQMTPH
DEWDYDGINEMILTDQK-VDGKERPLLTHFDRNGFAYTLDRANGEVLVAEKFDPVVNWAS
KVDLDKGSKNYGRPLVVSKYSTDQNGE--DVNSKGICPAALGTKDQQPAAFSPKTQLFYV
PTNHVCMDYEPFKV-----TYTPGQPYVGATLSMYPA--------PGG---HGGMGNFIA
WDNISGKIKWSNPEQFSVWSGALATAGDVVFYGTLEGYLKAVDS-KTGKELYKF-KTASG
VIGNVMTYTHKGKQYVGVLSGVGGWAGI-----GLAAGLTD----PN-----AGLGAVGG
YAALSQYTNL--GGQLTVFALPN--------------
>WP_010975084_Sinorhizobium_meliloti_5A14_c__Alphaproteobacteria_o__Rhizobiales_f__Rhizobiaceae_characterized
-----------MKRLLTMLA----IMSIGGGAQVAFANDELQKLIDDPNQWAIQTGDYAN
LRYSKLDQINKDNVGKLQVAWTFSTGVLRGHEGSPLVI--------GD-LMYVHTPFPNT
VYALDLSKDGQIVWKYEPKQD---------PNV-IPVMCCDTVNRGVAY-ADNKIFLHQA
DTTVVALDAKTGKVIWSVKNG-----DATKGE----------------TNTATVMPVKDK
ILVGISGGEFGVRGHVTAYSMADGKVLWRGYSMGPDSDTLIDPE--KTTHLG--------
---------------------KPVGKDSGLTTW------------------EGDQWKIGG
GTTWGWYSYDPEENLVYYGTGNPSTWNPTQRPGDNRWSMTIFARDVDTGMAKWLYQMTPH
DEWDYDGVNEMILTEQQ-IDGKDRKLLTHFDRNGFGYTMDRVTGELLVAEKYDPTVNWAT
EVVMDPKSDKYGRPQVVAQYSTEQNGE--DTNTTGVCPAALGTKDQQPAAYSPKTELFYV
PTNHVCMDYEPFRV-----SYTAGQPYVGATLSMYPP--------KDS---HGGMGNFIA
WDNKEGKIKWSLPEPFSVWSGALATAGDVVFYGTLEGYLKAVDA-ATGKELYRF-KTPSG
VIGNVMTYAREGKQYVAVLSGVGGWAGI-----GLAAGLTN----PT-----EGLGAVGG
YSALSNYTAL--GGTLTVFKLPE--------------
>WP_011088953_Bradyrhizobium_diazoefficiens_USDA110_c__Alphaproteobacteria_o__Rhizobiales_f__Xanthobacteraceae_characterized
------MRKVLLATYLGSAA----ALAV-G---SASANDELIKMSQNPKDWVMPAGDYAN
TRYSKLNQINAQNVGKLQVAWTFSTGVLRGHEGGPLII--------GN-MMYVHTPFPNK
VYAIDLSNENKIVWKYEPKQD---------PNV-IPVMCCDTVNRGLSY-GDGKIILHQA
DTNLVALDAKTGQVAWSATNG-----DPSKGQ----------------TGTSAALVVKDK
VLVGISGGEFGVQCHVTAYDLKSGKQVWRAFSEGPDDQIKVDPA--KTTSLG--------
---------------------KPVGADSSLKTW------------------QGDQWKIGG
GCTWGWMSYDPALNLVYYGSGNPSTWNPKQRPGDNKWSMTIFARDADTGMAKWVYQMTPH
DEWDYDGVNEMILSDQQ-INGQARKLLTHFDRNGLGYTMDRESGELLVAEKYDPKVNWTS
GVDMDKNSPTYGRPKVLDAASTDKAGE--DHNVKGICPAALGTKDEQPAAYSPDTQLFYV
PTNHVCMDYEPFKV-----SYTAGQPYVGATLSMYP---------PQG---ESHMGNFIA
WDGKTGKIVWSNKEQFSVWSGALATAGGVVFYGTLEGYLKAVDA-KSGKELYKF-KTPSG
IIGNVTTYENGGKQYVAVLSGVGGWAGI-----GLAAGLTD----PT-----AGLGAVGG
YAALSNYTAL--GGTLTVFSLPAN-------------
```

```
>
WP_002539484_Grimontia_marina_CECT8713_c__Gammaproteobacteria_o__Enterobacterales_f__Vibrionaceae_g__Enterovibrio_characterized
----M-IGAVRLRRCLAAVATLSLTLLFPA---ASQANDELIKLQEDPNQWVMWGGDYSG
TRYSPLNQINAENAKNLQVAWTFSTGVLRGHEGGPLVL--------GD-TMYIHTPFPNK
VFAIDLKTK-ALKWEYEPKQD---------ASV-IAVMCCDTVNRGLAY-ADGKIFLQQA
DTTLVALDQNTGKVVWKVVNG-----DPKVGA----------------TNTNAPLVVKDK
IITGISGGEFGVRGYLTAYNIKDGSVAWRAYSTGPDEEMLVDPE--KTTEML--------
---------------------KPIGKDSSLESW------------------EGDQWKIGG
GTTWGWFSYDPELNLIYYGTGNPSTWNPSQRPGDNKYSMTIMARDADTGMAKWLYQMTPH
DEWDYDGVNEMILVDKK-FKGKDRKLLVHFDRNGFGYTLDRETGELLVAEKFDPAVNWAT
HVDMET-----GRPQVVAKYSTAQNGE--DVDTKGICPAALGSKDQQPATYSPRTGLFYV
PTNHVCMNYEPFEV-----SYTAGQPYVGATLSMFPA--------PDS---HGGLGNFIA
WDAEKGEIVWSLPEPFSVWSGALATGGGVVFYGTLEGYLKAVDE-KTGKELYRF-KTPSG
IIGNVNTYMHDGKQYVAVLSGIGGWAGI-----GMAAGLEG----DT-----DGLGAVGA
YRKLSDYTQL--GGVLTVFALPN--------------
>
QBC28924_Methylomonas_LW13_c__Gammaproteobacteria_o__Methylococcales_f__Methylomonadaceae_characterized
------MKKP-VKNWLLAST-VATLLAAPT---VSTANSDVEKLTQDPANWATWGGDYAG
TRYSKLSQINAQNVKNLQPAWSFSTGVLRGHEGGPLVV--------NG-VMFIHTPFPNT
VYAIDQKTK-AVIWEFTPTMD---------ADVTIPVMCCDTVNRGLAY-GDGKIFLQQS
DTVLTALDAKTGKRVWSVQNG-----DPKLGM----------------TNTNAPLVVKDK
VITGISGGEFGVRGFLAAYNIRTGQLEWKGYSMGPDKDTLLKPG--KSTTWQD-------
----------------GK--VTPVGADSGTSTW------------------KGDQWKIGG
GTTWGWYSYDPKLNLVYYGSGNPSTWNPVQRPGDNKWSMSLWARDADTGEVKWVYQMTPH
DEWDYDGINETVLVDQE-VKGKMHKTIVHFDRNGFGYTLDRETGELLVAEKFDKSVNWAS
HVDLKS-----GRPEVVPEFSTEHNGE--DVNTVGTCPAALGSKNQQPVSYSPQTGLFYI
SGNHLCMEYEPFEV-----SYTAGQPYVGATLSMMPAGADVLTGKKDG---TTNLGQFTA
YDAKTGKIAWSNKEQFSVWSGSVATAGGVVFYGTLEGYLKAVDA-KTGKELYKF-KTPSG
IIGNVNTWEFEGKQYVGVLSGIGGWAGI-----GIAAGIDD----GTSASSSEGLGAVGA
YRSLSSYTKL--GGTLTVFALPN--------------
>MBH71000_Alphaproteobacteria_SP38_o_GCA-002720895
------MKKS-MKIALLASS----AIAIAA---GASANSGLAKLINNSNNWAHPSGDFNL
HRHTKLKQINKSNVKDLKVAWTFSTGVLRGHEGGPLVI--------GD-TLYIHSAFPNN
VYALDLNEPGRIIWEYNPKQD---------PDV-IPVMCCDTVNRGLSY-GDGKIFLQQA
DTVLVALDAKSGKLIWSATNG-----DPSVAQ----------------TNTNAPFVMKDK
VITGMSGAEFGVQAWINAYDIKTGKRVWRGYSMGPDDQTLIVPG--KSKGSWG-------
----------------GKSWGVALPANSGTSTW------------------EGDQWKIGG
GSVWGWWPYDKANNSLYYGNGNPSTWNPVQRPGDNKWSMSMFSRDVDSGVVNWVFQMTPH
XEWDYDGINESIMADIK-VGGKTRKTIVHFDRNGFAYTWDRKTGEPLVIEKYDPATNWSR
GQSLKT-----GLHDRVKKYSTEAGGE--DVNTKNICPAALGTKDQQPAAXDPKRATFAV
PTNHVCMDYEPFRV-----SYTAGQPYVGATLSMFPA--------PGG----THLGNFIS
WDAGKGKINWSKKEPFSVWSGALTSAGDVWYYGTLEGYIKAVDP-DNGKTLWKF-RTPSG
IIGNVXXWAHGGKQXIGVLSGVGGWAGI-----GLAAGLTE----DT-----AGLGAVGA
YKALHNYTKL--GGTLTAFSL----------------
>Ga0456544_00244_35_1837_108m_BATS_129_RPKM
------MKKS-MKIALLASS----AIAIAA---GASANSGLAKLINNSNNWAHPSGDFNL
```

```
HRHTKLKQINKSNVKDLKVAWTFSTGVLRGHEGGPLVI--------GD-TLYIHSAFPNN
VYALDLNEPGRIIWEYNPKQD---------PDV-IPVMCCDTVNRGLSY-GDGKIFLQQA
DTVLVALDAKSGKLVWSATNG-----DPSVAQ----------------TNTNAPFVMKDK
VITGMSGAEFGVQAWINAYDIKTGKRVWRGYSMGPDDQTLIVPG--KSKGSWG-------
----------------GKSWGVALPANSGTSTW------------------EGDQWKIGG
GSVWGWWPYDKANNSLYYGNGNPSTWNPVQRPGDNKWSMSMFSRDVDTGVVNWVFQMTPH
DEWDYDGINESIMADIK-VGGKTRKTIVHFDRNGFAYTWDRKTGEPLVIEKYDPATNWSR
GQSLKT-----GLHDRVKKYSTEAGGE--DVNTKNICPAALGTKDQQPAAFDPKRATFAV
PTNHVCMDYEPFRV-----SYTAGQPYVGATLSMFPA--------PGG----THLGNFIS
WDAGKGKILWSKKEPFSVWSGALTSAGDVWYYGTLEGYIKAVDP-DNGKTLWKF-RTPSG
IIGNVNGWAHGGKQYIGVLSGVGGWAGI-----GLAAGLTE----DT-----AGLGAVGA
YKALHNYTKL--GGTLTAFSL----------------
>MBK31765_SP289_c__Gammaproteobacteria_o_UBA4486
------MSKLSLKKMLIAIL----ALGISA---SVVANDEVRKLNANPNYWAFPGGDYNN
WRYTELNQINTKNASKLVNAWTFSTGRLQGHEGGPLVLPGSATGRSND-TLYIHSSFPND
VFAIDLDTL-EIVWYFEPQQD--------EAET-VPVMCCDIVNRGLGY-SMGQIYLQQA
DTLLVALDAATGKVKWTAENGREIGYGPAAGD----------------TNTNAPHPIKDK
VFTGCSGAEFGVRCWIAAFNAKDGSLAWRAFSMGPDEDIIFDAN---TTSLG--------
---------------------KKVGKNSSLNSWCSNADWKAPATGPNKCKGLSDAWKSGG
GSVWGWWPADFDENLVYYGNGNPSTWNPVVRPGDNKWAMTIFARDIDTGVARWVYQMTPH
DEWDYDGVNEMILADIK-VKGKMTPALVHFDRNGFAYTMNRKNGALLVAEKYDPAVNWAT
HIDMKT-----GYPQTLNQYSTHYNGE--DVVSKDICPAALGTKDQQPAAFSPRTNLFYV
PTNHVCMTYEPVSYAAGGNQYTAGAAYVNAALTMYPAGAVCPDC-PNNNAAKDNMGNMIA
WDAGKGKIVWNIVEQWSVWSGVLTTAGDVVFYGTLDAYAKAVDA-KTGKLLWKH-KAPSG
IIGNFNSWSHKGKQYVGVLSGIGGWAGAVVSIEGLAAAPDD-----------AALGAVGG
YRKLLNSSRN--AGVLMVYALP---------------
>Ga0509209_000470_14_2014_BATS_110m_260_RPKM
------MSKL-IKKMLIAIL----ALGICA---PAFANDEVRKLIDNPNYWAFPGGDYNN
WRYTELNQINTKNASKLVTAWTFSTGRLQGHEGGPLVLPGSATGLSGD-TLYIHSSFPND
VFAISLDTL-EIVWYFEPQQD--------EAET-VPVMCCDIVNRGLGY-SMGNIYLQQA
DTLLVALNAATGKVVWTAENGREIGYGPAAGD----------------TNTNAPHPIKDK
VITGCSGAEFGVRCWIAAFNAKDGSLAWRAFSMGPDEDILFDAN---TTSLG--------
---------------------KKVGKNSSLNSWCANADWKAPSRGPNKCNGLSDAWKMGG
GSVWGWWPADFDENLVYYGNGNPSTWNPVVRPGDNKWSMTMFARDIDTGVARWVYQMTPH
DEWDYDGVNEMILADIK-VKGKMTPALVHFDRNGFGYTMNRKTGALLVAEKYDPAVNWAT
HVDMKT-----GYPQTLDKYSTHHNGE--DVVSKDICPAALGTKDQQPAAYSPRTNLFYV
PTNHVCMTYEPVSYAAGGNQYTAGAAYVNAALTMYPAGAVCPEC-PNNNAQNDNMGNMLA
WDAGKGKIVWNIVEQWSVWSGVLATAGDVVFYGTLDAYAKAVDA-KTGKLLWKF-KAPSG
IIGNFNSWAHKGKQYVGVLSGIGGWAGAVVSIEGLAAAPDD-----------AALGAVGG
YRKLLNSSRN--SGVLMVFSLP---------------
>MAS84057_NAT232_c__Gammaproteobacteria_o_UBA4486
------MSKL-IKKMLIAIL----ALGICA---PAFANDEVRKLIDNPNYWAFPGGDYNN
WRYTELNQINTKNASKLVTAWTFSTGRLQGHEGGPLVLPGSATGLSGD-TLYIHSSFPND
VFAISLDTL-EIVWYFEPQQD--------EAET-VPVMCCDIVNRGLGY-SMGNIYLQQA
DTLLVALNAATGKVVWTAENGREIGYGPAAGD----------------TNTNAPHPIKDK
VITGCSGAEFGVRCWIAAFNAKDGSLAWRAFSMGPDEDILFDAN---TTSLG--------
---------------------KKVGKNSSLXSWCANADWXAPSXGPXKCNGXSDAWXXGG
GSVWGWWPADFDENLVXYGNGNPSTWNPVVRPGDNKWSMTMFARDIDTGVARWVYQMTPH
```

```
DEWDYDGVNEMXLAXIK-VKGKMTPAXVHXDRNGXXXTMNRKTGALLVAEKYDPAVNWAT
HVDMKT-----GYPQTLDKYSTHHNGE--DVVSKDICPAALGTKDQQPAAYSPRTNLFYV
PTNHVCMTYEPVSYAAGGNQYTAGAAYVNAALTMYPAGAVCPEC-PNNNAQNDNMGNMLA
WDAGKGKIVWNIVEQWSVWSGVLATAGDVVFYGTLDAYAKAVDA-KTGKLLWKF-KAPSG
IIGNFNSWAHKGKQYVGVLSGIGGWAGAVVSIEGLAAAPDD-----------AALGAVGG
YRKLLNSSRN--SGVLMVFSLP---------------
>644897261_Methylotenera_mobilis_JLW8_XoxF4-1_Mmol_2048_characterized
----MQINKIKLALGLVAG------MAMPL---IASAAADQEAAMKDANNWAHPRGQHNN
NGYSALSQVNKGNVKNLKAAWTFATGVNRGHEGSPIVV--------GN-MMYVHTAFPNN
VYALDLDNNQKIVWSYFPKQD---------PSV-QAVLCCDNVSRGLGF-GDGKIYLQQN
DGNLVALDAKTGAKVWSTLIN-----DPKVGA----------------TNTNAPHVIKDK
VLTGCSGAEFGVRCFIAAYNAKDGSLAWKAYSTGPDSEVLIGADFNKETPEYSALSVYQD
VNGGNKEGGSFKKLPNDQ--IKFGEKDLGTRTWL-------------KPQAVKDGWQHGG
GSVWGWWPYDARTNLVYYGTGNPSVWNPDVRPGDNKWSMTVFARDLDTGVAKWGMQMTPH
DEWDYDGINEVILFD-----KGGKTYAWHHDRNGFAYTWNAHNGKLQAAQKVHPFVNWAT
DVDMKT-----GVPNKLASASTHQ-----DYNAKGVCPAALGTKDQQPAAYSPKTGLVYS
PLNHVCMTYEPVES-----KYVAGQPWVGATLTMFPG--------PDG-----VMGGFGA
YDPMTNKKVWYNKEKFSAWGGALTTATDLVFYGTLDRWFKAVDA-KSGKELWKF-QVGSG
VIGNAFTYGNKGKQYVGVLSGIGGWAGV-----AMNLGMTT----PT-----DALGAAGG
YAELVKYNAAPGGGALNVFSL----------------
>Ga0453800_00330_46_1923_SanFranciscoBay_Methylophilaceae_BS307-5m-G33_214_RPKM
----MQLNKIKLALGFVAGL----TVAMPM---VASAAADQEKAMANANNWAHPRGQHDN
QAYSKLTQVNKGNVKNLKAAWTFATGVNRGHEGAPLVI--------GN-TMFVHTAFPNN
VYALDLNNDQKIVWSYFPKQD---------PSV-QAVLCCDNVNRGLGF-GDGMIFLQQN
DGMLVALDAKTGAKKWDVSNT-----DPKVGA----------------TNTNAPHVIKDK
VLTGCSGAEFGVRCFIAAYNIADGSLAWKAMSTGPDSEVLIGADFNKENPLFSALSVYED
VNGGNKQGGSFKKIPTDQ--LQGGVADLGVKTWL-------------KPQAVKDGWQHGG
GSVWGWWPYDAKTNLVYYGTGNPSVWNPDVRPGDNKWSMTVFARDLDTGMARWGMQMTPH
DEWDYDGINEVILFD-----KGGKTYAWHHDRNGFAYTWQADNGTLVAAEKVHPFVNWAT
DVDLKS-----GVPNKVAEHSTHQ-----DYNTKGTCPAALGTKDQQPAAYSPKTGLIYS
PLNHVCMTYEPVES-----KYVAGQPWVGATLTMFAG--------PDG-----VMGGFAA
YDPMTNKKVWYNKEKFSAWGGALTTDSDLVFYGTLDRWLKALDA-KSGKELWKF-QVGSG
VIGNAFTYGHKGKQYVGVLSGIGGWAGV-----AMNLGLTN----DT-----DALGAAGG
YKELTKYNAAPGGGALTVFSL----------------
>EAV46784_Methylophilales_HTCC2181_o__Burkholderiales_f__Methylophilaceae_g__BACL14
MEINMQLNKIKLALGFAAAA----TMAMPM---VASAAADQEKAMSNANNWAHPRGQHDN
QAYSKLTQLNKGNVKNLKAAWTFATGVNRGHEGSPLVI--------GN-MMYVHTAFPNN
VYALDLNNDQKIVWSYFPKQD---------PSV-QAVLCCDNVNRGLGF-GDGKIFLQQN
DGMLVALDAKTGAKVWDASNV-----DPKVGA----------------TNTNAPHVIKDK
VLTGCSGAEFGVRCFMAAYNIADGSLAWKAMSTGPDSEVLIGADFNKENPLYSALSVYED
VNGGNKQGGSFKKIPTDQ--LQGGVADLGVKTWL-------------KPQAVKDGWQHGG
GSVWGWWPYDAKTNLVYYGTGNPSVWNPDVRPGDNKWSMTVFARDLDTGMARWGMQMTPH
DEWDYDGINEVILFD-----KGGKTYAWHHDRNGFAYTWTAAEGTLVAAEKVHPFVNWAT
DVDLKS-----GVPNKLAEHSTHQ-----DYNTKGTCPAALGTKDQQPAAYSPKTGLIYS
PLNHVCMTYEPVES-----KYVAGQPWVGATLTMFAG--------PDG-----VMGGFAA
```

```
YDPMTNKKVWYNKEKFSAWGGAMTTASDLVFYGTLDRWFKALDA-KSGKELWKF-QVGSG
VIGNAFTYAHKGKQHVGVLSGIGGWAGV-----AMNLGLTN----DT-----DALGAAGG
YKELTKYNAAPGGGGLTVFSL----------------
>Ga0206682_100207233_Monterey Bay_40m_342_RPKM
----MKLNKIKLALGFVTAA----AMAMPI---VASAAADQEKAMKNSANWAHPRGQHDN
QAYSKLTQINKGNVKNLKAAWTLSTGVNRGHEGSPLVI--------GN-MLYVHTAFPNN
VYALDLNNNQKVVWSYFPKQD---------PSV-QAVLCCDNVNRGLGF-GNGKIFLQQN
DGNLVALNAKTGAKVWSTLNT-----DPKVGA----------------TNTNAPHVIKDK
VLTGCSGAEFGVRCFIAAYNIEDGSLAWKAMSTGPDSEVLIGADFNKENPLYSALSVYED
VNGGNVEGGSFKRIPNDQ--LQAPVADLGLKTWL-------------KPQAVKDGWQHGG
GSVWGWWPYDAKTNLVYYGTGNPSVWNPDVRPGDNKWSMTVFARDLDTGMARWGMQMTPH
DEWDYDGINEVILFD-----KGGKTYAWHHDRNGFAYTWVASNGTLVSAEKVHPFVNWST
GIDLKS-----GVPQKEAAHSTHQ-----DYNTKGTCPAALGTKDQQPAAYSPKTGLIYS
PLNHVCMTYEPVES-----KYVAGQPWVGATLTMFAG---------PDG-----VMGGFAA
YDPMTNKKVWYNKEKFSAWGGALTTASDLVFYGTLDRWFKALDA-KSGKELWKF-QVGSG
VIGNAFTYSHKGKQHVGVLSGIGGWAGV-----AMNLGLTN----DT-----DALGAAGG
YKELTKYNAAPGGGALTVFAL----------------
>CAH7806198_Methylopumilus_planktonicus_o__Burkholderiales_f__Methylophilaceae
----MKINKIKLALGLVAGL----SVAMPM---IASAAADQEKAIANPDNWAAPRGDYTN
QAYSKLTQINQGNVKNLKAAWSFATGVNRGHEGSPVVI--------GN-MMYVHTAFPNN
IYALDLNDNQKIVWSYFPKQD---------PSV-QAVLCCDNVSRGLGY-GDGKIFLQQN
DGTLVALDAKSGKKIWDAKIT-----DPKVGA----------------TNTNAPHVIKDK
VLTGCSGAEFGVRCFIAAYTL-DGKLAWKAYSTGPDAEVLIGKDFNKDTPLYSALSVYED
VNGGNKQGGSFKKIDTAK--LKGGEKDLGVKTWL-------------KPQAVKDGWQHGG
GSVWGWWPYDPKTNLVYYGTGNPSVWNPDVRPGDNKWSMTVFARDLNTGVARWGMQMTPH
DEWDYDGINEVILFD-----NGGKTYAWHHDRNGFAYTWQADNGVLVAAEKVHPFVNWAT
DIDKKS-----GVPNKLATASTHQ-----DYNAKGVCPAALGTKDQQPASYSPKTGLIYS
PLNHVCMTYEPVES-----KYVAGQPWVGATLTMFAG---------PDG-----VMGGFAA
YDPMTNKKAWYNKEKFSAWGGSLNTASDLVFYGTLDRWFKALDA-KTGKELWKF-QVGSG
VIGNAFTYTNGGKQHVGVLSGIGGWAGV-----AMNLGLTN----PT-----DALGAAGG
YAELTKYNAAPGGGALTVFSL----------------
>644896980_Methylotenera_mobilis_JLW8_XoxF4-2_Mmol_1770_characterized
MN-HLFKRKFGVSLALSAGL----FGMLST---PAFAEDELSTLMKDDNQWVHPRKDFSN
SGYSRLSQINKGNVKNMKLAWSFATGVNRGHEGAPLVV--------NG-VMYVHTAFPNN
VYALDLNDNQKILWSYFPKQD---------PST-QALLCCDNVNRGLGY-GEGKIFLQQN
DGLLVALDAKTGKKLWDTQVT-----NPKEGA----------------SNTNAPHVIKDK
VITGCSGGEYGVRCYISAYNIKDGALVWRAYSTGSDQDVLIGNDFNKDNPHYSALSLYQD
VNGGNKIGGSFKALPKDQ--LKFPEKDLGVKTWL-------------KPQAKKDGWQNGG
GPTWGWYSYDPKLNLMYYGTGNPSTWNPDVRPGDNKWSMTIFARDVDTGEAKWGYQMTPH
DEWDYDGMNEIILWE-----AKGKKLASHFDRNGFGYTFDRTNGSLLVANKMHPFVNWAF
NVDLKT-----GIPNKDPKFSTHQ-----DYNAQGICPSALGVKDQQPAAFSPKTGMFYV
PLNHVCMTYEPVES-----KYVAGQPWVGSTLTMFPG---------PDG-----VMGGFMA
WDGLKGKSAWYKKEQFSAWGGALVTSTNLVFYGTLDRWFKAVDA-NNGKELWKF-QVGSG
VVGNAITYSNKGKQYVGVLSGIGGWAGV-----AMNLGLTN----ST-----DGLGAAGG
YKDLVKYNAAPGGGAMNVFSL----------------
```
