## Supplementary material for "Diel cycle of lanthanide-dependent methylotrophy by TMED127/Methylaequorales bacteria in oligotrophic surface seawater": NuoL_alignment

```
>WP_085441360_Magnetofaba_australis_c__Magnetococcia
-----MSTYKLIVLLPLMGSIIAGLF----GRRLGNQMSQIVTIGGIGLALLLSIKAFFE
IALG-DAPAVHETF-FTWIPSGDFVVTLGVLVDRLTAIMLIVVTGVSTLVHIYSVNYMEE
DPDVPRFFSYLSLFSFAMLSLVTAPNFLQLFFGWEGVGLASYLLIGFWFKKESACNAAIK
AFLVNRVGDFGFALGVFGVFMVFGTLDFLGDKGVFMALTNY----QA--TMMFLGHEFNT
MTLICLLLFMGAMGKSAQFFLHTWLPDAMEGPTPVSALIHAATMVTAGVFLVCRASPLF-
EQSETALMVVTVIGAVTAIFAASVGLVQNDIKRVIAYSTCSQLGYMFFAAGVSAYAASMF
HLMTHAFFKALLFLGAGAVIHAMHHEQDMRKMGGLWKKIPLTYALMMIGTLALT-----G
FPY-----LA---GWWSKDAILESAWAAHTGVGTFAWVIGLIAAFMTTFYSFRLVFMTFH
GKPH-------------DEHHYDH--------------A-HEAPWFMRAPNLLLAVGALL
AGYLGHGIIEA-------GWF-KDAIFLAE-GHDALA-HAHH-APAHVKWLPFVMFLGGL
FLALLMYIWVPT---LPK------KVSELCPRGYQFLLNAWYFDKLYDAIFIKPAKAIG-
KGLWQTGDATIIDGYGVNGTANLMVRMGAVLKRMQSGYVYHYAFAMLAGVLVLITFYA--
---------------
>WP_011715163_Magnetococcus_marinus_c__Magnetococcia
-----MSTYKLIVLLPLLGSLIAGLL----GRTIGTRMSQMVTIGGIGLSLLLSIQAFWQ
IAVN-DGEVVREIF-WSWVISGDFQVTLGVLVDRLTAVMLIVVTGVSTLVHIYSVDYMHE
DPDNPRFFSYLSLFSFAMLMLVTSPNFLQLFFGWEGVGLASYLLIGFWFKKESACNAAIK
AFLVNRVGDFGFALGVLAIFMVFGTLDY---SAVFGAARGEF---NQ--TMHFLGYEFTT
ITLICLLLFLGAMGKSAQLFLHTWLPDAMEGPTPVSALIHAATMVTAGVFLVCRASPLF-
ELSETALAVVTIIGALTAFFAATVGMVQNDIKRVVAYSTCSQLGYMFFAAGVSAYAASMF
HLMTHAFFKALLFLGAGAVIHAMHHEQDMRKMGGLVKKIPLTYGLMMIGTLALT-----G
FPG-----LA---GFFSKDAILESAYAAHSATGTFAFWLGIMAAGMTTFYSFRLVFMTFH
GKPK-------------DHHAYDH--------------A-HEAPWFMRGPNIALAIGALF
AGYLGVPLIEA-------GWF-KEAIVLAA-GHNALE-HAHH-VPAWVKWLPFVMFVLGL
SVAVVLYVLAPT---LPA------KIAQMCPRGYNFLLNAWYFDKLYDKLFVKGAQCLG-
QGLWKTADETIVDGYGVNGWARTLGRWGASLRRMQSGYVYHYAFAMFVGVLALATFYS--
---------------
>WP_090018703_Limimonas_halophila_o__Kiloniellales
-------MEVVAVFLPLIGAALAGLF----GGVLGDRGSQLVTCGLLIVSAVLSVIVFVD
VAL--YDNARTTEL-FTWFASGDLELSWAIRMDTLSAVMLATVTVISALIHVYSIGYMEH
DASIQRFFSYISLFTFFMLMLVTADNFLQLFFGWEGVGLASYLLIGFWYQKPSANAAAIK
AFLVNRVGDIGFALGIAATFFVFQTTSF---DQVFAAAPEMA---DT--GFSFLGIEAPA
LTIISILLFIGAMGKSAQLGLHTWLPDAMEGPTPVSALIHAATMVTAGVFMLARVSPIL-
EQAPTALAVITIIGALTAFFAASVGMVQNDLKRVIAYSTCSQLGYMMFAVGVSAYGAAIF
HLMTHAFFKALLFMGAGSVLHAMDEEHDMRKMGGIWRMLPITYAVMWIGSLALA-----G
IPP-----FA---GFFSKDMILEAAYAAHSGTGQLAFWLGVIAAGMTAFYSWRLLFMTFH
GKPR------------ASAETMKH--------------V-HESPKVMTLPLMALAVGAVL
AGWLGYSAFVG---HGFEHFW-SGSIL----GHEAIK-AAHH-VPLWVKLLPLVVSVGGI
AVAYVMYIARPE---LPA------ALASRLRPVHAFLYNKWYFDELYDLVFVRPAWALG-
NGLWRAGDIAVIDGLGPNGVSSVARGLARRASQLQTGYVYHYAFAMLIGVVALVTWYMM-
---TQIG--------
>RMD64542_Alphaproteobacteria_J153_o_Kiloniellales
-------METLIVLLPLIGAAIAGLF----GRFIGDRGAQLVTCGLLGLSAILSIFVFKD
VAI--DGNTRVTEL-FTWVDSGTFEVSWAVKLDTLSAVMLLVVTVISSLIHVYSIGYMAH
DKSIPRFFSYLSLFTFFMLALVTADNLVQMYFGWEGVGLASYLLIGFWYTRPSACAAAIK
AFLVNRVGDAGFALGIVGTFMLFGAVGF---EEIFATAPQMA---GA--TLTFIGLEGHA
LTILCLLLFIGAMGKSAQIVLHTWLPDAMEGPTPVSALIHAATMVTAGVFMVARLSPMF-
```

```
EYAPAALEVVTVVGATTAFFAATVGLTQFDIKRVIAYSTCSQLGYMFFAAGVSAYSAAIF
HLMTHAFFKALLFLGAGSVIHAMSDEQDMRRMGGIWRMIPVTYAVMWIGSLALA-----G
IPP-----FA---GYFSKDMILEAAYAAHSAVGSYAFWLGIAAAFMTAFYSWRLLFMTFH
GSPR------------ADAEVMAH--------------V-HESPKVMIIPLLALALGAVF
AGWIAVDAFVG---EGRDEFW-GEAILVLA-SNDAIE-AAHH-VPGWVKILPLIMGVSGI
ALAWVFYVARPD---LPG------RVAAAARPVYLFLYNKWYFDELYDFLFVRPAFYLG-
RGLWRGGDGAVIDGVGPDGIAAATWDLARRIAALQTGYLYHYAFAMLIGVVGLATWYVV-
---TLIG--------
>WP_193370550_Pelagibius_NBU2595_o_Kiloniellales
-------MDQLIVLLPLLGAIIAGFF----GRLIGDRGAMYVSSGLLLISMVLSCIVFYD
VAF--AHNARVTEV-FTWIDSGSFEVSWALKVDTLTAVMLIVVTGVSSMVHVYSIGYMSH
DPYKPRFFAYLSLFTFFMLMLVTADNFVQMFFGWEGVGLASYLLIGFWYERPSACAAAIK
AFLVNRVGDIGFALGICGAFVLFGTAGF---DQVFASVPGMA---DA--QFEAFGISGHA
LTIICILLFIGAMGKSAQLGLHTWLPDAMEGPTPVSALIHAATMVTAGVFMVARLSPMF-
EYAPTALELVTYVGALTAFFAASVGLTQNDIKRVIAYSTCSQLGYMFFALGVSAYSAGIF
HLMTHAFFKALLFLGAGSVIHAMSDEQDMRKMGGIYKMIPGTYLLMWIGTIALL-----G
VGIPGVFGFA---GFYSKDIVVEAAYASHSTHGTFAFVMGVVAAFMTAFYSCRLMFMTFH
GTPR------------ASKEVMSH--------------V-HESPQVMLVPLYVLAFGAVF
AGLAAYEYFVG---QHMEEFW-GEALKVLP-ENNSVE-AAHH-VPLWVKLSPLVAGLGGI
GLAYVFYIASPA---LPA------KTAKALRPLYLFSLNKWYFDELYDRVFVRPARVLG-
FGLWKGGDGAVIDRLGPDGIAATTQNASRRTSLLQSGYVYHYAFAMLIGVFVLVTWYFL-
---SQLG--------
>WP_354078307_Constrictibacter_MBR-5_o_SHVR01
-------MYAAIVFLPLVGALVAGFG----GRFIGDRGAQVATCGAMLLAALLSVVAFFD
VAV--GGSPQTVVL-FDWIRSGTFEATWALRFDTLTAVMLIVVTIVSAAVHVYSVGYMSH
DPHIPRFMAYLCLFTFAMLMLVTADNLLQMFFGWEGVGLCSYLLIGFWFDRHSANAAAIK
AFVVNRVGDFGFIMGILGVYLVFDTIQF---DTIFAAAPGQA---GS--SLEFLGMQFPT
LEIICFLLFVGAMGKSAQVPLHTWLPDAMEGPTPVSALIHAATMVTAGVFMVARLSPVF-
EYAPTTLGIITVIGATTAIFAASVGLVQNDIKRVIAYSTCSQLGYMFFAIGVSAYPAAIF
HLMTHAFFKALLFLGAGSVIHAMSDEQDMRRMGGIWREIPITYAMMWIGSLALA-----G
VPI-----FA---GYYSKDMILESAWGAHSAVGDYAFWLGILAAAMTAFYSWRLIFMTFH
GKPR------------ADEHVMAH--------------V-HESPPVMLAPLFVLAVGAVL
AGIVGYGAFVG---DGMADFW-NGAILILP-EHHAME-NAHH-APFWVKIAPLVVGVVGI
AVAFLLYIRDPA---MPG------RIAAKAQGLYRFLLNKWYFDELYDRIFVRPAAWLG-
VGFWKGGDGAVIDGLGPNGLAAATRGLARQASRLQTGYVYHYAFVMLIGVVVLVTWYLY-
---RVGG--------
>SDF10219_Thalassobaculum_litoreum_o_Thalassobaculales
-------MYAAIVFLPLIGALIAGFG----NKKLGDRGAQIVTCGAMLTSAVLGIVAFWE
VAL--QGNAQTVEL-FTWIDSGSFEASWALRWDTLTAVMVIVVTVVSSCVHVYSVGYMSH
DPSIPRFMAYLSFFTFAMLMLVTSDNLIQMFFGWEGVGVASYLLIGFWYDRPSANAAAIK
AFVVNRVGDFGFALGIFGCFLLFDAVSF---DAIFAAAPEMA---GT--TFGFLWWEEVD
ALTVAILLFIGAMGKSAQLGLHTWLPDAMEGPTPVSALIHAATMVTAGVFLVARMSPLF-
EYAPTALVVVTVVGAATAFFAATVGTTQNDIKRVIAYSTCSQLGYMFFALGVSAYPAAIF
HLMTHAFFKALLFLSAGSVIHALHDEQDMRNMGGIWRKIPWTYAMFWIGSLALA-----G
IPF-----FA---GYYSKDMILESAFAAHTGAGQFAFWAGIVAALLTAFYSWRLLFMTFH
GKPR------------MDQHTFDH--------------A-HESPPVMLVPLIVLAIGAVF
SGFIGYEYFVG---HEMDAFW-GSAIKVLE-ENNVIE-AAHH-SPTWVKLLPLVVGVIGI
GAAYVAYILNPG---IPA------KVTSAIRPVYLFLLNKWYFDELYDWLFVKRSWQAG-
```

```
LMFWKTGDGTIINGFGPDGVSAMTRWCAGQVARLQTGYLYHYAFAMLIGVVALVSWYLA-
---FGG---------
>EDP66356_Alphaproteobacterium_BAL199_o_Thalassobaculales
-------MYSAIVFLPLVGALIAGFG----GRVIGDRGAQIVTCGAMIVSSVLALVAFWE
VAI--GGRTFTVDV-LTWIASGSFEANWALRWDTLTAVMVLVVTVVSTMVHVYSVGYMSH
DASIPRFMAYLSIFTFAMLMLVTADNLVQLFFGWEGVGVASYLLIGFWFHKPSANAAAIK
AFVVNRVGDFGFALGIFGCFMLFDTVSL---DVIFAAAPEKA---GT--TLNFLGMQGDA
LTIIAILLFIGAMGKSAQLGLHTWLPDAMEGPTPVSALIHAATMVTAGVFLVARMSPLF-
EYAPIALQVVTIVGAATAFFAATVGMCQNDIKRVIAYSTCSQLGYMFFALGVSAYPAAIF
HLMTHAFFKALLFLSAGSVIHALSDEQDMRNMGGIWKKIPFTYAMMWIGSLALA-----G
VPL-----FA---GFYSKDMILESAYGSHTGVGQFAFWMGIAAALMTAFYSWRLIIMTFH
GTPR------------MSDEVLHH--------------V-HESPKVMLIPLAFLAAGAVL
SGWAGYEWFVG---EGMAEFW-RESIKILP-DNNSIA-AAHH-VPFWVKAAPLVVGVIGI
GLAYLMYMFRPH---LPG------LLAARFRPIYLFLLNKWYFDELYDWLFVKRSWQVG-
VAFWQGGDRAIIDGFGPDGVSAVARNLAARAGRLQTGYVYHYAFSMLIGVVALVSWYLA-
---FGG---------
>MBL27380_Rhodospirillaceae_SP28_o_GCA-2721365
-------MYVLSVFLPLIGAVIAGLF----GRVLGDRGSQLITSGAMVLSFVFAVVILFQ
VAI--GGDPRSVEL-LTWIQSGEFDVSWALQFDSLSAVMIFVVTGVSTLVHIYSIGYMSH
DPHKPRFMAYLSLFTFSMLMLVTADNFVQLFFGWEGVGLCSYLLIGFWYSKPSANAAAIK
AFLVNRVGDFGFALGIVAIFFVFDAVDY---ETVFTVAPDFA---GT--TTTFLSWEVDT
LTLICLLLFIGAMGKSAQIGLHTWLPDAMEGPTPVSALIHAATMVTAGVFMVARCSPLF-
ELSPFALGVVTVVGGTTAIFAATVGLVQNDIKRVIAYSTCSQLGYMFFACGVSAYQAGIF
HLMTHAFFKALLFLGAGSVIHAMSDEQDMRKMGGIWRMIPLTYALMWIGSLALA-----G
IPP-----FA---GFFSKDMILEAAWGAHSFVGGYAFWLGTIAAALTAFYSWRLLFMTFH
GEPR------------ADHHVMAH--------------V-HESPKVMTIPLILLAVGAVV
AGYLGLPMV-----EPEGEFW-GEAIYILP-GHHPLE-AAHH-APFWVKIMPLIMGVGGI
AVAYLFYIADPT---LPR------RLAGRFQGLYRFLLNKWYFDELYDRIFVRPAKAIG-
YGLWKSGDGDVIDGIGPDGISAAIVQIARRAGRLQSGYIYHYAFVMLIGVAAFITWYIV-
---GTGL--------
>PPR38919_Alphaproteobacteria_MarineAlpha9_Bin5_o_UBA7887
-------MYTTCIFLPLIASIISGLF----GRIVGDRGAQLITSGALTLSFLISLMIFND
VAL--EGNIHRVEL-LTWITSGSFQVAWVLQFDALTAVMVFVVTTVSAIVHIYSIGYMAH
DPHIPRFMAYLSLFTFAMLMLVTSNNLVQLFFGWEGVGLCSYLLIGFWYTRPSANAAAIK
AFLVNRVGDFAFALGIFAIFLTFGSIEF---DTIFATAPKLV---GH--TSIFLGMEVDT
LTLICLLLFVGAMGKSAQIGLHTWLPDAMEGPIPVSALIHAATMVTAGVFLVVRCSPLF-
ELSDTALIVVTFVGASTAIMAATIGLVQNDIKRVIAYSTCSQLGYMFFACGVSAYSAAIF
HLMTHAFFKALLFLGAGSIIHAMSDEQDMRKMGSIWRLIPITYALMWIGSLALA-----G
IPP-----FA---GYFSKDLILEAAYGAHSAIGNYAFWLGVIVVLMTAFYSWRLLFMTFH
GESR------------ADEKVVAH--------------I-HESSKVMLLPLFVLAAGSIG
AGYVGYNMT-----DLNSDFW-GQSIFVAS-SHQALE-NLHH-VPVWVKVLPITLAIVGI
VLAYLLYIRFPK---APA------KIVVSTRGLYYFLLNKWYFDEFYDWLIVKPIKRLG-
LGLWKSGDGDLIDGVGPDGIAAAVVNLSRRASNIQSGYIYHYAFAMLIGIALLVTWYIA-
---WAGR--------
>MEE2980351_Pseudomonadota_XB_MAG1186_o_JAAXGF01
-------MYTASIFLPLLGAALAGFF----GRALGARTSAVIACVMLGLAMVAGWYAFYE
VVF--NGQSRTIEL-ATWIDSGAFEISWALRFDVLTAVMVFVVTTTSFLIHVYSIGYMAH
DPHVPRFFAYLSLFTFAMLMLVSADNFVQMFFGWEGVGLCSYLLIGFWYTRPVACAAAIK
```

```
AFLVNRVGDLGFALGIMAIFFVFNAVEF---ATVFAAAPDYV---DT--SIEFLDVQFDT
LTVICLLLFVGAMGKSAQFLLHTWLPDAMEGPTPVSALIHAATMVTAGVFMVARLSPLF-
EYSPVALGVVTVIGATTAIFASTVALCQTDIKRVIAYSTCSQLGYMFFACGVSAYAAGIF
HLMTHAFFKALLFLAAGSVIHAMSDEQDMRKMGGLWRKIPVTYAVMWIGSLALA-----G
VPF-----FA---GYFSKDIILESAWGAHSMVGQYAFWLGVLAAVLTAFYSWRLLFMTFH
GAPR------------ASPEVMAH--------------V-HESPKVMTIPLMLLAVGAIF
AGYFGYKIGMV---DADGAFW-RGSILVLS-SHDALE-AAHH-VPLWVKGLPVVVAAAGI
IAAYVLYISVPG---LSD------KIATALRPVYLFLLNKWYFDELYDAVFVRPANYIG-
HGLWKAGDGAVIDGVGPDGFAAAAIDIAKRTVRLQSGYLYHYAFAMLIGVALLVTWYLV-
---MGVG--------
>WP_014745483_Tistrella_mobilis_o_Tistrellales
-------MYVALILLPLAAAIVAGFF----GRVIGDRGSQLVTTGAVGLAALLSVVAFFD
VAV--GGNATTVNL-FTWIRSGELAFSWALRIDSLTAVMLIVVNGVSFMVHIYSIGYMSH
DPHKPRFMAYLSLFTFAMLTLVTADNLVQMFFGWEGVGLASYLLIGFWYKRPSANKAAMK
AFVVNRVGDFGFSLGIFALFFVTGSVGF---DAIFADLGTVA---GT--SIDFLGMTLPT
IEVIAVLLFIGAMGKSAQLGLHTWLPDAMEGPTPVSALIHAATMVTAGVFMVARLSPVF-
ELTSFTSDMITVIGASTAFFAATVGLTQTDIKRVIAYSTCSQLGYMFFAAGVGAYPAAIF
HLMTHAFFKALLFLGAGSVIHAMSDEQDMRKMGGIWKLIPVTYTFMWIGSLALA-----G
LPP-----FA---GFFSKDMILETAYAAHTGVGAYAFWLGIAAAFMTAFYSWRLLFMTFH
GKPR------------ADKHTMDH--------------V-HESPLVMTAPLFPLAVGAIV
AGYIGLPMV-----DPELHFW-NGAITMLG-EHNILE-EAHH-VPGWVKLLPLVMSVGGI
ALAWFLYIRRPD---LPG------KIARDFSGLHKFLLNKWYFDELYDAIFVRPSLWIG-
RVLWQAGDRKIIDGLGPDGIAAVSVDLARRAGRMQSGYVYHYAFAMLIGLALIVSYFF--
---GLEA--------
>WP_188574487_Tistrella_bauzanensis_o_Tistrellales
-------MYVALILLPLAAAIVAGFF----GRVLGDRGSQFVTTGAVGLAALLSVVAFFD
VAV--YGNATTIEL-FTWIRSGGLDFAWALRIDALTAVMLIVVNGVSFMVHLYSIGYMSH
DPHKPRFMAYLSLFTFAMLTLVTADNLVQMFFGWEGVGLASYLLIGFWYKKPSANKAAMK
AFVVNRVGDFGFSLGIFGVFLVTGSVTF---DAVFGELGNVA---GT--TIDFLGMSLPT
IEVLGVLLFIGAMGKSAQLGLHTWLPDAMEGPTPVSALIHAATMVTAGVFMVARLSPLF-
ELTVFTSDMITVIGASTAFFAATIGLTQTDIKRVIAYSTCSQLGYMFFAAGVGAYPAAIF
HLMTHAFFKALLFLGAGSVIHAMSDEQDMRRMGGIWKLIPVTYMMMWIGSLALA-----G
IPP-----FA---GFFSKDMILETAYAAHSTVGAYAFWMGIAAAFMTAFYSWRLLFMTFH
GKPR------------ADKHTMEH--------------V-HESPLIMTVPLFLLAVGAIT
AGYIGQPMV-----DAGLAFW-NGAITMLG-EHNILE-EAHH-VPGWVKLLPLVMSAAGI
GLAWLFYIRRPD---LPG------KVASDFSGLYKFLLNKWYFDELYDAIFVRPALWIG-
RVLWQGGDRRIIDGFGPDGIAAVSLGLARRAGRMQSGYVYHYAFAMLIGLALIVSYFFF-
---GLEAGW------
>MBL6935107_Alphaproteobacteria_BS150m-G7_o_UBA11136
-------LYVATIFLPLIGAIAAGLF----GRWVGDRGAQLIACGMMVLSGLFSIIIFRD
VAL--DGNERVIEL-FTWIRSGGFEASWALKFDTLSAVMVFVVSVVSALIHIYSIGYMAK
DPSIPRFFSYLSLFTFFMLMLVSADNLVQLFFGWEGVGLCSYLLIGFWYDRDSANAAAIK
AFVLNRIGDFGFALGIFAVFYLFGSVSF---ETIFAAAPAQA---ET--SFEFLGLEVHA
LTLICLLLFIGAMGKSAQIGLHVWLPDAMEGPTPVSALIHAATMVTAGVFMVVRMSPLF-
ELAPSVLALVAVIGALTAFMAGTIALAQNDIKRVIAYSTCSQLGYMFFAIGVSAYGAAIF
HLMTHAFFKSLLFLGSGSVIHAMSDEQDMRKMGGIWRKIPLTYAVMWVGSLALV-----G
IPF-----FA---GYYSKDIIIESAWGAHSLVGNFAYVIGVVVAFLTAFYSCRLLFMTFH
GEPR------------ADEKVMAH--------------V-HESPKVMTVPLVLLAVGAIF
```

```
SGSLGYDLFVG---DGGEGFW-RASILVLP-GHDALA-EAHH-APWYIKLTPLVLAVLGM
ALAFLVYFMFPG---LAG------HAAERFPRVHAFLYNKWYFDEIYDRLLVRPALILG-
RGLWKSGDGELIDGIGPDGIAASVMRMGRRAAALQSGYVYHYAFAMIIGVVGFISWYLY-
---GLMG--------
>MAG23701_Rhodospirillaceae_ARS27_o_UBA11136
-------LYVATIFLPLIGAVVAGLC----GRWTGDRGAQLVTCGLMVLSGLFSIVIFRE
VAL--DGNERAIEL-FTWIRSGGLEAAWALKFDTLTAVMVFVVSVVSALIHIYSIGYMAK
DPSIPRFFSYLSLFTFFMLMLVSADNLVQLFFGWEGVGLCSYLLIGFWYDRDSANAAAIK
AFVLNRIGDFGFVLGIFAVFYLFGSVNF---ETIFTAAPDQA---ET--TFEFLGLEVHA
LTLICLLLFVGAMGKSAQLGLHVWLPDAMEGPTPVSALIHAATMVTAGVFMVVRMSPLF-
ELAPLVLALVAVVGALTAFMTATIALAQNDIKRVIAYSTCSQLGYMFFAIGVSAYGAAIF
HLMTHAFFKSLLFLGSGSVIHAMSDEQDMRKMGGLWRRIPLTYTVMWVGSLALA-----G
IPF-----FA---GYYSKDIIIESAWAAHGVVGNFTYILSVVVAFLTAFYSYRLLFMTFH
GESR------------ADERVMAH--------------V-HESPKVMTVPLVLLSVGAIF
SGYLGYDLFVG---DGGEEFW-RASILVLP-GHDALA-EAHH-VHWYIKIVPLVMAGLGI
LLAFVVYFMFPG---LAG------HAAERFPRLHTFLYNKWYFDEVYDRLLVRPALAFG-
RALWKTGDGELIDGIGPDGIAASVMRMGRRAAALQSGYVYHYAFAMIIGVVGFISWYLY-
---ELAG--------
>MBV23930_Rhodospirillaceae_NP1051_o_UBA11136
-------LYVATIFLPLIGAVVAGLC----GRWTGDRGAQLITCGLMVLSGLFSIVIFRE
VAL--DGNERAIEL-FTWIRSGGLEAAWALKFDTLTAVMVFVVSVVSALIHIYSIGYMAK
DPSIPRFFSYLSLFTFFMLMLVSADNLVQLFFGWEGVGLCSYLLIGFWHDRDSANAAAIK
AFVLNRIGDFGFVLGIFAVFYLFGSVNF---ETIFTAAPDQA---ET--TFEFLGLEVHA
LTLICLLLFIGAMGKSAQLGLHVWLPDAMEGPTPVSALIHAATMVTAGVFMVVRMSPLF-
ELAPLVLALVAVVGALTAFMTATIALAQNDIKRVIAYSTCSQLGYMFFAIGVSAYGAAIF
HLMTHAFFKSLLFLGSGSVIHAMSDEQDMRKMGGLWRRIPLTYXVMWVGSLALA-----G
IPF-----FA---GYYSKDIIIESAWAAHGVVGNFTYILGVVVAFLTAFYSCRLLFMTFH
GESR------------ADERVMAH--------------V-HESPKVMTVPLVLLSVGAIF
SGYLGYDLFVG---DGGEEFW-RASILVLP-GHDALA-EVRY-APQYIKIAPLAMAGLGI
LLAFVVYFMFPG---LAG------HAAERFPRLHTFLYNKWYFDEVYDRLLVRPALAFG-
RALWKSGDGELIDGIGPDGIAASVMRMGRRAAALQSGYVYHYAFAMIIGVVGFISWYLY-
---ELAG--------
>MBT3790209_Alphaproteobacteria_SI054_bin61__o_UBA2966
-------MYTALVFLPLLGAVLAGLF----GRILGDRGSQAVTLIGITTSALLSVYVFFD
VTG--NNNPVVVDL-FTWIESDSFEVSWALRVDQLTAVMLMVVCVVSAVVHWYSVGYMSH
DKAIPRFMSYLSLFTFAMLMLVTADNFLQMFFGWEGVGLCSYLLIGFWYDRPSANAAAIK
AFLVNRVGDFGFALGIMAIFFTFNSVEF---DVVFAAAPEMA---GK--TFMFLGQEWDI
LTTITMLLFVGAMGKSAQLGLHTWLPDAMEGPTPVSALIHAATMVTAGVFMLARCSYLF-
EYAPFTLEVVTVIGATTAFVAATIGLVQNDIKRVIAYSTMSQLGYMFFAIGVSAYPAAIF
HLLTHAFFKALLFLGAGSVIHAMSDEQDMRNMGGIYKLIPGTYLMMWIGSLALA-----G
VPL-----FA---GYYSKDMIIEVAFA-GSGVGTYAFTMGILAALMTAFYSWRLLFMTFH
GPAR------------ADEKVMAH--------------V-HESPKVMMLPLLVLALGAIF
AGWFWAGDMIG---DGRVAFWGGESIFIQP-ANHIFE-NAHH-VSAWVKYSPTALGAIGI
LLAWWFYIKNPN---IPK------AIATMHREAHLFLLNKWYFDELYDLIFVNPAKRLG-
HFLWKFGDGKIIDGLGPDGISRLVLNITRRAQTLQSGYVYHYAFAMMLGVAGFVSWFVF-
---GLGG--------
>MBT7747133_Alphaproteobacteria_SI037_bin135__o_UBA2966
-------MYTALVFLPLLGAVLAGLF----GRILGDRGSQAVTLIGITTSALLSVYVFFD
```

```
VTG--NNNPVVVDL-FTWIESDSFEVSWALRVDQLTAVMLMVVCVVSAVVHWYSVGYMSH
DKAIPRFMSYLSLFTFAMLMLVTADNFLQMFFGWEGVGLCSYLLIGFWYDRPSANAAAIK
AFLVNRVGDFGFALGIMAIFFTFNSVEF---DVVFAAAPEMA---GK--TFMFLGQEWDI
LTTITMLLFVGAMGKSAQLGLHTWLPDAMEGPTPVSALIHAATMVTAGVFMLARCSYLF-
EYAPFTLEVVTVIGATTAFVAATIGLVQNDIKRVIAYSTMSQLGYMFFAIGVSAYPAAIF
HLLTHAFFKALLFLGAGSVIHAMSDEQDMRNMGGIYKLIPGTYLMMWIGSLALA-----G
VPL-----FA---GYYSKDMIIEVAFASGSGVGTYAFTMGILAALMTAFYSWRLLFMTFH
GPAR------------ADEKVMAH--------------V-HESPKVMMLPLLVLALGAIF
AGWFWAGDMIG---DGRVAFW-GESIFIQP-ANHIFE-NAHH-VSAWVKYSPTALGAIGI
LLAWWFYIKNPN---IPK------AIATMHREAHLFLLNKWYFDELYDLIFVNPAKRLG-
HFLWKFGDGKIIDGLGPDGISRLVLNITRRAQTLQSGYVYHYAFAMMLGVAGFVSWFVF-
---GLGG--------
>MBT4017013_Alphaproteobacteria_SI053_bin31_o_UBA2966
-------MYTALVFLPLLGAVIAGLF----GRVLGDRGSQAVTLIGITTSALLSLYVFFD
VTG--NNNPVVIEL-FTWMESDSFEVSWALRIDQLTAVMLMVVTVVSAVVHWYSVGYMSH
DKAIPRFMAYLSLFTFAMLMLVTADNFMQMFFGWEGVGLCSYLLIGFWYDRPSANAAAIK
AFLVNRVGDFGFALGIMAIFFTFNSVEF---DVVFAAAPDVA---GK--TFVFLGQEWDI
LTTITMLLFVGAMGKSAQLGLHTWLPDAMEGPTPVSALIHAATMVTAGVFMLARCSPIF-
EFAPFTLEVVTIIGATTAFVAATIGLVQNDIKRVIAYSTMSQLGYMFFAIGVSAYPAAIF
HLMTHAFFKALLFLGAGSVIHAMSDEQDMRKMGGIYKLIPGTYLMMWIGSLALA-----G
VPL-----FA---GYYSKDMIIEVAFASNSGVGMYAFTMGILAALMTAFYSWRLIFMTFH
GPAR------------ADEKVMAH--------------V-HESPKVMMLPLLVLAFGAIF
AGWFWFDDMVG---DGRAAFW-GQSIFIQP-ANQIFE-NAHH-VSAWVKYSPTALGAFGI
LLAWWFYIKNPG---IPK------TIATAHREVHSFLLNKWYFDELYDLIFVRPAKRIG-
YFLWKIGDGKIIDGFGPDGISRLVLNITRRAQALQTGYVYHYAFAMMLGVAGFVSWFVF-
---GLGG--------
>MAG97780_Rhodospirillaceae_ARS1032_o_UBA2966
-------MYVALVFLPLAAAIVAGFF----GRLIGDRAAMLVTCLGVTLSALISFVVLYQ
VGM--QGRVEIVHL-FTWIDSQAFEVSWVLRFDQLTAVMLIVVTGVSALVHWYSIGYMSH
DGSKPRFFAYLSLFTFMMLMLVTADNFVQMFFGWEGVGLCSYLLIGFWYDRPSANAAAIK
AFIVNRIGDFGFALGIMAIFLIFKAVDF---DTVFAAAPDFA---GK--SLDFLGTDVDI
LTLITLLLFVGAMGKSAQLGLHTWLPDAMEGPTPVSALIHAATMVTAGVFMVARCSPLF-
EYAPISLAVVTIVGALTAVVTATIGLVQNDIKRVIAYSTCSQLGYMFFAAGVSAYSASIF
HLMTHGFFKALLFLGAGSVIHALADEQDMRKMGGLYTLLPLTYTVMWIGSLALA-----G
MPF-----FA---GFYSKDIVLEAAYAAHSQVGGFAFVLGVIAAFLTAFYSWRLLFMTFH
GKAR------------ASEKVMAH--------------A-HEAPKVMMIPLLLLAVGAMF
SGFLAYDLLVG---EG-STFW-GNSILVLP-ANDSVT-AAHH-VPLWVKILPTAIGVAGI
ALAWWFYIKRPA---IPE------RLASVHRELYLFLLNKWYFDELYDFLFVRPAKFIG-
WSLWRGGDGRIIDGFGPDGVAATVLRTARRATRIQTGYVYHYAFAMMVGLVLFVSWSLY-
---LMVGG-------
>MBM3510012_Alphaproteobacteria_K_Offshore_80m_m2_115_o__SHVV01
------MFYAAIVFLPLVGAIAAGLF----GRLLGVRASQLVTCLALTLSGVLATWGLYE
TAVI-GPDKQQIVL-FTWIVSGDFDVSWTLRIDTLTAVMLFTVSVISSIIHWYSIGYMAH
DPHIPRFFAYLSLFTFAMLVLVTADNFLQLYFGWEGVGLCSYFLIGFWYDRPSANAAAIK
AFVVNRVGDFGFALGIMAMFFTFNSVNF---DTVFAGAPAVA---GK--TFEFLGAEYDI
LTVTTLLLFIGAMGKSAQVPLHTWLPDAMEGPTPVSALIHAATMVTAGVFLVARCSPLF-
EYAPTTLAIVTVIGAFTAFFAATVGLVQNDIKRVIAYSTCSQLGYMFFACGVGAYAVAIF
HLFTHAFFKALLFLGSGSVIHGFSDEQDMRRMGGVWRKMPVTYAVMWVGSLALA-----G
```

```
IPF-----FA---GYYSKDLIIESAFAASTEVGYLAFYLGVGAALLTAFYSWRLLFMTFH
GPTR------------ANPEVYEH--------------V-HESPKIMTVPLLALAVGALF
AGVAGYDAFVG---DGRAAFW-REAIFLLP-GHDVIE-AAHH-VPLWVKLLPIAIASAGI
ALAWQFYIRRTE---LPG------ALAKTHADLYQFLLNKWYFDELYDAIFVRPAKFLG-
RALWHGGDGRIIDGFGPNGIAQVVIQLARRASLLQTGYLYHYAFAMLIGVAALVTWFTR-
---TGGGAP------
>MSP51668_Alphaproteobacteria_Baikal-deep-G48_o__SHVV01
-------MYAAIVFLPLIGAIVAGLF----GRLIGVRASQLVPCLALSVSGILAAIGLYD
MAA--GQAKIQVRL-FSWIVSGDFDVSWSLRIDALTAVMMFTVSVVSAIIHWYSIGYMAH
DPHRPRFFAYLSLFTFAMLMLVTSDNFLQLYFGWEGVGVCSYFLIGFWFDRPSANAAAIK
AFVVNRVGDCGFALGIMAVFFTFNSIEF---DVVFANAPAVA---GK--SFEFLGAEYDI
LTVTTLLLFIGAMGKSAQVPLHTWLPDAMEGPTPVSALIHAATMVTAGVFLVARCSPLF-
EYAPTTLAVVTVIGAFTAIFAASIGLVQNDIKRVIAYSTCSQLGYMFFAAGVGAYQVAIF
HLMTHAFFKALLFLGSGSVIHGFSDEQDMRKMGGAWRLMPVTYAVMWIGSLALA-----G
IPF-----FA---GYYSKDLIIESAFAAGTDVGYLAFYLGIGAAALTAFYSWRLLFMTFH
GQSR------------ADAEVLSH--------------V-HESPKIMTIPLILLAIGALF
AGIVGYDALVG---EGRVLFW-RDAILVLP-GHDVIE-AAHH-VPWWVKVLPILVAIGGI
ALAWQYYIRRPE---KPG------QLVKVHPDLYQFLLNKWYVDELYEVLFVRPAKFIG-
RVFWIVGDGRIIDGFGPNGVAQAVIALARRASILQSGYVYHYAFAMLIGVAALVTWFFA-
---RTGGGLP-----
>MBM3506331_Alphaproteobacteria_K_Offshore_80m_m2_162_o__CAIVPW01
------MLYTLIVFLPLIGAALAGLF----GRVLGVSGSHVVTTGLVTMSALLSWVAFWF
VAI--QGNAEVVEL-FPWIESGALEAMWSIRVDQLTAVMLVVVNTVAALVHWYSIGYMHD
DPHQPRFFAYLSLFTFAMLTLVTADNFVQMFFGWEGVGLCSYLLIGFWYHKPSANAAAIK
AFIVNRVGDFGFALGICAVFFVFQNVEF---EPVFEAVPRFV---GT--EIEFLGGHFDT
LTVITILLFIGAMGKSAQIGLHTWLPDAMEGPTPVSALIHAATMVTAGVFMVARLSPMF-
EYSEFTLALVTFVGALTAIMAATIGMVQNDIKRVVAYSTCSQLGYMFFACGVSAYQAGIF
HLFTHAFFKALLFLGAGSVIHAFSGEQDMRKMGGVWRMIPLTYALMWIGSLALA-----G
IPL-----FA---GFYSKDVVLEAAWGAHSAVGELAFVLGILAALLTAFYSWRLLFMTFH
GHSR------------ASHEVLHH--------------V-HESPPVMTVPLVVLAIGAVF
AGMLFAPMFVG---ED-HGFW-GESIFVLA-SHTALE-EAHH-VPLWVKLSPTVVSVLGI
ATAWWMYIARPD---LPG------RLAAQHAELYRFFLNKWYFDELYDVLFVRPARAIG-
SFFWKKGDEATIDGFGPNGVASRVLDLARRATQLQTGYVYHYAFVMLIGVVAFVTWYLL-
---VVGM--------
>MDP6874949_Alphaproteobacteria_Arabian9_MAG_8_o_GCA-2731375
-------MYHAIVFLPLLAFLIVGAL----GRILGDKPSQWITCGAVITSAILSLVSFWQ
VAL--GGQPVKVEI-ASWIVSGTFDVSWALRIDALTAVMLVVVNGVSALVHVYSVGYMSH
DPHKPRFMAYLSLFTFAMLMLITSDNFLQMFFGWEGVGLASYLLIGFWYHKPSANAAAIK
AFLVNRIGDFGFGLCIMATFMVFGSVDF---DTVFAAAPDQV---GK--TLHFLNWEWDV
MTTICILLFIGAMGKSAQVPLHTWLPDAMEGPTPVSALIHAATMVTAGVFLVARCSPMF-
EYSPVALTVVTIVGASTAFFAATVGLVQNDIKRVIAYSTCSQLGYMFFALGVGAYPVAIF
HLFTHAFFKALLFLGSGSVIHAMSDEQDMRRMGGIFKYVPQTAIMMWIGSLALA-----G
LPI-----FA---GFYSKDAILESAFAAHTGAGDYAFYLGLAAAVMTAFYSWRLLFMTFN
GTPR------------ASQEVMSH--------------V-HESPPSMLIPLYFLAAGSIL
SGILFSHYFIG---EGHEAFW-GQGIFMLP-DNDILE-ALHH-VPMWVKIAPIAAGIIGI
LVAWQFYIRRTD---LPA------KLAKVHHEAYLFLLNKWYFDELYDLIFVNPAKWLG-
RLLWKGGDGRIIDGFGPDGIAATVMRVARRASALQSGYLYHYAFAMLIGVGLVTWFYT-
---SSGGGH------
```

```
>MBV40747_Rhodospirillaceae_NP113_o_GCA-2731375
-------MYHAIVFLPLLAFLIVGTF----GRVFGDKPSQWITCGAVIASAILSLVAFWQ
VAL--GGNPVKVEI-ASWIVSGTLDVSWALRIDALTAVMLVVVNGVSALVHVYSIGYMSH
DPHKPRFMAYLSLFTFAMLMLITSDNFLQMFFGWEGVGLASYLLIGFWYHKPSANAAAIK
AFLVNRVGDFGFGLGIMATFMVFGSIDF---NTVFAAAPDQV---GK--TFHFLSWEWDV
MTTICILLFIGAMGKSAQVPLHTWLPDAMEGPTPVSALIHAATMVTAGVFLVARCSPMF-
EFSPTALVVVTVVGGSTAFFAATVGLVQNDIKRVIAYSTCSQLGYMFFALGVGAYPVAIF
HLFTHAFFKALLFLGSGSVIHAMSEEQDMRRMGGIFKYVPQTAILMWVGSLALA-----G
LPI-----FA---GFYSKDAILESAFAAHTGAGDYAFYLGLAAAVMTAFYSWRLLFMTFH
GIPR------------ASQAVMSH--------------V-HESPLVMLMPLYFLAAGSIL
SGILFVHYFVG---DGHQAFWRGQAIFMAE-GNDILE-SFHH-VPIWVKVAPIAAGVFGI
FVAWLFYIRNQD---LPA------KLAKLHYPLYLFLLNKWYFDELYDLILVNPAKWLG-
RLLWKGGDGKIIDGFGPDGIAATVMNLARRASALQSGYLYHYAFAMLIGAVVLVTWYYT-
---STGGGH------
>MBT4689853_Rhodospirillaceae_SI039_bin92_o_GCA-2731375
-------MYHAIVFLPLIAFLIVGPF----GRVLGDKPSQWITCGAVITSAVLSIVAFWQ
IALG-GGDTVKVEI-ASWIVSGTFDVSWALRIDALTAVMLVVVNGVSALVHVYSVGYMSH
DPHKPRFMAYLSLFTFAMLMLITSDNFLQMFFGWEGVGLASYLLIGFWYHKPSANAAAIK
AFLVNRVGDFGFGLGIMAIFMVFGSLDF---DTVFAAVPGQV---GN--TFHFLNWEFDI
ITTICILLFIGAMGKSAQVPLHTWLPDAMEGPTPVSALIHAATMVTAGVFLVARCSPMF-
EFSPTALVVVTIVGGFTAFFAATVGLVQNDIKRVIAYSTCSQLGYMFFALGVGAYPVAIF
HLFTHAFFKALLFLGSGSVIHAMSDEQDMRRMGGIFKYVPQTAIMMWIGSLALA-----G
LPI-----FA---GFYSKDAILESAFAAHTGAGMFAFYAGMAAAVMTAFYSWRLLFMTFN
GDPR------------ASQEVMSH--------------V-HESPPSMLIPLYFLAAGSIL
SGMLFVHYFVG---DGHQAFW-GDAIFMGP-ENHILE-AMHH-VPTWVKIGPIAAGIIGI
FVAWLFYIKRKD---IPV------NLAKVHHEVYLFLLNKWYFDELYDLIFVRPAKWLG-
RILWRGGDGRIIDGYGPDGIAAMVMHLARRASALQSGYLYHYAFAMLIGVAVLVTWYYT-
---TGGG-H------
>MBL6951394_Alphaproteobacteria_BS150m-G9_BlackSea_o_GCA-2731375
-------MYHAIVFLPLLAFLIVGSL----GRVLGDRPSQWITCGAVIISALLSLLAFWQ
VAL--GGQAVRVDI-GSWIVSGTFDVSWALRIDQLTAVMLVVVNGVSALVHVYSVGYMSH
DPHKPRFMAYLSLFTFAMLMLVTADNFLQLFFGWEGVGLASYLLIGFWYHRPTANAAAIK
AFLVNRVGDFGLALGIMAIFTVFGSLDF---DTVFAATPGQV---GK--SFNFLSWEWDV
MTTICILLFIGAMGKSAQVPLHTWLPDAMEGPTPVSALIHAATMVTAGVFLVARCSPMF-
EFSPTALVVVTIVGGTTAFFAATVGLVQNDIKRVIAYSTCSQLGYMFFALGVGAYPMAIF
HLFTHAFFKALLFLGSGSVIHAMSDEQDMRRMGGIYKYVPQTTGLMWVGSLALA-----G
FWP-----LA---GYYSKDAILESAFAVHTGAGEYAFWLGMAAAVMTAFYSWRLLFMTFA
GEPR------------ASQEVMRH--------------V-HESPLSMLIPLYFLAAGSIL
SGVLFVHYFVG---EGHVEFW-GQAIFMAP-GNDVME-AVHH-VPTWVKVAPLVAGIVGI
LVAWQFYIRRRD---IPA------GLAKINHEVYEFLLNKWYFDELFELIFVRPAKWLG-
RLLWKTGDGRIIDGFGPDGIAASVLNLARRASALQSGYLYHYAFAMLIGVAALVSWYYI-
---SSGGGH------
>WP_085883634_Oceanibacterium_hippocampi_o_Sneathiellales
-------MYVLIVFLPLAAALIAGFF----GRQLGDRGAQVVTSGAVLTSALLSWVALFQ
VAF--GHAPVTITL-FDWVVSGDFAASWAIRVDALTAVMLVVVNTVSSLVHVYSIGYMSH
DKAKGRFMAYLSLFTFAMLMLVTADNLLQLYFGWEGVGLASYLLIGFWYHKASANAAAIK
AFVVNRVGDFGFALGVVGIFYIFGSIEF---DQIFAAAPEQV---GK--SFVFMGMEVDV
LTTLCLLLFLGAMGKSAQLFLHTWLPDAMEGPTPVSALIHAATMVTAGVFMVARLSPLF-
```

```
EYAPDALTVVTVVGGSTAFFAATVGLVQNDIKRVIAYSTCSQLGYMFFAAGVSAYPVAIF
HLFTHAFFKALLFLCAGSVIHAVSDEQDMRRMGGLWKHIKATYVLMWIGSLALA-----G
IWP-----FA---GYFSKDMVLEAAYAAGTTVGSYAFWLGIAAAIMTAFYSWRLLFMTFH
GAPR------------ASKEVMDH--------------V-HESPKVMLIPLYVLALGSIC
AGFIFAPYMVG---HHYQEFW-GNSILVLE-EHVAMA-EAHD-VAAWVKFLPLVAGVVGI
LIAFQLYIRRPE---MPA------ELARTHQALYQFLLNKWYFDELYDWIFVRPATWLG-
RVLWKGGDGRIIDGLGPDGIAMTVVRVAARAKQIQTGYIYHYAFAILIGIAAFVTLFFY-
---GVD---------
>OUR77105_Alphaproteobacteria_46_93_T64_o_Sneathiellales
-------MYVLIVFLPLLAALIAGFG----ARQLGDRGSQIVTSGAVSASAILSWVALFQ
VAF--GSAPTTIEL-FTWINSGTFDVSWSLRVDSLTAVMLVVVNTVSALVHIYSIGYMSH
DPHKPRFMSYLSLFTFAMLMLVTSDNFLQMYFGWEGVGLASYLLIGFWFNKPSANAASIK
AFVVNRVGDFGFALGIMAIYLVFDSISF---DTVFAAVPEKV---GE--TFNFLGYEVDV
ITTIALLLFLGAMGKSAQLGLHTWLPDAMEGPTPVSALIHAATMVTAGVFMVARCSPIF-
EFSPTALAVVTVVGASTAFFAATVGLVQNDIKRVIAYSTCSQLGYMFFAIGVGAYPVAIF
HLFTHAFFKALLFLGSGAVIHAVSDEQDMRKMGGLGKHIKITYAMMFIGTVALT-----G
VPF-----FA---GFYSKDAIIESAFMAGTDVGNYAFWCGIGAALMTAFYSWRLMFMTFH
GKPR------------ASKEVMSH--------------V-HESPNVMLIPLYVLAAGAIG
AGFVFYEPMVG---HGYKEFW-GNAILLRE-SNTIMT-DFHN-IPSWVFWAPMIAMIAGF
LLAFNMYIRRTD---IPV------QLAKTHSALYQFLLNKWYFDELYDVVFVRPARWIG-
SKLWTIGDGKIIDGLGPDGIAARVLDLAKRASMLQSGYLYHYAFAMMIGVAAFVSFFFL-
---AGGGH-------
>WP_206376079_Sneathiella_chungangensis_o_Sneathiellales
------MMYVLIVFLPLLAALIAGLF----GRSLGDRGSQLITCGAVIISAILSWVALYQ
IAF--QQTTEAVEL-LTWINSGDFDVNWALRIDSLTAVMLVVVNTVSAVVHVYSIGYMSH
DPHKPRFMAYLSLFTFAMLMLVTSDNFVQLYFGWEGVGLASYLLIGFWYKKPAANAAAIK
AFVVNRVGDFGFALGIMGIYFVFNSVSF---DEVFAAAPSQA---GQ--TFEFLGYQVDI
LTTLCLLLFVGAMGKSAQIGLHTWLPDAMEGPTPVSALIHAATMVTAGVFMVARCSPIF-
EYSPEALMVVTVVGATTCFFAATIGLVQNDIKRVIAYSTCSQLGYMFFALGIGAYPAAIF
HLFTHAFFKALLFLSAGSVIHAVSDEQDMRKMGGLWKHIKVTYAMMWIGSLALA-----G
IPL-----FA---GYYSKDLIIESAFAAHTGVGDYAFWAGVAAALLTAFYSGRLIFMTFH
GEPR------------ASKEVMAH--------------V-HESPMVMLIPLGVLAAGAIF
AGMLFAGPMTG---EDFREFW-GASILMLE-NSTAML-DAHH-VPLWVKLLPLVLGVFGI
ALTYQMYIRRTD---IPV------QLAKTHSALYQFLLNKWYFDELYDFLFVRPAKAIG-
RFFWTFGDGKVIDGLGPDGIAGRVLVLARRASQLQSGYMYHYAFAMLLGVAAFVSYFFF-
---AGGH--------
>PJK29752_Minwuia_thermotolerans_o_Minwuiales
-------MYSLIVFAPLLGAILAGFF----GRMIGDRGSMLVTSGLMTLAAVLSWIVFLD
VTF--GHAPTHVEL-LTWITAGEFRVNWALQVDQLTAVMLIVVNTVACLVHWYSIGYMAH
DPHKARFFSYLSLFTFAMLMLVTSDNFVQLYFGWEGVGLASYLLIGFWYHKPSANAAAMK
AFIVNRVGDFGFALGIAGIFLVFEDVTF---AGVFAATPEVA---GQ--TFQFLGYEVDI
LTTICLLLFMGAMGKSAQLPLHTWLPDAMEGPTPVSALIHAATMVTAGVFLVVRCSHMF-
EYSPDALAVVTVVGASTAFFAATIGLVQNDIKRVIAYSTCSQLGYMFFAAGLSAYPVAIF
HLFTHAFFKALLFLGAGSVIHAVADEQDMRKMGGLARHIKKTYIMMWIGSLALAGI---G
IPG-----FAYVFGFYSKDLIVESAFGAGSGVGQFAYIMGVGAAIMTAFYSWRLLFMTFH
GKPR------------APRETMAH--------------I-HDSPPVMMIPLYILAVGAVF
SGLLFIGPLTG---HHWHDFW-GDSILILP-QHGAME-AAHE-VPLWVKLSPLVASLVGI
LIAWVMYVRKTD---LPG------NFANTHRPLYLFLLNKWYFDELYDWIFVRPAKALG-
```

```
RILWKGGDGRIIDGLGPDGISASVLRATRRITMLQTGYVYHYAFAMLIGIAVLVTFYFY-
---AIGV--------
>WP_144258814_Ferrovibrio_terrae_o_Ferrovibrionales
-------MYHLIVFLPLLAAIIAGLF----GRRIGDVASMAVTCVAVIISAVLSWVAFYL
VIS--KGMVVTVKV-LDWIDSGTFSVDWALKIDQLTAVMLVVVNTVSAVVHVYSVGYMSH
DPHRSRFFSYLSLFTFAMLMLITADNFVQLYMGWEGVGLASYLLIGFWFHKPSANAASIK
AFVVNRVGDFGFALGIMALFFATGSVTF---EAVFAAAPDLA---GK--TFHFLWKDWDI
LTVATFLLFLGAMGKSAQLGLHTWLPDAMEGPTPVSALIHAATMVTAGVFMVARCSPLF-
EYAPDTLAFVTVIGASTAFFAATVGLAQNDIKRVIAYSTCSQLGYMFFALGVSAYPAAIF
HLFTHAFFKALLFLGAGSVIHAVDGEQDMRRMGGLWKHIKITYAMMWIGSLALA-----G
FPI-----FA---GYYSKDMILEVAYAA-GGVGQYAYIMGMAAAILTAFYSWRLLFMTFH
GKPR------------ADHHVMEH--------------V-HESPMVMLIPLFVLATGAVL
AGIVFYDGFVG---HHWKEFW-GSSILILE-NHKAMD-EAHH-VPFLVKVGPIIVGVIGI
FIAWIAYIRDTS---LPG------RTAAQHHMLYSFLLNKWYFDELYDRVFVRPTFWLG-
NLLWKGGDGKIIDGLGPDGLAATVVRLARRASILQSGYVYHYAFAMLIGVAVLVTYFFS-
---AMGH--------
>MBM3570281_Alphaproteobacteria_M_DeepCast_65m_m2_129_o_VGDZ01
------MLYGLIVFLPLIAAIVAGFG----GRALGDRGAQVVTCGALIVSALLSGVAFYQ
VAW--KGQPVTVEL-LTWIDSGAFDVMWSLRIDQLTAVMLVVVTWVSAVVHVYSVGYMAH
DPHIPRFMAYLSLFTFAMLMLVTSDNFVQLYFGWEGVGLASYLLIGFWYHKPSANAAAIK
AFVVNRIGDFGFGLGICGVFLVFGTVEF---DPVFQTAPGFA---GK--TFEFLGMKLDM
LTVICLLLFMGAMGKSAQVPLHTWLPDAMEGPTPVSALIHAATMVTAGVFLVARCSPLF-
EYAPDALAVVACIGAFTAFFAATVGLCQNDIKRVIAYSTCSQLGYMFFAAGLSAYAAAIF
HLFTHAFFKALLFLGSGSVIHGMSGEQDMRRMGGLLAPLKITYVLMWIGSLALA-----G
VPL-----FA---GYYSKDAIVETAFMAGGKLGSFAFVMGVAAAFMTAFYSWRLLFMTFH
GQNR------------HPDRVIEH--------------AHHESPAVMLVPLYVLAAGSIF
AGALFAQAFIG---EG-SPIF-GASILVLP-GHDALA-QAHH-APLWVKLAPVAVGLAGI
ALAWWMYVRRPD---LPG------KLAAMHGDAHRFLLNKWYFDELYDALFVRPAKWLG-
HALWKRGDGKVIDGLGPDGIAALTLDIAKRATRLQTGYVYHYAFAMLIGVALLVSWYLY-
---RGGM--------
>WP_109919883_Zavarzinia_compransoris_o_Zavarziniales
-------MIAAIVFLPLLGAVIAGFF----GRLIGARGAQIVTSTLLVVAAVLSWVTFFQ
VTGH-EGGAYKMRL-MTWVLSGAFEVDWAFRVDTLTAVMLVVVTSVSSLVHIYSVGYMAE
DPHIPRFHAYLSLFTFAMLMLVTADNFLQVFFGWEGVGLASYLLIGFWYKKPSANAAAIK
AFVVNRVGDFGFALGIFAIFLVFNSVSF---DTVFAAAAGKA---GQ--TFNFLGYDVDI
MTTICLLLFVGCMGKSAQLGLHTWLPDAMEGPTPVSALIHAATMVTAGVFLVARASPLF-
EFAPSALAVVTVVGATTAFFAATVGLVQNDIKRVIAYSTCSQLGYMFFAAGVGAYEAAVF
HLFTHAFFKALLFLGAGSVIHAMHHEQDMRNMGGTWKYIKFTYAMMWIGNLALA-----G
VPF-----FA---GFYSKDMVLEVAYAAHSGVGTYAFWLGIIAASFTAFYSWRLLFLTFH
GEKR------------WGGHGHGHGHDDHGHGHGHAHTP-HESPLAMTIPLALLAVGAVF
SGFWFYDSFVG---EHREAFW-GGAIFVAE-ANKVIE-HAHH-VPGWVKLLPLIVTAGGI
FVAWVFYILKPK---WPD------QLAKTHREAYAFLLNKWYFDELYDLLFVKGARLFG-
RVFWKQGDGAVIDGLGPNGISARVLDIARRAAKLQSGFVYHYAFAMLIGVALLVTWYVY-
---RMGVTP------
>WP_119778097_Oleomonas_K1W22B-8_o_Zavarziniales
-------MIAAIVFLPLLGAIIVGFF----GRRLGNRGAQVITSGFLVVAAVLSWITFFQ
ATG--HATDEPLKVHLTWMLSGSLEVEWAFRIDALTAVMLVVVNTVSSLVHIYSAGYMEE
DPHIPRFQAYLSLFTFAMLMLVTADNFVQVFFGWEGVGLASYLLIGFWYKKPSANAAAIK
```

```
AFVVNRVGDFGFALGIFAIFFVFNSVSF---DTVFAAAAGET---GK--TFNFLGYDVDI
LTTICLLLFVGCMGKSAQLGLHTWLPDAMEGPTPVSALIHAATMVTAGVFLVARCSPLF-
EHAPVALEVVTIVGASTAFFAATVGLVQNDIKRVIAYSTCSQLGYMFFAAGVGAYEASIF
HLFTHAFFKALLFLGAGSVIHAMHHEQDMRNMGGLWRHIKLTYALMWIGNLALA-----G
IPL-----FA---GYYSKDTILEVAFAAHTGVGTYAYILGTLAAMMTAFYSWRLLFMTFH
GEARWGHGHGHGDHGHGDHGHAHT--------------P-HESPLVMTIPLIVLALGAAA
AGIIFFDDFVG---HHREAFW-GGAIFVAE-TNKVLE-EAHH-IPEWAALLPLVLSVFGI
FLAWVFYIARPK---WPA------ALAQTHKELYAFLLNKWYFDELYDLLFVRPAKAIG-
RLLWKGGDGRIIDGLGPDGISARVLDIARRAVKLQSGFVYHYAFAMLIGVAALVSWYVY-
---RIGVTP------
>MBM3483662_Alphaproteobacteria_K_Offshore_80m_m2_187_o__UBA828
-----MGLAVAIVFLPLIGAAIAGLF----GRVLGQRVAPLVTVAAMGVAAIFGVIVFID
VAW--GNNPGKVEL-FDWITSGTFQAAWAIRVDELTAVMIFVVTVVSFLVHVYSLGYMSH
DEHKPRFFAYLSLFTFFMLALVTADNLLQLYFGWEGVGLASYLLIGFWYKRPSANAAAIK
AFVVNRVGDFGFGLGILAIYLTFGSIEY---DVIFAKAPEMA---NQ--TFSFLSGEFPI
LEVCCVLLFIGAMGKSAQFLLHTWLPDAMEGPTPVSALIHAATMVTAGVFMVARCSPLF-
EYAPTALSVVTFFGATTAIFAATIAITQNDIKRVIAYSTCSQLGYMFFACGVSAYQAGIF
HLTTHAFFKALLFLGAGSVIHAMSNEQDMRKMGGIWKLIPLTYGLMWIGNLALA-----G
IPF-----FS---GFFSKDAILESAFAAHTGLGEYAFWLGFLAAILTAFYSWRLLIMTFH
GKPR------------ADHHTMEH--------------V-HESPWIMTLPLIVLAAGAIV
SGYLLWPII-----DTHHHFF-GESILVLE-HHKALE-EAHH-VPFWVKALPTAAGVIGI
FLAYMLYMAFPS---IPG------RLATRLRPLYLFSFNKWYFDELYEVLFVQPAKYIG-
YGLWKRGDGGLIDGVGPDGIAGAVRQIAQRAVKLQSGYVYHYAFAMLLGVAAIVTWYLF-
---ASMP--------
>MBR72819_Rhodospirillaceae_RS434_o_UBA828
-----MALYVLPIFLPLIGSLISGVL----CRKLSDRQAQVITCLLMGTSAALSIVIFHN
VIL--DGNIQQIKL-FDWISSGSFYVDWSIRIDELTAVMMFVVTSVSFLVHIYSIGYMSH
DAHIPRFFSYLSLFTFFMLALVTSDNFLQLFFGWEGVGLASYLLIGFWFKKPTANAAAIK
AFVVNRVGDFGFALGILAIFLTFDSVSY---DIVFNAAPDIS---GT--NFVFLGHSFDV
LTVICILLFVGAMGKSAQFLLHTWLLDAMEGPTPVSALIHAATMVTAGVFMVARCSPIF-
EFAPIALSVVTFFGATTAIFAATVALTQNDIKRVIAYSTCSQLGYMFFACGVSAYSAGIF
HLTTHAFFKALLFLGSGAVIHAMSDEQDMRKMGGIWKMIPLTYVMMWIGSLALA-----G
VPF-----FS---GFFSKDIILESAFAAETGLGQYAFWLGILAAALTALYSWRLIILTFH
GKPR------------ADEKVMAH--------------V-HESPKTMTIPLVLLALGAIL
SGYFLLPMV-----SEYHDFW-GTSILVLP-EHSALH-DAHH-VAFWVKALPTIAAVIGI
GLAYYLYMFRPD---LPP------KIAKTFRPLYLFSYNKWYFDELYNYCFIKPAKYLG-
YGLWKQGDKALIDGIGPDGMANAAVHIAKRAIKLQSGYVYHYAFAMLVGLAAVITWFLF-
---SRS---------
>MBE04991_Gammaproteobacteria_SAT1367_o_UBA828
-----MALYVLPXFXPLIGAAIAGIF----GRGXSDRQXQIVTCVLMGASAALSIVIFHN
VIL--NGEVQNIQL-FDWISSGSFDVXWSIRIDELTAVMMFVVTFVSFLVHIYSIGYMAH
DKHIPRFFSYXSLFTFFMLALVTSDNXXQLFFGWEGVGLASYLLIGFWFKKPTANAAAIK
AFVVNRIGDFGFALGILAIFLTFDSISY---DVVFAESPNIA---DT--SFQFLGHSFDT
LTVICILLFVGAMGKSAQFLLHTWLPDAMEGPTPVSALIHAATMVTAGVFMVARCSPIF-
EFAPIALAVVTFFGATTAIFAATIALTQXDIKRVIAYSTCSQLGYMFFACGVSAYSAGVF
HLTTHAFFKALLFLGSGAVIHALSDEQDMRKMGGIWKSIPLTYSMMWIGSLALA-----G
VPF-----FS---GFFSKDIILESAFAAETGLGQYAFWLGILAAALTALYSWRLIILTFH
GKPR------------ADEKVMAH--------------V-HEAPATMTXPLILLALGAIF
```

```
SGYFLLPMV-----XEYXDFW-GTSILILP-EHSALH-DAHH-VAFWVKALPTIAGVIGI
GAAYYLYMIRPD---LPS------KIALKIRPLYLFSYNKWYFDELYEFCFIRPAKYLG-
YGLWKQGXKGLIDGIGPDGLANTALAIARRTVKLQSGYLYHYAFAMLLGLAAIVTWFLF-
---SRS---------
>WP_007089417_Thalassospira_xiamenensis_o_Rhodospirillales_A
-------MYPLIVFLPLIAATIAGIIDHHHGVSTSDRISQLITVGGVVTAFIISIFAFVD
VAL--GGNPVTVQL-FTWISSGNFTAEWALRFDTLTCVMLIVVTGVSSMVHIYSIGYMSH
DPDKPRFMAYLSLFTFAMLMLVTADNLIQLFFGWEGVGLASYLLIGFWYSKPSASAAAIK
AFVVNRVGDFGFALGIFAVYVLFDSVQF---DVIFANAEEVA---GT--TLLFLGHEFSA
LEITCFLLFIGAMGKSAQLGLHTWLPDAMEGPTPVSALIHAATMVTAGVFMVARLSPIF-
EYAETTLAIITVVGASTAFFAATIGCVQNDIKRVIAYSTCSQLGYMFFACGVSAYSAGVF
HLMTHGFFKALLFLGAGSVIHAMSDEQDMRKMGGIYKMVPLTFAVMMVGTLAIT-----G
FPG-----LA---GFYSKDLVIESAFAAHTAVGTYAFWAGILAALMTSFYSWRLIFMTFF
GTPR------------ADERTMAH-------------V-HESPAVMTLPLLFLTIGAIF
AGWFAKDWFGGGDYEEMLAFW-NGAIFMAE-GHQALE-HAHH-VPGWVKLAPFVAMVSGF
VIALVMYKLVPS---LPR------QLANTFNGLYRFALNKWYFDELYDKIFVKPAFYLG-
YGFWKSGDGALIDGVGPDGVAAACRNIARRVSALQSGFVYHYAFAMLIGIAALVSYTIW-
---KMG---------
>WP_073954579_Thalassospira_TSL5-1_o_Rhodospirillales_A
-------MYALIVFLPLIAAIIAGLIDHHHGVSTTDRLSQLVTVAGVVIAAALSVAAFIE
VAL--NGNPVTLDL-FTWISSGDFTANWSLRFDTLTCVMLIVVTGVSSMVHIYSIGYMSH
DPDKPRFMAYLSLFTFAMLMLVTADNLIQLFFGWEGVGLASYLLIGFWYSKPSASSAAMK
AFIVNRVGDFGFALGIFAVYVLFGTVQF---DTIFANAEEIS---GS--TLVFLGHDFSA
LNIATFLLFIGAMGKSAQLGLHTWLPDAMEGPTPVSALIHAATMVTAGVFMVARLSPIF-
EYAETTLAIVTVVGATTAFFAATIGCVQNDIKRVIAYSTCSQLGYMFFACGVSAYSAGVF
HLMTHGFFKALLFLSAGSVIHAMSDEQDMRKMGGIWKMVPLTFGVMIIGTLAIT-----G
VPG-----FA---GYYSKDMIIESAFAAHTSVGLYAFWMGVIAAFMTSFYSWRLIFMTFF
GAPR------------ADERTMAH-------------V-HESPAVMTVPLLILAVGAVF
AGWFAVNWFGG-NYEEMLSFW-NGAIFVAE-NHQALE-HSHH-VPEWVKMAPFVAMMLGL
ALAVLMYRLVPS---LPR------MLANTFNGIYRFLLNKWYFDELYDAIFVKPAFALG-
RGFWKAGDGALIDGVGPDGVAAACRNIARRISTLQSGFVYHYAFAMLIGIAALVSYTIW-
---KMG---------
>MBT5765548_Kordiimonadaceae_SI072_bin64_o_Sphingomonadales
-------MYQTIVFLPLLASIIAGLF----GNRIGVRGAQGVTCGALIVSAVLSWVAFFQ
VAMAPGDHATHVQV-LSWIVSGDFNVNWAFQIDTLTAVMLVVVNTVSCLVHIYSVGYMSH
DPHIQRFMSYLSLFTFAMLMLITSDNFVQMFFGWEGVGLASYLLIGFWYKKPSACAAAIK
AFVVNRVGDFGFALGIFAIFMTFGSADF---AVVFASVGDHV---DK--TIHFLGMELDL
LTTICLLLFIGAMGKSAQLGLHTWLPDAMEGPTPVSALIHAATMVTAGVFMVARCSPLF-
EYSPDALAVVAVIGASTAFFAATVGMSQFDIKRVIAYSTCSQLGYMFFALGVSAYGAAVF
HLFTHAFFKALLFLGSGSVIHASSDEQDMRNMGGLRKQIPVTFWMMTIGTLALT-----G
FPF-----LA---GYYSKDMIIESAFAAHSGVGTYAFGLGVAAALMTSFYSWRLVFMTFF
GESR------------ASKEVQDH-------------V-HESPQVMLIPLYVLALGALL
SGFVFKEYFVG---HNYEEFW-AGALFLI--GDNVME-AAHH-VPTWVIASPFIAMVVGF
ATAWYFYVKEPS---MPG------KFTSTFRFAHALSFNKWYFDELYDIIFVKPAMKLG-
MIFWKRGDENIIDRYGPDGVSAAVVRVAQKFKQFQTGYLYHYAFVMLIGLSALVTFFVM-
---GGL---------
>MBT6031505_Kordiimonadaceae_SI072_bin126_o_Sphingomonadales
-------MYQTIVFLPLLASLIAGLF----GNRIGDRGAQIVTCGAMIVSAVLSWVAFYQ
```

```
VALTPGDNSLHVQV-LSWIVSGDFNVNWAFQIDSLTAVMLVVVNTVSCLVHIYSVGYMSH
DPHIPRFMSYLSLFTFAMLMLITSDNFVQMFFGWEGVGLASYLLIGFWYKKPSACAAAIK
AFVVNRVGDFGFALGIFAIFMTFGSADF---AVVFADVGSYT---DK--SIHFLGMELDL
LTTICILLFIGAMGKSAQLGLHTWLPDAMEGPTPVSALIHAATMVTAGVFMVARCSPLF-
EYAPAALEVVCIVGASTAFFAATVGMSQFDIKRVIAYSTCSQLGYMFFALGVGAYGAAIF
HLFTHAFFKALLFLGSGSVIHASSDEQDMRNMGGLRKKIPLTFWMMTIGTLALT-----G
FPF-----LA---GYYSKDMIIESAFAAHSSVGSYAFAMGVAAALMTSFYSWRLVFMTFF
GESR------------ASKEVQDH--------------V-HESPQVMLVPLYVLAFGALF
SGFIFHEYFVG---AHYEDFW-AGALFLV--GDNVME-AAHH-VPMWVVEAPFVAMTVGL
ATAYYFYIRRPE---MPG------KFVETFRFLHAISFNKWYFDELYDILFVRPAMKLG-
YILWKRGDENIIDRYGPDGVSAAVVRVAEKFRSLQTGYLYHYAFAMLIGLSAFVTWFVM-
---GGQ---------
>WP_027135330_Geminicoccus_roseus_o_Geminicoccales
-------MYTILVFAPLVGALIAGLF----GRLIGDRASQVVTCGFMAISAICAISSFFL
NIY---GEPFKVVL-FDWIVVGSFDTQWALRIDGLSATMMLVVSFISFLIHVYSVGYMSH
DASIPRFMSYLSLFTFAMLMLVTSDNLLQLFFGWEGVGVASYLLIGFWYEKPSACAAAMK
AFIVNRVGDFGLILGLAGCYLVFDSIQY---DVIFPQVAEFA---DS--SIVLFSYEFNT
LTIIGVLLFIGAMGKSAQLGLHTWLPDAMEGPTPVSALIHAATMVTAGVFLVARFSPLY-
EYAPTALAMVTLVGASTAFFAATIGTVQNDIKRVIAYSTCSQLGYMFFACGVSAYSAGVF
HLFTHAFFKALLFLSAGSVIHAMSDEQDMRKMGGIWRKIPYTYAMMWIGSLALM-----G
VPF-----FA---GYYSKDAILEAAYASHAPFAGYAFWLGLAAAFLTAFYSGRLLWMTFH
GKPR------------ADHHTMEH--------------V-HESPWVMLVPLFCLAAGAVF
AGWLAKGAL-----DPDAAWW-GGAIFMAH-EPNIME-ELHH-VPALVKFAPLIVGIAGF
GLSWVFYLMMPG---IPG------RLASAARPVYLFLLNKWYFDELYDRIFVRPSKMIG-
SALWRGGDVFVIDGFGPDGIAATTAQLAKQTARLQTGYVYHYAFAMLIGVAALVTWYLF-
---FVS---------
>WP_088559410_Arboricoccus_pini_o_Geminicoccales
------MLHTILVLCPALGALIVGLF----GRYLGDRASQIITCGLMAITALCAWIGVFT
YM---GEPAFKVHW-FTWISVGGLEADWSLRIDSLSTIMMLVVGFISFLIHVYSVGYMSH
DPSKPRFMAYLSLFTFAMLMLVTSDNIIQLFFGWEGVGVASYLLIGFWYTRPSACAAAMK
AFIVNRVGDFALMLGIGALFLVFDSVSY---DVIFNKAASLA---DT--QIVLFGGTFQT
LNIITLLLFIGAMGKSAQIGLHTWLPDAMEGPTPVSALIHAATMVTAGVFLVARFSPVF-
EFAPSTLAFVTFIGATTCIFAATVGMTQFDIKRVIAYSTCSQLGYMFFACGVGAYQAGVF
HLMTHAFFKALLFLGAGSVIHAMSDEQDMRKMGGIWKKIPLTYAMMWIGTLALIGFGIPG
IGG-----FA---GFYSKDAILEAAYASNGSFHLYAFWMGVLAAVLTAFYSGRLIFLSFH
GEPR------------ADHHTMEH--------------V-HESPPIMTVPLMVLGVGAVI
AGFVAFGMV-----EPEGNWW-GGAIFTGP-ENHVLH-AMHE-VPLWVSLAPFFAMAFGL
GLSWVFYIAMPG---TAP------KIAAAVRPLDNFLRNKWYFDELYDRIFVRPALALG-
RGLWHGGDQGIIDRFGPDGVAGTTWGTALRIARLQTGYVYHYAFVMLIGVAVFVSWYLF-
---LQR---------
>TVQ34661_Geminicoccaceae_DLM2.Bin45_o_Geminicoccales
-------MYTAIVFLPLLGAIVAGLF----GRIIGDRAAQIVTCSLLCISALLSGIALLT
VPF---GEPFKVPL-ATWIVSGSFEVDWALRIDALTVVMLVVVTWVSALVHIYSIGYMSH
DASIARFMSYLSLFTFAMLMLVTADNFLQLFFGWEGVGVASYLLIGFWYEKASANNAAMK
AFIVNRVGDFALILGIAAIFLVFDSVDF---DTVFAAAPAFA---EH--QIALFGVTWST
LDLICILLFIGAMGKSAQLGLHTWLPDAMEGPTPVSALIHAATMVTAGVFLIARVSPII-
EYAPVALAVIVLIGATTAFFAATVALTQNDIKRVIAYSTCSQLGYMFFALGVGAYAAAIF
HLFTHAFFKALLFLAAGSVIHGMSNEQDMRRMGGIWRHMKFTYAMMWIGSLALV-----G
```

```
VPF-----FA---GFYSKDLILEAAFAA-GGVGFYAFALGIAAAVMTAFYSGRLLFMTFH
GKPR------------ASEEVMHH--------------V-HESPLVMTIPLGLLAVGAVF
AGYLGLPMV-----ADDHAFW-GGSIYLEM-ERNIIY-LAHY-VPTWVVIAPIVAGFVGL
GLAYVIYIQRTE---LAD------RLAERFRPLYQFSLNKWYFDELYDALFVRPARQAG-
TFLWRRVDAGIIDDYGPNGVAATALEGTKRVVKLQTGYIYHYAFVMLIGVAALVSWFLF-
---VARG--------
>MEO1090806_Pseudomonadota_J06638_39_o__Geminicoccales
-------MYAVLVFAPLVGAIIAGFF----GKVIGDRGAQLVTCSLMLLSAVLAGVALAT
VPY---GAPIQVPL-LTWIHVGSFEVDWALRIDGLTAVMLVVVTWVSTLVHVYSVGYMSH
DETIPRFMSYLSLFTFAMLMLVTADNFVQLFFGWEGVGLASYLLISYWYHKPSANNAAMK
AFIVNRVGDFALLLGLAGIFFVFDSLDF---DTVFAAAPTYA---DT--TIVLFGMALPT
LDVLCILLFIGAMGKSAQLGLHTWLPDAMEGPTPVSALIHAATMVTAGVFLVARVSPMM-
EYAPIALAMVTIVGASTAFFAATIALTQNDIKRVIAYSTCSQLGYMFFACGVGAYSAGIF
HLFTHAFFKALLFLGAGSVIHAMSDEQDMRRMGGIWRQLKFTYAVMWIGSLALV-----G
MPV-----FA---GFYSKDLILEASFAAHSGVGLYAFWLGIAAAMMTAFYSGRLLLMTFH
GKPR------------ASDEVMAH--------------I-HESPLVMTVPLGLLALGAIG
AGYFGLPMV-----NADHAFW-GGGIVVRDLAHNVIE-AAHH-VPLWVKLMPIVVGVAGL
VLAWYMYVAKTD---LPE------RGATRFRWAYELSFNKWYFDEAYEAAFVQPARAVG-
SFLWRRIDAGVIDDFGPNGVAASAMAGTRRVVKLQTGYVYHYAFVMLVGVAALVTWYLF-
---SMWG--------
>WP_150006121_Iodidimonas_muriae_o_Sphingomonadales
-MSQFMSQFHIIVFGPLLGFLIAGLF----GRRLGDRGAMGVTTGLVTLSAVLSAFALKD
VAF--DHNQYRILV-LEWVRSGTLSFDWVLNIDTLTAVMLVVVNTVSALVHWYSIGYMSH
DPHRSRFFAYLSLFTFAMLMLVTSDNLVQMFFGWEGVGLASYLLIGFWFKKPSANAAAIK
AFVVNRVGDFGFSLGIFALFMLTGSVVF---ADIFASLDGVA---DA--RFIFLGYEVHA
LTTIAVLLFIGAMGKSAQLFLHTWLPDAMEGPTPVSALIHAATMVTAGVFLMARMSPIL-
ELAPGAMTLIMVIGGATAFFAASVGLVQNDIKRVIAYSTCSQLGYMFVAIGASAYGAAIF
HLFTHAFFKALLFLGAGSVIHAMSDEQDMRKMGGLSKMIPLTYVMMMVGTLALT-----G
FPY-----TA---GFFSKDMIIEAAFATHNSMGSFGFLMTVVAAFMTSFYSWRLVFMTFH
GTPR------------ADEKVMSH--------------V-HESPAVMWVPLAILAVGAIA
AGFVFSDYFIG---HDRFDFW-AGAL-VTS-HGDVMD-EAHH-IPGWIKWLPTGMMALGF
LLSVLFYVVKPG---IPK------SMAATFPGAYKFLLNKWYFDELYDRIFVRPAFWLG-
RVLWIRGDQKTIDGFGPDGIAATVLAAAKRIRVLQSGYVYHYAFAMLIGIAIVISYFVL-
---TGGGF-------
>Alphaproteobacteria_CM1_5m_P81_o_Puniceispirillales
-------MVYLIVFLPLLGSIIAGFW----GRRLGDKLSLYLTSTLLLISMTLSWITFWQ
LSS--NHVDKVYPL-MEWMNIGDFNVVWSLRHDMLSAVMMIVITTVSAMVHIYSIGYMSH
DNSKPRFMSYLSLFTFAMLMLVTSDNLIQLFFGWEGVGLASYLLIGFWYHKPSAHTAAMK
AFIVNRVGDFGFILGIIAIYSVFGSLYF---DEIFSDVEGLH---KL--SIVMLGFKINV
IELIAILLFIGAMGKSAQIGLHTWLPDAMEGPTPVSALIHAATMVTAGVFLVCRMSPLL-
EFAPFALNVITIVGAMTAIFAATIGLTQFDIKRVIAYSTCSQLGYMFFAAGVSAYPASMF
HLTTHAFFKALLFLGAGSVIHAMSDEQDMRKMGGIWKKIPITYALMWVGSLALA-----G
IPF-----FA---GYYSKDLILEAAWAASSNSGYFAFWLGCLAAFLTAFYSWRVLLLTFH
GNFN------------SSKEVLEH--------------V-HESPIVMLIPLFVLALGAIF
SGWYFYNDFVG---YNWKEFW-GNSIFISE-KNKAFK-LAHY-VPLWVKYLPIFLAILGI
LCAYLFYVLNPN---LPK------ILSKKFSPIYNLFFNKWYFDELYDYLFVKSFIKFG-
NYFWKKGDEGTIDRFGPNGISKLVKNISSKSIIIQSGYIYHYAFAMLIGLVVLITWLLL-
---DYGI--------
```

```
>MDA9734455_SAR116_AH-736-O14_o_Puniceispirillales
-------MVYLIVFLPLLGSIIAGFW----GRRLGDKISLYLTSSLLLISMTLSWITFWQ
LSS--NHVDKVYPL-MEWMNVGDFSIVWSLRHDMLSAVMMIVITTVSAMVHIYSIGYMSH
DNSKPRFMSYLSLFTFAMLMLVTSDNLIQLFFGWEGVGLASYLLIGFWYHKPSAHTAAMK
AFIVNRVGDFGFILGIIAIYSVFGSLYF---DEIFSDVEGLH---KL--SIVILGFKINV
IELIAILLFIGAMGKSAQIGLHTWLPDAMEGPTPVSALIHAATMVTAGVFLVCRMSPLL-
EFAPFALNVITIVGAMTAIFAATIGLTQFDIKRVIAYSTCSQLGYMFFAAGVSAYPASMF
HLTTHAFFKALLFLGAGSVIHAMSDEQDMRKMGGIWKKIPITYALMWVGSLALA-----G
IPF-----FA---GYYSKDLILEAAWAASSNSGYFAFWLGCLAAFLTAFYSWRVLLLTFH
GNFN------------SSKEVLEH--------------V-HESPIVMLIPLFVLALGAIF
SGWYFYNDFVG---YNWKEFW-GNSIFILA-KHKAFK-LAHY-VPLWVKYLPIFLAILGI
LCAYLFYILNPN---LPK------ILSKKLSPIYNLFYNKWYFDELYDYLFVKSFIKFG-
NFFWKKGDEGTIDRFGPNGISNLVKNISSKSIIIQSGYIYHYAFAMLIGLVVLITWLLM-
---DYGI--------
>PPR16856_Alphaproteobacteria_MarineAlpha9_Bin3_o_TMED109
-------MLKLLVFLPLMGSVLSGLF----PKFLGKKGVMWLPTLFSFLSLLFSIILLSQ
AI---TGESYITHL-GYWVNSEALTVSWSLRLDVLSAVMLFVVMLVSSLVHLYSIGYMDH
DEHKSRFFSYLSLFTFAMLMLVTSNNFLQLFFGWEGVGLCSYLLIGFWFKKQSANAAAIK
AFIVNRVGDLGLILGICIIYKLTGSLDF---DKVFEKTSLFL---ES--SILFFNFSIPY
IELACFLLFIGAMGKSAQIGLHTWLADAMEGPTPVSALIHAATMVTAGVFLVARCSPLF-
EYAPVALNFVILVGAITSIFAACIAITQNDIKKIIAYSTCSQLGYMFFACGVSAYSAGMF
HLMTHAFFKALLFLGAGSVIHALSDEQDIRKMGGLAKVLPYTCAAMWIGSLALA-----G
LPP-----FA---GFFSKDIILEAAWGHHSIFGYIAFYLGLIAAFLTAFYSWKILFMVFH
GKPR------------TD---ISK--------------A-HESPLYILIPLLILSIGAII
SGWLGYSMV-----DTHHDFW-LESILILD-NHQALV-NAHH-VPLLVKLSPIIVAILGI
YFAWIFYIKKVD---LPK------KNADKFPKLYNISKNKFWFDEIYDLILTRQLKNIG-
NILWTKGDIRIIDRFGPNGIAYFCSRLASKIRSFQSGYVYHYAFTMFFGLIILFSWQFL-
---IYFGW-------
>MAU28776_Pelagibacterales_MED965 o__TMED109
-------MLKLLVFLPLFGSILSGIF----PSFLGKKGVLWVPTLFCSLSLIFSIILLIQ
AI---SGESYVVNL-GYWIKSESLSVSWAFKFDMLSAIMLFVVMLVSTLVHLYSIGYMDH
DEHKSRFFSYLSLFTFAMLMLVTSNNFLQLFFGWEGVGLCSYLLIGFWFKKQSANAAAIK
AFLVNRVGDLGLILGICVIYKLTGSLDF---KIVFDSASSLV---NS--NFLFFNYSIPY
IELACFLLFIGAMGKSAQIGLHTWLADAMEGPTPVSALIHAATMVTAGVFLVARCSPLF-
EYAPVSLNFVIIVGAVTSIFAACIAITQNDIKKIIAYSTCSQLGYMFFACGVSAYSAGMF
HLMTHAFFKALLFLGAGSIIHALSDEQDIRKMGGLAKALPYTCIAMWIGSLALA-----G
LPP-----FA---GFFSKDIILEAAWGHHSSLGYLAFYLGLIAAFLTALYSWKILFMVFH
GKAR------------SD---ISK--------------A-HESPLYILIPLSILSVGAII
SGWIGYVMV-----DTHHDFW-QGSILILD-SHKALV-NAHH-VPLLVKLSPIIVALLGI
YFAWLFYLKKIH---LPK------IYSEKFPKIYSISKNKFWFDEIYEFALTKPLKSLG-
NILWNKGDVGFIDKFGPNGIAYLCTRLANKIRSFQSGYVYHYAFTMFFGIIIIFSWQFL-
---IYFGLI------
>MCH2678992_Alphaproteobacteria_ALOHA_A2.0_2_o__TMED109
-------MLKLLVFLPLLGSIFSGLF----PAILGRKGSIWIPTLFCFVSLIFSIILLIQ
AI---GGESYIVHL-GYWIKSESLTVSWALKLDMLSAIMLFVVMLVSSLVHLYSIGYMEH
DEHKPRFFSYLSLFTFAMLMLVTSNNFLQLFFGWEGVGLCSYLLIGFWFKKQSANAAAIK
AFLVNRVGDLGLILGICVIYKLTGSLDF---NIVFANVSSFI---NT--SILFFNINIPY
IELACFLLFIGAMGKSAQIGLHTWLADAMEGPTPVSALIHAATMVTAGVFLVARCSPLF-
```

```
EYAPISLNFVILVGAATSIFAACIAITQNDIKKIIAYSTCSQLGYMFFACGVSAYSAGMF
HLMTHAFFKALLFLGAGSVIHALSDEQDIRKMGGLAKTLPYTCIAMWIGSLALA-----G
LPP-----FA---GFFSKDIILEAAWAHNSGFGYIAFYLGLIAAFLTAFYSWKILFMVFH
GKAR------------TD---ISK--------------A-HESPLYILIPLFILSIGAII
SGWIGYRMV-----DTHHDFW-LGSILILD-NHKALA-NAHY-VPLLVKLSPIIVALLGI
YFAWLFYIKKLT---LPK------IYSEKFPRLYRVSKNKFWFDELYEFVFTKQLKKMG-
NMLWTVGDIRIIDRFGPNGIAYFCSRLANKIKLLQSGYVYHYAFTMFFGLIILFSWQFL-
---VYFGLM------
>MBJ85870_Pelagibacterales_SP299_o__TMED109
-------MLKLLVFLPLLGSIFSGLF----PTIIGRKGSIWIPTLFCFVSLFFXIILLMQ
AI---GGESYIVHL-GYWIKSESLNVSWSLKLDMLSAIMLFVVMLVSSLVHLYSIGYMEH
DEHKPRFFSYLSLFTFAMLMLVTSNNFLQLFFGWEGVGLCSYLLIGFWFKKQSANAAAIK
AFLVNRVGDLGLILGICVIYKLTGSLDF---NIVFAXVSSFI---NT--SILFFNLNIPY
IELACFLLFIGAMGKSAQIGLHTWLADAMEGPTPVSALIHAATMVTAGVFLVARCSPLF-
EYAPISLNXVILVGAATSIFAACIAITQNDIKKIIAYSTCSQLGYMFFACGVSAYSAGMF
HLMTHAFFKALLFLGAGSVIHALSDEQDIRKMGGLAKTLPYTCIAMWIGSLALA-----G
LPP-----FA---GFFSKXIILEAAWAXNSXFGYIAFYLGLIAAFLTAFYSWKILFMVFH
GKAR------------TD---ISK--------------A-HESPLYILIPLFILSXGAII
SGWIGYRMV-----DTHHDFW-LGSILILD-NXKAXA-NAHY-VPLLVKLSPIIVALLGI
YFAWLFYIKKLT---LPD------IYSEKFPRLYRVSKNKFWFDELYEFAFTKQLKKMG-
NILWTVGDIRXIDRFGPNGIAYLCXRXANKIKLLQSGYVYHYAFTMFFGLIILFSWQFL-
---VYFGLM------
>HIK87875_Alphaproteobacteria_SZUA-1021_o__TMED109
-------MLTLLVFLPLLGSLLSGLL----PSFVGKKGVIWVPTVLCLLSLILSVILMFQ
AI---SGESYTIIL-GNWINSGNLKIGWALRLDMLSAMMLFVVMLVSTLVHIYSIGYMDH
DPHKARFFSYLSLFTFAMLMLVTSNNFLQLFFGWEGVGLCSYLLIGFWFKKQSANAAAIK
AFLVNRVGDLGLILGIAIIYYLTGSLDY---DVVFNNISSIL---HL--DIKFLNYNIPY
IEITCFLLFIGAMGKSAQIGLHTWLADAMEGPTPVSALIHAATMVTAGVFLVARCSPLF-
EYAPIALNFVIIVGALTSIFAAFIAISQNDIKKIIAYSTCSQLGYMFFACGLSAYSAGMF
HLVTHAFFKALLFLGAGSIIHALTDEKDIRKMGGLARLLPYTCGAMWIGSLALV-----G
LPP-----FA---GFFSKDIILEAAWAHQSSNGYIAFYLGLLAAFLTAFYSWKILFLVFH
GKAR------------SD---ISK--------------A-HESPLYILIPLFILSVGAVI
SGWIGYIMV-----DTHHDFW-HGAILILD-NHKALI-NAHH-VPILVKASPIIVALLGI
LLAWLFYIKRTN---LPL------IFSSKFSFLYNISKNKFWFDELYELLLCHPLKVMG-
NFLWNKGDIGVIDKFGPDGITYICKKFASKIKSFQSGYVYHYAFTMFFGLIALFSWQFL-
---IFLGWI------
>MBF87419_Pelagibacterales_SP94_o_TMED109
-------MLKLLVFLPLLGALLSGIF----PKFIGKKGAMYIPTILCGISFIFSIILLHN
AI---LGESYVINL-ATWINSGYFQINWALKLDMLSSVMLFVVMLVSTLVHLYSIGYMDR
DPHQARFFSYLSLFTFAMLMLVTSNNFIQLFFGWEGVGLCSYLLIGFWFKKQSANAAAIK
AFLVNRVGDLGLILGICLIYFKFNSLDY---NTVFESVTLIK---DN--SFSLLNIEIPI
IELCCFLLFIGAMGKSAQIILHTWLADAMEGPTPVSALIHAATMVTAGVFLVARCSPIF-
EYAPYALGFVTIVGALTSIFAAYVAITQNDIKKIIAYSTCSQLGYMFFACGVSAYSAGMF
HLMTHAFFKALLFLGAGSVIHALSDEQDIRKMGGLYKALPYTCVGMWIGSLALA-----G
LPP-----FA---GFFSKDIILEAAWASHTIIGGFAFYIGLLAALLTAFYSWKILFMVFH
GKSR------------GD---ISS--------------A-HESPLTILIPLTILSIGAIF
SGWLGKPMI-----DTHHDFW-HGTILILD-NHEALK-NAHH-APIWVKISPIIVALIGI
FIAWVLYIKRLN---FTQ------FLCERFSFLYRCSLNKLWFDEVYEYLICQPLKKCG-
```

```
DILWKKGDIGVIDRFGPDGIVNVCRKIAMRIKSIQTGYVYHYAFAMFIGLIAIFSWQFL-
---LLFGWV------
>MBT36108_Rickettsiales_RS508 o__TMED109
-------MLKLLVFLPLLGALLSGIF----PKFIGKKGAMYIPTILCGISFIFSIILLHN
AI---LGESYVINL-ATWINSGYFQINWALKLDMLSSVMLFVVMLVSTLVHLYSIGYMDH
DPHQARFFSYLSLFTFAMLMLVTSNNFIQLFFGWEGVGLCSYLLIGFWFKKQSANAAAIK
AFLVNRVGDLGLILGICLIYFKFNSLDY---NTVFESVTLIK---DN--SFXLLNIEIPI
IELCCFLLFIGAMGKSAQIILHTWLADAMEGPTPVSALIHAATMVTAGVFLVARCSPIF-
EYAPYALGFVTIVGALTSIFAAYVAITQNDIKKIIAYSTCSQLGYMFFACGVSAYSAGMF
HLMTHAFFKALLFLGAGSVIHSLSDEQDIRKMGGLYKALPYTCVGMWIGSLALA-----G
LPP-----FA---GFFSKDIILEAAWASHTIIGGFAFYIGLLAALLTAFYSWKILFMVFH
GKSR------------GD---ISG--------------A-HESPLTILIPLTILSIGAIF
SGWLGKPMI-----DTHHDFW-HGTILILD-NHEALK-NAHH-APIWVKISPIIVALIGI
FIAWVXYIKRLN---FTQ------VLCERFSFLYRCSLNKLWFDEVYEYLICQPLKKCG-
DILWKKGDIGVIDRFGPDGIVNVCRKIAMRIKSIQTGYVYHYAFAMFIGLIAIFSWQFL-
---LLFGWV------
>MBT34931_Rickettsiales_RS508 o__TMED109
-------MLKLLVFLPLLGALLSGIF----PKFIGKKGAMYIPTILCGISFIFSIILLHN
AI---LGESYVINL-ATWINSGYFQINWALKLDMLSSVMLFXVMXVSTLVHLYSIGYMDH
DPHKARFFSYLSLFTFAMLMLVTSNNFIQLFFGWEGVGLCSYLLIGFWFKKQSANAAAIK
AFLVNRVGDLGLILGICLIYFKFNSLDY---NTVFESVTLIK---DN--SFSLLNIEIPI
IELCCFLLFIGAMGKSAQIILHTWLADAMEGPTPVSALIHAATMVTAGVFLVARCSPIF-
EYAPYALGLVTIVGALTSIFAAYVAITQNDIKKIIAYSTCSQLGYMFFACGVSAYSAGMF
HLMTHAFFKALLFLGAGSVIHSLSDEQDIRKMGGLYKALPYTCVGMWIGSLALA-----G
LPP-----FA---GFFSKDIILEAAWASHTIIGGFAFYIGLLAALLTAFYSWKILFMVFH
GKSR------------GD---ISG--------------A-HESPLTILIPLTILSIGAIF
SGWLGKPMI-----DTHHDFW-HGAILILD-NHEALK-NAHH-APIWVKISPIIVALIGI
FIAWVLYIKRLN---FTQ------VLCERFSFLYRCSLNKLWFDEVYEYLICQPLKKCG-
DILWKKGDIGVIDRFGPDGIVNVCRKIAMRIKSIQTGYVYHYAFAMFIGLIAIFSWQFL-
---LLFGWV------
>PPR31275_Alphaproteobacteria_MarineAlpha9_Bin2_o_TMED109
-------MLSMAVFLPLIGSILSGLF----PKFLGKKGVVWVPSILCVFSFIISTFLFHD
AI---SGESYEFHI-FNWIQSSSINISWSLKLDVLSGIMLFIVSLVSSIIHIYSIGYMSH
DPGQARFFSYLSLFTFSMLILVSSSNFVQLFLGWEGVGLCSYLLIGFWYKKNEANNAAIK
AFIVNRIGDLGLILGICAIYKTFGSLVY---SEVFFRAYSFA---PV--DILILGQPIPV
LELIGFMLFIGAMGKSAQLGLHTWLADAMEGPTPVSALIHAATMVTAGVFLVVRCSPIY-
EYTTYALGFITIIGALTAIFAASIAITQNDIKKIIAYSTCSQLGYMFFACGVSAYSAGIF
HLMTHAFFKALLFLGAGSVIHALS-EQNIKKMGGLSKHLPFTCIAMWIGCLALA-----G
IPP-----FS---GFFSKDIILEAAWSSHTLLGSISFYLGIVTAFLTAFYSWRIIFLVFH
VSTK------------NN---YEG--------------V-RESPMTMLVPIGLLSFGAIF
SGWLGLDLV-----DTTKDFW-VDSITVIA-RHDTLQ-YTHH-VPLLIKLLPAIIAVSGI
ALAWIIYVKFPK---LAT------IFSIKLRLLYVISYNKYWFDEIYEKYICDPIKSLG-
KFLWEKGDIQIIDKFGPDGISNFCINFASRLRKLQSGYVYHYAFSMFFGLIIFISWHFL-
---VFLGVY------
>PPR29499_Alphaproteobacteria_bacterium_MarineAlpha9_Bin1_o_TMED109
-------MLSMAVFLPLIGSILSGLF----PKFLGKKGVVWVPSILCVFSFIISTFLFHD
AI---SGESYEFHI-FNWIQSSSINISWSLKLDVLSGIMLFIVSLVSSIIHIYSIGYMSH
DPGQARFFSYLSLFTFSMLILVSSSNFVQLFLGWEGVGLCSYLLIGFWYKKNEANNAAIK
```

```
AFIVNRIGDLGLILGICAIYKTFGSLVY---SEVFFRAYSFA---PV--DILILGQPIPV
LELIGFMLFIGAMGKSAQLGLHTWLADAMEGPTPVSALIHAATMVTAGVFLVVRCSPIY-
EYTTYALGFITIIGALTAIFAASIAITQNDIKKIIAYSTCSQLGYMFFACGVSAYSAGIF
HLMTHAFFKALLFLGAGSVIHALS-EQNIKNMGGLSKHLPFTCIAMWIGCLALA-----G
IPP-----FA---GFFSKDIILEAAWSSHTLLGSISFYLGIVTAFLTAFYSWRIIFLVFH
VSTK------------NN---YEG--------------V-RESPMTMLVPIGLLSFGAIF
SGWLGLDLV-----DTTKDFW-VDSITVIA-RHGTLQ-YTHH-VPLLIKLLPTIIAVSGI
ALAWIIYVKFPK---LAT------IFSIKLRLLYVISYNKYWFDEIYEKYICDPIKSLG-
KFLWEKGDIQIIDKFGPDGISNFCINFASRLRKLQSGYVYHYAFSMFFGLIIFISWHFL-
---VFLGVY------
>MBE53038_Pelagibacterales_SAT136 o__TMED109
-------MLTTAVFLPLIGSIFSGLL----PKTLGKNGAVWIPTLSCIISFILSIFLLLS
AL---SGDTYEVLL-LNWIQSSLLDISWSLRLDVLSAIMIFTVSLXSSIVHLYSVGYMSN
DPNQERFFSYLSLFTFSMLILVAXSNFVQLFLGWEGVGLCSYLLIGFWYKKDKANKAAIK
AFIVNRVGDLGLVLGLCVLYKTFGSLDY---SIVFSQAYDYR---LV--DISILGYSIPS
LELIGILLFIGAMGKSAQLGLHTWLADAMEGPTPVSALIHAATMVTAGVFLIARCSPLY-
EHTTYALGFIIIIGALTAIFAASIAIAQDDIKKVIAYSTCSQLGYMFFACGVSAYSAGIF
HLMTHAFFKALLFLAAGSVIHALSGEQNLKKMGGLSKYLPLTCASMWIGSFALA-----G
IPP-----FA---GFFSKDIILEAAWGSHSVLGSIAFYLGIAAAFMTAFYSWRVIFLVFX
VKTX------------NX---YKN--------------I-HESPFTMLIPIGLLSLGAIF
SGWIGLDMI-----SXSXDFW-MGSIQVIS-GHDAXA-NAXH-APIYIKLLPIVIATSGI
ILAWVFYIKLPH---MPN------VISDKMRFMYLILYNKYWFDEFYDRXICFPLKRFG-
QFLWEKCDIKVIDKYGPDGISNYCIILASRIRKFQSGYVYHYAFSMFFGLIIFISWHFI-
---VYLGVF------
>AIL65955_Rickettsiales_Ac37b_o_Rickettsiales
----MKNEIVFLVFIPLITSVISGLF----MQRVNAALAQFLTCIGMAISSFLSLIIFYN
IIIK-GYNYNTVEL-LRWIEINELVVNWALKVDALTAVMLVVVTVVSFLVHVYSVGYMHD
DQHKPRFMSYLSLFTFFMLMLVTSDNFLQLFLGWEGVGLCSYLLIGFWYQKNSANLAAMK
AFIVNRVGDFGFILGIFSIYLIFNSLQF---DQIFKLVNEYS---YK--TFNILGYNISA
IEISCMLLFIGCMGKSAQLGLHIWLPDAMEGPTPVSALIHAATMVTAGVFLLVRCSWLF-
QYAPIVLYIITVIGALTCIFAATIALTQNDIKKIIAYSTCSQLGYMFFACGVSAYNVALF
HLATHAFFKALLFLGAGSVIHAMSGEQDINKMGGIWRKIPFTYLMMWIGSLALA-----G
IYP-----FA---GYYSKDMIIEAAYLSPSESGRFAYWIGISAAFCTAFYSWRLLIKVFH
GKFK------------NEPTVWKR--------------V-HESPLFMTIPLFILSLGAIF
SGIIGEKLHIV---DNTNKFW-HGAIEIFH-SH------SHH-IPVLYKFMPMGVGILGI
VIAVIYYVVFKV---LPD------ITKSTISPLYNLSYYKYYWDEIYEYIVIKPIRCLS-
VVLWKKIDNVIIDGLGPNGVVKIINLLSTKVTKMQTGYLYHYAYIMLGAVMVLTSWYIF-
---NIIN--------
>AEI88434_Midichloria_mitochondrii_o_Rickettsiales
MNMYLHNLSLSVVFLPLLASIPVGLN----TNKVKPVIAQIITISFVSIAAIFSWIIFYY
TCF--DGQIIHLKL-INWLNLGELKADWSIYIDPLTAIMFLVVNTVSTLVHIYSVSYMSH
DPNQPRFFSYLSLFTFFMLVLVSADNFAQLFVGWEGVGLSSYLLIGFWFHKKSAYSAAMK
AFLVNRVGDIGLAIGIFLIAMKFGSVEY---ATVFSKTKYLS---EE--TINFLSVDTRL
LTVICIALFIGCMGKSAQLGLHTWLPDAMEGPTPVSALIHAATMVTAGVFLLARCSYLF-
EYSEIALDLVTIVGAATCLFAATIATAQNDIKKIIAYSTCSQLGYMFFACGVSAYSAGIF
HLLTHAFFKALLFLGAGSIIHALADEQNIKKMGGIWKKLPYTHALMWIGSIALA-----G
IPP-----LS---GFFSKDLILEAAYASDSRFGHLAYWLGVAAATLTAFYSCRLLFLVFY
APTN------------ADKKIFTE--------------I-HEAPQSMMIPLMVLGIGSLI
```

```
CGYIGYHIIDI---SN-LSFW-SGAIFVLQ-EHDSID-KIDT-IGHWVAQIPLLAALFAM
LLAYYSYILKPI---VSS------WSYKNFRNINSFLEKKWYFDEVYEFILIKPVKLLG-
TSLWRFVDVGFIDGI-PNYSARAVARFAKVACLLQTGYIYHYAMAMLIGVMILVYWYLPW
---------------
>Aquarickettsia_rohweri_Acer_BE_o_Rickettsiales
-MLSIENLAIYIVFLPLMASIIVGLN----TNNISVKTSQILTSFSVFISAIFSIIVFFY
VIH--GHNIIHIKL-LKWLDVSFFKAYWSIYIDSLTAVMLCVVNIVSFLVHLYSVGYMSH
DNHKQRFMAYLSLFTFFMLVLVTSDNLLQLFVGWEGVGLSSFLLIGFWFKKKSAANAAIK
AFLVNRVGDIGLALSIILIALTFKTIEY---QQIFIQLDRSY---EE--CFSLFGIKFSV
IDSICLLMFIGCMGKSAQIGLQTWLPDAMEGPTPVSALIHAATMVTAGVFLVARCSFIF-
EQSIIALNMITIIGAITCLFAASIAIVQNDIKKVIAYSTCSQLGYMFFACGVSAYSVAIF
HLMTHAFFKALLFLGAGSVIHAINDEQDIRKMGGLYKLLPKSYTLMWIGSLALA-----G
IYP-----FA---GYFSKDLILESAYLSNTSYGDFAFCMGIIAALLTAFYSWRLIILVFH
KKTK------------LEKNIVQN--------------I-HDAPKSMFIPMIVLAIGAIF
SGYVGEYIFEMT--NSSSDFW-NHSLAVTQ-KSNG---ELYM-LPFLVKHMPIIAGLFGI
LIAYIIYANTKK---IAC------FISEKFNFIYKILLNKYYFDEIYNFLFVDFIKKIS-
NIFWKIFDNSIIDGI-PNILARCVYNVSRISRGLQTGVINHYSLIMFIGLLTIMIFIIFY
---------------
>MAT32988_Alphaproteobacterium_MED-G10_o_TMED127
f__TMED127;g__MED-G10;s__MED-G10
-------MFFLSIFLPLLSYFFCVFF----KDYLASKKIAYMSCVFMSLSTISSILSFYF
FKL---NSESFIIF-TNWINSGSFSVDWAIQLNLLSASMILMVNFVSTLIHIYSVGYMKK
DPKISVFMGYLGLFTFFMLLLVSSNNLLQLFLGWEGVGLTSYLLIGFWNYKDSANSAAIK
AFVVNRVGDFGLLMALFTIFVVFGTLNI---NEIMILVNSYS---DT--YFDFIGLKVHS
ITLISLLLFIGCMGKSAQFGFHTWLPDAMEGPTPVSALIHAATMVTAGVYLLILMSPLI-
EASTFAGDFILIVGCLTCIFAASVAVFQNDIKKIIAYSTCSQLGYMFMAVGASAYTSAYF
HLLSHAFFKALLFLGAGSVIHSMSDEQNIKNMGGLYNKIPLTYLTMIIGSLSLV-----G
IPF-----FS---GYFSKDLILEILYLGDSSLSFYAFLVGITVVLLTTLYSMRLIIYVFH
RKSM------------ADEKVIAH--------------I-HESPVIMMIPLAILSVFAIF
FGMFCHSMFIG---IDYLNLW-SDVLYVNK-VTDQNY-IYEN-VPTFFKKLPLIVILIGA
IISYITYFSLVS---FLP------KIKSRLNMIYKFFKNKWYIDEIYQKTFVFLAFYLG-
NGFWKSVDKDLIDNLGPNGLSRFVSSFSGVVAKVQSGFLYHYVLSIIIGLTLLISLYTY-
---IF----------
>fig|2026788.20.peg.1116 MED999 f__TMED127;g__MED-G10;s__MED-G10
-------MFFLSIFLPLLSYFFCVFF----KDYLASKKIAYMSCVFMSLSTISSILSFYF
FKL---NSESFIIF-TNWINSGSFSVDWAIQLNLLSASMILMVNFVSTLIHIYSVGYMKK
DPKISVFMGYLGLFTFFMLLLVSSNNLLQLFLGWEGVGLTSYLLIGFWNYKDSANSAAIK
AFVVNRVGDFGLLMALFTIFVVFGTLNI---NEIMILVNSYS---DT--YFDFIGLKVHS
ITLISLLLFIGCMGKSAQFGFHTWLPDAMEGPTPVSALIHAATMVTAGVYLLILMSPLI-
EASTFAGDFILIVGCLTCIFAASVAVFQNDIKKIIAYSTCSQLGYMFMAVGXSAYTSAYF
HLLSHAFFKALLFLGAGSVIHSMSDEQNIKNMGGLYNKIPLTYLTMIIGSLSLV-----G
IPF-----FS---GYFSKDLILEILYLGDSSLSFYAFLVGITVVLLTTLYSMRLIIYVFH
RKSM------------ADEKVIAH--------------I-HESPVIMMIPLAILSVFAIF
FGMFCHSMFIG---IDYLNLW-SDVLYVNK-VTDQNY-IYEN-VPTFFKKLPLIVILIGA
IISYITYFSLVS---FLP------KIKSRLNMIYKFFKNKWYIDEIYQKTFVFLAFYLG-
NGFWKSVDKDLIDNLGPNGLSRFVSSFSGVVAKVQSGFLYHYVLSIIIGLTLLISLYTY-
---IF----------
>MAR63623_Rickettsiales_NAT268_f__TMED127;g__MED-G10;s__MED-G10
```

```
-------MFFLSIFLPLLSYFFCVFF----KDYLTSKKIAFMSCVFMSLSTISSILSFYF
FKL---NSESFIIF-TNWINSGSFSVDWAIQLNLLSASMILMVNFVSTLIHIYSVGYMKK
DPKISVFMGYLGLFTFFMLLLVSSNNLLQLFLGWEGVGLTSYLLIGFWNYKDSANSAAIK
AFVVNRVGDFGLLMALFTIFIVFGTLNI---NEIMILVNSYS---DT--YFDFIGLKVHS
ITLISLLLFIGCMGKSAQFGFHTWLPDAMEGPTPVSALIHAATMVTAGVYLLILMSPLI-
EASTFAGDFILIVGCLTCIFAASVAVFQNDIKKIIAYSTCSQLGYMFMAVGASAYTSAYF
HLLSHAFFKALLFLGAGSVIHSMSDEQNIKNMGGLYNKIPLTYLTMIIGSFSLI-----G
IPF-----FS---GYFSKDLILEILYLGDSSLSFYAFLVGITVVLLTTLYSMRLIIYVFH
RKSM------------ADEKVIAH--------------I-HESPVIMMIPLAILSVFAIF
FGMFCHSMFIG---IDYLNLW-SDVLYVNK-VTDQNY-IYEN-VPTFFKKLPLIVILIGA
IISYITYFSLVS---FLP------KIKSRLNIIYKFFKNKWYIDEIYQKTFVFLAFYLG-
NGFWKSVDKDLIDNLGPNGLSRFVSSFSGVVAKVQSGFLYHYVLSIIIGLTLLISLYTY-
---IF----------
>fig|2026788.340.peg.1297_139573_S174_metabat_bins_2.tsv.090_sub_f__TMED127;g__MED-G10;s__MED-G10
--------------------------------------------MSLSTISSIVSFYI
FKL---NNESFIIF-TNWINSGSFSVDWAIQLNLLSASMILMVNFVSTLIHIYSVGYMKK
DPKISVFMGYLGLFTFFMLLLVSSNNLLQLFLGWEGVGLTSYLLIGFWNYKDSANSAAIK
AFVVNRVGDFGLLMALFTIFVVFGTLNI---NEIMILVNSYS---DT--YFDFIGLKVHS
ITLISLLLFIGCMGKSAQFGFHTWLPDAMEGPTPVSALIHAATMVTAGVYLLILMSPLI-
EASTFAGDFILIVGCLTCIFAASVAIFQNDIKKIIAYSTCSQLGYMFMAVGASAYTSAYF
HLLSHAFFKALLFLGAGSVIHSMSDEQNIKNMGGLYNKIPLTYLTMIVGSLSLI-----G
IPF-----FS---GYFSKDLILEILYLGDSSLSFYAFLIGITAVLLTTLYSTRLIIYVFH
RKSM------------ADEKVIAH--------------I-HESPVIMMIPLVILSVFAIF
FGMFCHSMFIG---IDYLNLW-SDVLYVNK-VTDQNY-IYEN-VPTFFKKLPLIVILIGA
IISYITYFSLVS---FLP------KIKSRLNMIYKFFKNKWYIDEIYQKTFVFLAFYLG-
NGFWKSVDKDLIDNLGPNGLSRFVSSFSGVVAKVQSGFLYHYVLSIIIGLTLLISLYTY-
---IF----------
>MBD22427_Alphaproteobacteria_SAT166_f__TMED127;g__MED-G10;s__MED-G10
MGLKDKKLFFLSIFLPLLSYCVCVFF----KNYLNSKKIAFFSCGLLIMSTILSASSFYF
LKI---VGESTILI-SNWISSGSFSADWSLNFNLLSVSMVLMVNFVSTLIHIYSIGYMKK
DPKRSIFMGYLGLFTFFMLLLVSSNNLLQLFLGWEGVGLTSYLLIGFWNYKDSANNAAIK
AFVVNRIGDFGLLIALFTIFVVFGTLNI---NEIMILVNSYA---DT--YFDFLGIEIHS
ITLISVLLFIGCMGKSAQFGLHTWLPDAMEGPTPVSALIHAATMVTAGVFLLILMSPLI-
EVSMFASNLILIVGSLTCIFAASVAIFQNDIKKIIAYSTCSQLGYMFMAIGASAYSVAYF
HLISHAFFKALLFLGAGSVIHSMSDEQNIKNMGGLYNKIPLTYLTMIIGSLSLI-----G
IPF-----FS---GYFSKDLILETLYLRDSQLSSYAFFIGIIAVLFTTLYSLRLIIYVFH
RKSM------------ADEKVVAH--------------I-HESPLIMTFPLIILSIFAIF
FGMFSHSLFIG---SDYLNTW-SEIIFVNK-TLNENY-IYEN-MPIFYKKLPLIIIIFGV
TVSYLTYFSLDS---FIP------LIKKRLKIIHNFFKNKWYIDEIYKNTFISGAFYLG-
NGFWKSVDRDLIDNLGPNGMSKVIRSISSLVSKMQSGFLYHYVLSIIIGLTLLISLYTY-
---IF----------
>fig|91750.200.peg.1084 AG-901-J22 f__TMED127;g__MED-G10;s__MED-G10
-------MFFLSIFIPLISYLFCVIF----KSYLSGKQISLISCGLLSIATTLSLISFYF
FHI---NGDSVVLI-SNWISSGSFSVDWSINFNLLSCSMVLMVNFVSTLIHIYSVGYMNK
DPKRSTFMGYLGLFTFFMLLLVSSNNLLQLFLGWEGVGLTSYLLIGFWNYKNSANDAAIK
AFVVNRIGDFGLLLAIFTIFVVFGTLNI---NEIMILVNSYS---DS--YFNFLGIKIHS
ITLISVLLFIGCMGKSAQFGLHTWLPDAMEGPTPVSALIHAATMVTAGVFLLILMSPLI-
```

EVSEFASKLILIVGSLTCIFAASVAIFQNDIKKIIAYSTCSQLGYMFMAVGASAYSLAYF
HLISHAFFKALLFLGAGSVIHSMSDEQNIKNMGGLYNKIPLTYLTMIIGSLSLI-----G
VPF-----FS---GYFSKDLILETLFLRDSELSSYAFLTGLLAVLFTTLYSLRLIIYVFH
RKSM------------ADEKVVAH--------------I-HESPFIMTFPLIVLSIFAIF
FGMFSNSYFIG---SDYLSVW-SEIIFVDK-SFNENF-IYEN-MPIFYKKLPLIIIICGI
IISYLIYFSFDK---FLP------VIKNRLRIVHNFFKNKWYIDEIYKKTFVFLAFYLG-
KGFWKSVDRDLIDNLGPNGMSKFVSSIGLIVSKMQSGYLYHYVLSIIIGLTLLISLYTY-
---IF----------
>fig|91750.1103.peg.485_ERR599106_bin.64_MetaBAT_v2.12.1_MAG_f__TMED127;
g__MED-G10;s__MED-G10
-------MFFLSIFIPLISYLFCVIF----KSYLSGKQISLISCSLLSIATILSLISFYF
FHI---NGDSVVLI-SNWISSGSFSVDWSLNFNLLSCSMVLMVNFVSTLIHIYSVGYMNK
DPKRSTFMGYLGLFTFFMLLLVSSNNLLQLFLGWEGVGLTSYLLIGFWNYKNSANDAAIK
AFVVNRIGDFGLLLAIFTIFVVFGTLNI---NEIMILVNSYS---DS--YFNFVGIKIHS
ITLISVLLFIGCMGKSAQFGLHTWLPDAMEGPTPVSALIHAATMVTAGVFLLILMSPLI-
EVSEFASKLILIVGSLTCIFAASVAIFQNDIKKIIAYSTCSQLGYMFMAVGASAYSLAYF
HLISHAFFKALLFLGAGSVIHSMSDEQNIKNMGGLYNKIPLTYLTMIIGSLSLI-----G
VPF-----FS---GYFSKDLILETLFLRDSELSSYAFLTGLLAVLFTTLYSLRLIIYVFH
RKSM------------ANEKVVAH--------------I-HESPFIMTFPLIVLSIFAIF
FGMFSNSYFIG---SDYLSIW-SEIIFVDK-SFNENF-IYEN-MPIFYKKLPLIIIIFGM
IISYLIYFSFNN---FIS------VIKNRLSIVHNFFKNKWYIDEIYQKTFVFVAFYLG-
KGFWKSVDRDLIDNLGPNGMSKFVSSIGLIVSKMQSGYLYHYVLSIIIGLTLLISLYTY-
---IF----------
>fig|91750.494.peg.823_AG-891-P19_f__TMED127;g__MED-G10;s__MED-G10
-------MFFLSIFIPLISYFFCVIF----KSYLSGKQISLISCSLLSIATILSLISFYF
FHI---NGDSVVLI-SNWISSGSFSVDWSLNFNLLSCSMVLMVNFVSTLIHIYSVGYMNK
DPKRSTFMGYLGLFTFFMLLLVSSNNLLQLFLGWEGVGLTSYLLIGFWNYKNSANDAAIK
AFVVNRIGDFGLLLAIFTIFVVFGTLNI---NEIMILVNSYS---DS--YFNFVGIKIHS
ITLISVLLFIGCMGKSAQFGLHTWLPDAMEGPTPVSALIHAATMVTAGVFLLILMSPLI-
EVSEFASKLILIVGSLTCIFAASVAIFQNDIKKIIAYSTCSQLGYMFMAVGASAYSLAYF
HLISHAFFKALLFLGAGSVIHSMSDEQNIKNMGGLYNKIPLTYLTMIIGSLSLI-----G
VPF-----FS---GYFSKDLILETLFLRDSELSSYAFLTGLLAVLFTTLYSLRLIIYVFH
RKSM------------ANEKVVAH--------------I-HESPFIMTFPLIVLSIFAIF
FGMFSNSYFIG---SDYLSIW-SEIIFVDK-SFNENF-IYEN-MPIFYKKLPLIIIIFGM
IISYLIYFSFNN---FIP------VIKNRLSIVHNFFKNKWYIDEIYQKTFVFVAFYLG-
KGFWKSVDRDLIDNLGPNGMSKFVSSIGFIVSKMQSGYLYHYVLSIIIGLTLLISLYTY-
---IF----------
>MBC10688_Rickettsiales_SP106_f__TMED127;g__GCA-2711515;s__GCA-2711515
-------MFFLSIFLPLLSFILCISF----KDYLKSKKLNFLSCSLLVISTICSFISFYL
LTK---NGETTITL-GEWITSGSFSVDWSIQHNLLSGSMILMVNFVSTLIHIYSVGYMEK
DPKNVLFMGYIGLFTFFMLFLVSSSNLVQLFLGWEGVGLTSYLLIGFWNYKDSANEAAFK
AFXANRVGXFGLLLALFTIFIVFGTLNI---SEIMILVNAHT---ET--YFEFIGLKIHA
ITLICILLFVGCMGKSAQFGLHVWLPDAMEGPTPVSALIHAATMVTAGVFLLILMSPLL-
QASEISREFILIIGCLTCLFASSVAIFQNDIKRIIAYSTCSQLGYMFMAIGVSAYSAAYF
HLLSHAFFKALLFLGAGSVIHSMSDEQDIKKMGNLFYKIPVTYFTMLIGSFSLL-----G
LPF-----FS---GYYSKDLIIEFLYLSESDLSYYFFFGSVLGVLFTSIYSIRLIIYVFH
GKNN------------ADEKVVAH--------------I-HESPMIMIIPLVILSFFAVF
FGMLSSNFFIS---SEYLESW-SKIMYINK-DLDKFF-IYEN-VPIFFKKLPLMMIILGT

```
LICLILYFSLQNS--VLP------FLRSNFSFIWSFFKNKWYVDELYEKVILKPIMYFG-
RGFWKSIDIDLIDNLGPNGVSRVVKSFSVMVSRLQSGYLYHYVLSVVIGLTLLVSLYTY-
---IF----------
>fig|91750.248.peg.938_AG-404-M02_f__TMED127;g__GCA-2711515;s__GCA-2711515
-------MFFLSIFLPLLSFFICVSL----KDFFKSQQLSFISCSLLFMASVCSLLSLIG
LKI---EEEKVIIL-GNWISSGSLDLNWAIQLNLLTGSMVLMVNFVSTLIHIYSVGYMNK
DPKNILFMGYIGLFTFFMLFLVSSSNLVQLFLGWEGVGLTSYLLIGFWNYKDSANKAALK
AFIANRVGDFGLLLALFSIFIIFGTLNI---NEIMVLVNSHV---DT--YFEFLGARIHA
ITLICILLFVGCMGKSAQFGLHVWLPDAMEGPTPVSALIHAATMVTAGVFLLIIMSPLL-
EASSFSRDFILIIGSITCLFASSVAIFQNDIKRIIAYSTCSQLGYMFMAIGASAYSAAYF
HLLSHAFFKALLFLGAGCVIHSMSDEQNIKKMGGLFNKIPLTYFTMLIGSFSLL-----G
LPF-----FS---GYYSKDIIIELLYLSESKFSFYFFLASVLGVLFTSIYSVRLLIYVFH
GKSN------------SDEKVIAH--------------I-HESPMIMIIPLVILSSFAVF
FGMFSQSFFFG---SDHLDNW-SQLMHINK-KVDEFY-IYEN-VPTLFRKLPLIMIVIGS
SMCFLLYFSFRKS--VLP------FFKVNFSWIFSFFKNKWYIDELYEKLILRPTIYLG-
KGFWKSIDKDLIDNLGPDGISRVIQSFSLVVSRLQSGYLYHYVLSVVIGLTLLISLYNY-
---IF----------
>MAI29530_Rickettsiales_MED689
f__TMED127;g__GCA-002690875;s__GCA-002690875
--MKGLKLFLISVFLPLLSFIVCIFL----QDNISKKHLNFFSCIMMVSATILSFCSLIF
VKI---NGESSIIL-SRWIDSGNLSLDWSLQLNLLTASMVLMVNLVSTLIHIYSVGYMEK
DPKRILFMAYLSLFTFFMLFLVSSSNLLQLFLGWEGVGLTSYLLIGFWNYKEKANSAALK
AFIVNRVGDAGLLLALFTTFVVFGTLNI---SEIKILINSQN---ES--YFNFLGFDIHS
LTLISILLFVGCMGKSAQFGLHVWLPDAMEGPTPVSALIHAATMVTAGVFLLIVMSPLL-
QSSEFSLNFILIVGSVTCIFASSVAVFQNDIKRIIAYSTCSQLGYMFMAIGSSAYTLAYF
HLLSHAFFKALLFLGAGSVIHSMSDEQDIKKMGGLYNKIPLTYITMLIGSLSLI-----G
LPF-----LS---GYYSKDLILEVLYLNDYKYSFHFFLIGLIGVLFTSIYTLRLLVYVFH
RESL------------ADEKVKAH--------------I-HESPMIMVLPLIILSVFAIF
FGMLTKLIFTE---VSFLEMW-SSTMYVNK-GLDASF-ISQN-VPVLFKKLPLMMIFIGI
IVILFLYFSFKN---LSP------FLKLRLSFIYNFFKNKWYVDQLYEKIFVKPTFYFG-
KGFWKSVDKELIDNLGPNGFSRIILSFSFLISRLQSGFLYHYVLSIIIGLTLLISLYTY-
---IF----------
>fig|91750.150.peg.202 AG-899-D02
f__TMED127;g__GCA-002690875;s__GCA-002690875
------------------------------------------MMIIATIFSFFSLVF
VKI---NGESAIIL-SRWIDSGNLSLDWSLQLNILTASMVLMVNLVSTLIHIYSVGYMEK
DPKKILFMAYLSLFTFFMLFLVSSSNLLQLFLGWEGVGLTSYLLIGFWNYKEKANNAALK
AFIVNRVGDAGLLLALFTTFVVFGTLNI---GEIKLLINSQN---ES--YFNFLGFDIHS
LTLISLLLFVGCMGKSAQFGLHVWLPDAMEGPTPVSALIHAATMVTAGVFLLIVMSPLL-
QSSEFSLNFILIIGSITCIFASSVAVFQNDIKRIIAYSTCSQLGYMFMAIGSSAYTLAYF
HLLSHAFFKALLFLGAGSVIHSMSDEQDIKKMGGLYNKIPLTYITMLIGSLSLI-----G
LPF-----LS---GYYSKDLILEILYLNDFEYSLQFFTMGVIGVLFTSIYTLRLLVYVFH
RESV------------ADEKVIAH--------------I-HESPLIMILPLVILSVFAIF
FGMFMKLIFTE---INFLEMW-SYTIHVDK-SLDASF-ISQN-VPVLIKKLPLLMILIGT
IVILFLYFSFRN---LVP------FLKLRLSFIYNFFKNKWYVDQLYERIFVRPTFYFG-
KGFWKSVDKELIDNLGPNGFSRIVLSFSSLISKLQSGFLYHYVLSIIIGLTLLISLYTY-
---IF----------
>fig|91750.541.peg.1061_AG-893-J11_f__TMED127;g__GCA-002690875;s__GCA-
```

```
002690875
---------------------------------------------MMIIATIFSFFSLVF
VKI---NGESAIIL-SRWIDSGNLSLDWSLQLNILTASMVLMVNLVSTLIHIYSVGYMEK
DPKKILFMAYLSLFTFFMLFLVSSSNLLQLFLGWEGVGLTSYLLIGFWNYKEKANNAALK
AFIVNRVGDAGLLLALFTTFVVFGTLNI---GEIKLLINSQN---ES--YFNFLGFDIHS
LTLISLLLFIGCMGKSAQFGLHVWLPDAMEGPTPVSALIHAATMVTAGVFLLIVMSPLL-
QSSEFSLNFILIIGSITCIFASSVAVFQNDIKRIIAYSTCSQLGYMFMAIGSSAYTLAYF
HLLSHAFFKALLFLGAGSVIHSMSDEQDIKKMGGLYNKIPLTYITMLIGSLSLI-----G
LPF-----LS---GYYSKDLILEILYLNDFKYSLQFFTMGVIGVLFTSIYTLRLLVYVFH
RESL------------ADEKVIAH--------------I-HESPLIMILPLVILSVFAIF
FGMFTKLIFTE---INFLEMW-SYTMYVDK-SLNASF-ISQN-VPILIKKLPLLMVLIGT
IVILFLYFSFRN---LIP------FLKFRLSFIYNFFKNKWYVDQLYERIFVRPTFYFG-
KGFWKSVDKELIDNLGPNGFSRIVLSFSSLISKLQSGFLYHYVLSIIIGLTLLISLYTY-
---IF----------
>fig|91750.395.peg.552_AG-915-P07_f__TMED127;g__GCA-002690875;s__GCA-
002690875
----------------------------------------------MIIATIFSVCSLIF
VHF---NGESSIII-SRWINSGNLSLDWSLQLNLLTASMVLMVNLVSTLIHIYSVGYMEK
DPKKILFMAYLSLFTFFMLFLVSSSNLLQLFLGWEGVGLTSYLLIGFWNYKEKANSAALK
AFIVNRVGDAGLLLALFTTFVVFGTLNI---SEIKLLINSQN---ES--IFNFLGFDIHS
LTLISLLLFIGCMGKSAQFGLHVWLPDAMEGPTPVSALIHAATMVTAGVFLLIVMSPLL-
ESSEFSLNFILIVGSITCIFASSVAVFQNDIKRIIAYSTCSQLGYMFMAIGSSAYTLAYF
HLLSHAFFKALLFLGAGSVIHSMSDEQDIKKMGGLYNKIPLTYITMLIGSLSLV-----G
LPF-----LS---GYYSKDLILEIIYLNDFEYSFYFFVMGVIGVLFTSIYSLRLLVYVFH
RDNV------------ADEKVIAH--------------I-HESPSIMILPLVILSIFAIF
FGMFMKIIFTE---FNFLEMW-SSTMYVNK-SLDATF-ISQN-VPVIFKKLPLLMIVIGT
ILILILYFSFKN---LIP------FLKSRLSFLYNFFKNKWYVDQLYEKIFIKPTFYFG-
KGFWKSVDKELIDNLGPNGFSRIVLSFSFFVSRLQSGFLYHYVLSIIIGLTLLISLYTY-
---IF----------
>fig|91750.517.peg.790_AG-892-K07_fTMED127;g__GCA-002690875;s__GCA-
002690875
----------------------------------------------MIIATIFSVCSLIF
VQL---NGESSIII-SRWINSGNLSLDWSLQLNLLTASMVLMVNLVSTLIHIYSVGYMEK
DPKKILFMAYLSLFTFFMLFLVSSSNLLQLFLGWEGVGLTSYLLIGFWNYKEKANSAALK
AFIVNRVGDAGLLLALFTTFVVFGTLNI---SEIKLLINSQN---ES--IFNFLGFDIHS
LTLISLLLFIGCMGKSAQFGLHVWLPDAMEGPTPVSALIHAATMVTAGVFLLIVMSPLL-
ESSEFSLNFILIVGSITCIFASSVAVFQNDIKRIIAYSTCSQLGYMFMAIGSSAYTLAYF
HLLSHAFFKALLFLGAGSVIHSMSDEQDIKKMGGLYNKIPLTYITMLIGSLSLV-----G
LPF-----LS---GYYSKDLILEIIYLNDYEYSFYFFVMGVIGVLFTSIYSLRLLVYVFH
RDNV------------ADEKVIAH--------------I-HESPSIMILPLVILSIFAIF
FGMFMKIIFTE---FNFLEMW-SSTMYVNK-SLDATF-ISQN-VPVIFKKLPLLMIVIGT
ILILILYFSFKN---LIP------FLKSRLSFFYNFFKNKWYVDQLYEKIFIKPTFYFG-
KGFWKSVDNELIDNLGPNGFSRIVLSFSFFVSRLQSGFLYHYVLSIIIGLTLLISLYTY-
---IF----------
>fig|91750.143.peg.416
AG-349-G10_f__TMED127;g__GCA-002690875;s__GCA-002690875
---------------------------------------------MMVIATILSACSLIF
VQS---NGESSIII-SRWIDSGNLSLDWSLQLNLLTASMVLMVNLVSSLIHIYSVGYMEK
```

```
DPKKILFMAYLSLFTFFMLFLVSSSNLLQLFLGWEGVGLTSYLLIGFWNYKEKANSAALK
AFIVNRVGDAGLLLALFTIFVVFGTLNI---SEIKLLINSQN---ES--IFNFLEFDIHS
LTLISLLLFIGCMGKSAQFGLHVWLPDAMEGPTPVSALIHAATMVTAGVFLLIVMSPLL-
EFSEFSLNFILIIGSITCIFASSVAVFQNDIKRIIAYSTCSQLGYMFMAIGSSAYTLAYF
HLLSHAFFKALLFLGAGSVIHSMSDEQDIKKMGGLYNKIPLTYITMLIGSLSLV-----G
LPF-----LS---GYYSKDLILEIIYLNDYEYSFYFFLMGVIGVLFTSIYSLRLLVYVFH
RDSV------------ADEKVIAH--------------I-HESSSIMILPLVILSIFAIF
FGMFMKIVFTE---FNFLEMW-STTMYVNK-SLDATF-ISQN-VPVIFKKLPLLMILTGT
ILILILYFSFKN---LIP------FLKLRLSFFYNFFKNKWYVDQLYERVFIRPTFYFG-
KGFWKSVDKELIDNLGPNGFSRIVLSFSFLVSRLQSGFLYHYVLSIIIGLTLLISLYTY-
---IF----------
>MAI60522_Rickettsiales_MED715 f__TMED127;g__GCA-002690875;s__GCA-002690875
--MKGLKLFLVSVFLPLFSFLVCVFL----QNKLSKKYLNLLSCILMIIATILSFCSLIF
VKI---NGESTILI-SRWIDSGNLSLDWSLQLNILTSSMVLMVNLVSTLIHIYSVGYMEK
DPKKILFMAYLSLFTFFMLFLVSSSNLLQLFLGWEGVGLTSYLLIGFWNYREKANIAALK
AFIVNRVGDVGLLLALFTTFIVFGTLNI---NEIKILINSQY---ES--YFNFLGFNIHS
LSLISILLFIGCMGKSAQFGLHVWLPDAMEGPTPVSALIHAATMVTAGVFLLIVMSPLI-
ESSQFSLNLILVVGSITCIFASSVAVFQNDIKRIIAYSTCSQLGYMFMAIGSSAYTLAYF
HLLSHAFFKALLFLGAGSVIHSMSDEQDIKKMGGLYNKIPLTYITMLIGSLSLI-----G
LPF-----LS---GYYSKDLILEIIYLNDFKYSFHFFLMGVIGVLFTSIYTLRLLVYVFH
RENL------------ADEKVIAH--------------I-HESPLIMILPLVILSVFAIF
FGMLMNSIFIE---INFLEMW-SSTMYVDK-SLDASF-IKQN-VPVFFKKLPLLMIIIGA
IIILFLYFSFRN---IIP------FLKLKLSFIYNFFKNKWYVDQLYERIFVRPTFYFG-
KGFWKSVDNELIDNLGPNGFSKIIYSFSFLISKLQSGFLYHYVLSIIIGLTLLISLYTY-
---IF----------
>MAI77039_Rickettsiales_MED652 f__TMED127;g__GCA-2691245;s__GCA-2691245
-------MFLLSIFLPLISYFFCIFL----TNILKKNLINIISCSLMIISTLLSIISLIL
FKS---NGEHTVML-TNWISSGSLNIDWSVNYNLLTVSMVLMVNFVSTIIHVYSVGYMEK
DPRNTIFMGYLGLFTFFMLFLVSSSNLIQLFVGWEGVGLTSYLLIGFWSHKDEANKASIK
AFIANRVGDFGLLISLFTIFIVFGTINI---NEILLLVNSHS---QS--YFNFFGIQVHS
ITLIVMTMFVGCMGKSAQFGLHIWLPDAMEGPTPVSALIHAATMVTAGVFLLIVMSPLI-
EASNFSKNFILIVGSITCIFASCVAVFQNDIKRIIAYSTCSQLGYMFMAIGVSAYSAAYF
HLLSHAFFKALLFLSAGSVIHSMSDEQNIKKMGGIYNKIPLTYVSMIIGSLALM-----G
IPF-----LS---GYFSKDLILEFIYLSDLNMKLFAFTIGVVGVFLTTVYSSRLIIHVFH
RKNL------------SDEKVYAH--------------I-HESPPVMVLPLVILGVFSIF
FGMYNHDFFAG---QFLENLW-SDVIYVNS-ELNKTY-TLGV-IPKFIKKLPLVMIILGI
IISFLVYFKFPN---QTN------YLKQKLQIPILFFRNKCYIDELYELMFIKPSLYLG-
RGFWKSIDIDLIDNLGPNGMSRMIGRFGLLVSRLQSGYLYHYVLSVVIGLTLFISIYIY-
---IF----------
>fig|1986644.3.peg.1329_TMED65_f__TMED127;g__GCA-2691245;s__GCA-2691245
----------------------------------------------MIISTLLSIISLIL
FKS---NGEHTVML-TNWISSGSLNIDWSVNYNLLTVSMVLMVNFVSTIIHVYSVGYMEK
DPRNTIFMGYLGLFTFFMLFLVSSSNLIQLFVGWEGVGLTSYLLIGFWSHKDEANKASIK
AFIANRVGDFGLLISLFTIFIVFGTINI---NEILLLVNSHS---QS--YFNFFGIQVHS
ITLIVMTMFVGCMGKSAQFGLHIWLPDAMEGPTPVSALIHAATMVTAGVFLLIVMSPLI-
EASNFSKNFILIVGSITCIFASCVAVFQNDIKRIIAYSTCSQLGYMFMAIGVSAYSAAYF
HLLSHAFFKALLFLSAGSVIHSMSDEQNIKKMGGIYNKIPLTYVSMIIGSLALM-----G
```

```
IPF-----LS---GYFSKDLILEFIYLSDLNMKLFAFTIGVVGVFLTTVYSSRLIIHVFH
RKNL------------SDEKVYAH--------------I-HESPPVMVLPLVILGVFSIF
FGMYNHDFFAG---QFLENLW-SDVIYVNS-ELNKTY-TLGV-IPKFIKKLPLVMIILGI
IISFLVYFKFPN---QTN------YLKQKLQIPILFFRNKCYIDELYELMFIKPSLYLG-
RGFWKSIDIDLIDNLGPNGMSRMIGRFGLLVSRLQSGYLYHYVLSVVIGLTLFISIYIY-
---IF----------
>fig|91750.130.peg.1075 AG-898-B20
o__TMED127;f__TMED127;g__GCA-2691245;s__GCA-2691245
---------------------------------------------MIFSTIFAIISLFL
FKE---NSNQQVIL-TNWISSGSLNIDWSVNLNLLTVSMVVMVNFISTIIHIYSVGYMEK
DPRITIFMGYLGLFTFFMLFLVTSSNLIQLFVGWEGVGLTSYLLIGFWSHKEEANKASLK
AFIANRVGDFGLLISLFTIFIVFGTINI---NEIILLVNSHS---QS--YFNFFGFQVHS
ITLIVIMMFIGCMGKSAQFGLHVWLPDAMEGPTPVSALIHAATMVTAGVFLLILMSPLI-
EVSDLGRSIILIVGTITCLFASCVAVFQNDIKRIIAYSTCSQLGYMFMAIGVSAYSVAYF
HLLSHAFFKALLFLGAGSVIHSMSDEQDIKKMGGIYNKIPLTYTTMVIGSLALM-----G
LPF-----LS---GYFSKDMILEFIFLSDQPFKSFAFLVGVIGVFLTTIYSLRLLIHVFH
GKNR------------SDEKVFAH--------------I-HESPTVMVIPLVFLGIFSIF
FGMISHNFFAG---QFLENIW-SDFMYINS-ELNKSY-SLDV-IPKLVKKLPLVMIIVGT
IISFLVYFKFYR---YME------VVKQKMKIPIKFLRNKCYIDELYELIVIKPSLYLG-
RGFWKSIDIDLIDNLGPNGMSRMVSMFGSMVSRLQSGYLYHYVLSVVIGLTLFISIYIY-
---IF----------
>fig|91750.276.peg.716 AG-908-P05 f__TMED127;g__GCA-2691245;s__GCA-2691245
-------MFVLSIFLPFLSYIVCIFL----NNLVREKILNFVSCALMVCSTIASIISLLF
FHL---DGDKSFII-TNWISSGSLNIDWSLNLNLLTVTMVIMVNFVSTIIHVYSVGYMSK
DPRRIIFMGYLGLFTFFMLFLVSSSNLIQLFVGWEGVGLTSYLLIGFWSHKDEANKASLK
AFIANRVGDFGLLISLFTIFIVFGTINI---NEILLLVNSHS---QS--FFNFFGFQIHA
ITLIVVTMFIGCMGKSAQFGLHVWLPDAMEGPTPVSALIHAATMVTAGVFLLILMSPLI-
ETSIFSQNLILIIGTITCLFASCVAIFQNDIKRIIAYSTCSQLGYMFMAIGVSAYSVAYF
HLLSHAFFKALLFLGAGSVIHSMSDEQDIKKMGGIYNRIPLTYTSMVIGSFALM-----G
LPF-----FS---GYFSKDLILEFIYLSDNDFRVFAFVIGILGVFLTSLYSSRLLITVFH
GENC------------SDEKVYAH--------------I-HESPMIMIVPLLTLAFFSIF
FGMIFHNFFAG---SSIELIW-SKFMFVDS-EINQVY-SFGN-FPKLIKKLPLIMILLSV
MIVFLVYFKFPK---IKQ------MVKEKLNFFVNFFYNKCYIDELYEIIIIKPSYYLG-
SGFWKSIDLDLIDNLGPNGISRFVSSLGSKVSKLQSGYLYHYVLSVVIGLTLFISIYIY-
---IF----------
>fig|91750.481.peg.86 AG-891-J16 f__TMED127;g__GCA-2691245;s__GCA-2691245
-------MFVLSIFLPLLSYIVCIFL----NNIVKEKILNFVSCGLMVFSTIASIISLIF
FHL---EGDKSFVI-TNWISSGSLNVDWSLNLNLLTVTMIIMVNFVSTIIHVYSVGYMSK
DPRRNIFMGYLGLFTFFMLFLVSSSNLIQLFVGWEGVGLTSYLLIGFWSHKDEANKASLK
AFVANRVGDFGLLISLFTIFIVFGTINI---NEILLLVNSHS---QS--YFNFFGFQIHS
ITLIVVTMFIGCMGKSAQFGLHVWLPDAMEGPTPVSALIHAATMVTAGVFLLVLMSPLI-
ETSKFSQNLILFVGTITCLFASCVAVFQNDIKRIIAYSTCSQLGYMFMAVGVSAYSVAYF
HLLSHAFFKALLFLGAGSVIHSMSDEQDIKKMGGIYNRIPLTYTSMLIGSLALM-----G
LPF-----FS---GYFSKDLILEFIYLSDSDLKVFAFVTGIFGVFLTALYSSRLLIKVFH
GENN------------SDEKVFAH--------------I-HESPMIMVIPLIILAFFSIF
FGMIFHNFFAG---TLLESIW-SKFMFIDS-ETNQAY-SYGN-VPKLMKKLPLIMILFSI
IIVFLFYFKFPR---FKQ------NVKEKFNFIVKFFYNKCYIDELYEIVIIKPSRYLG-
RGFWKSIDIDLIDNLGPNGISRFVSSLGSKVSNLQSGYLYHYVLSVVIGLTLFISIYIY-
```

```
---IF----------
>fig|91750.508.peg.704_AG-892-F10_f__TMED127;g__GCA-2691245;s__GCA-2691245
-------MFVLSIFLPLLSYVVCIFL----NNVVREKILNFVSCALMVFSTISSIISLIF
FYL---EGDKSFII-TNWISSGSLNVDWSLNLNLLTVTMIIMVNFVSTIIHVYSVGYMSK
DPRRIIFMGYLGLFTFFMLFLVSSSNLIQLFVGWEGVGLTSYLLIGFWSHKDEANKASLK
AFVANRVGDFGLLISLFTIFIVFGTINI---NEILLLVNSHS---QS--YFSFFGFQIHS
ITLIVVTMFVGCMGKSAQFGLHVWLPDAMEGPTPVSALIHAATMVTAGVFLLILMSPLI-
ETSKFSQNLILFIGTITCLFASCVAVFQNDIKRIIAYSTCSQLGYMFMAIGVSAYSVAYF
HLLSHAFFKALLFLGAGSVIHSMSDEQDIKKMGGIYNRIPLTYTSMLIGSLALM-----G
LPF-----FS---GYFSKDLILEFIYLSDSDLKVFAFITGIFGVFLTALYSSRLLITVFH
RENN------------SDEKVFAH--------------I-HESPIIMIIPLVILAFFSIF
FGMIFHNFFAG---SSLETIW-SKFMFIDS-ETNQAY-SYGN-VPKFMKKLPLVMILLSL
ITVFLFYFKFPR---TKQ------LVKDKLNFIVKFFYNKCYIDELYEKVIIKPSKYLG-
RGFWKSIDIDLIDNLGPNGISRFISSLGSKVSNLQSGYLYHYVLSVIIGLTLFISIYIY-
---IF----------
>fig|91750.144.peg.278 AG-349-G20 f__TMED127;g__GCA-2691245;s__GCA-2691245
-------MFLLSIFLPLLSYFVCIFL----NKLVKDKTLNFVSCGLMVLSTICSILAFIF
FYL---NGEKSLVI-TNWISSGSLNVDWSLNLNLLTATMIIMVNFVSTMIHVYSVGYMSK
DPRRIIFMGYLGLFTFFMLFLVSSSNLIQLFVGWEGVGLTSYLLIGFWSHKDEANKASLK
AFVANRVGDFGLLISLFTIFIVFGTINI---NEILLLVNSHS---QS--YFNFFGFQVHS
ITLIVVTMFVGCMGKSAQFGLHVWLPDAMEGPTPVSALIHAATMVTAGVFLLILMSPLI-
ETSKFSQNLILLVGTITCLFASCVAIFQNDIKRIIAYSTCSQLGYMFMAIGVSAYSVAYF
HLLSHAFFKALLFLGAGSVIHSMSDEQDIKKMGGIYNRIPLTYTSVLIGSLALM-----G
LPF-----LS---GYFSKDLILEFIYLSDSDFKIFAFGIGIFGVFLTSLYSSRLLIRVFH
GENN------------SDEKVYAH--------------I-HESPMIMLVPVVILAFFSIF
FGMIFHNFFAG---PSIETIW-SKFMFVDS-EIHQAY-SIGS-VPKLIKKLPLIMIFFSV
TVVLLFYFKFSR---TMQ------MLKEKLRFFVKFFHNKCYIDELYEIIIIKPSYYLG-
RGFWKSIDNDLIDNLGPNGISRFVSSLGIKVSKLQSGYLYHYVLSVVIGLTLFISIYIY-
---IF----------
>fig|91750.172.peg.1012 AG-359-F09 f__TMED127;g__GCA-2691245;s__GCA-2691245
-------MFLLSIFLPLLSYFVCIFL----NNLVKDKTLNFVSCGLMVFSSICSIFAFIF
FHF---DGEKSLVI-TNWISSGSLNVDWSLNLNLLTATMIIMVNFVSTLIHVYSVGYMSK
DPRRIIFMGYLGLFTFFMLFLVSSSNLIQLFVGWEGVGLTSYLLIGFWSHKDEANKASLK
AFIANRVGDFGLLISLFTIFIVFGTINI---NEILLLVNSHS---QS--YFNFFGFQVHS
ITLIVVTMFVGCMGKSAQFGLHVWLPDAMEGPTPVSALIHAATMVTAGVFLLILMSPLI-
ETSKFSQNLILLVGTLTCLFASCVAIFQNDIKRIIAYSTCSQLGYMFMAIGVSAYSVAYF
HLLSHAFFKALLFLGAGSVIHSMSDEQDIKKMGGIYNRIPLTYTAVIIGSLALM-----G
LPF-----FS---GYFSKDLILEFIYLSESDFKLFAFGIGIFGVFLTSLYSSRLLVRVFH
GENN------------SDEKVYAH--------------I-HESPMVMLIPLVILAFFSIF
FGMIFHNFFVG---PSIESIW-SKFMFVDS-EMHQAY-SIGN-VPKLMKKLPLIMICLSV
IVVFLFYFKFSR---TKE------MLKEKLKFFVKFFYYKCYIDELYEIIIIKPSYYLG-
RGFWKSIDIDLIDNLGPDGISRFVSSLGIKVSKLQSGYLYHYVLSVVIGLTLFISIYIY-
---IF----------
>fig|91750.247.peg.694 AG-907-I19 f__TMED127;g__GCA-2691245;s__GCA-2691245
-------MFLLSIFLPLLSYFICVFF----FGLIKDKVLSFVSCFLMISSTLFSIISLIL
LQI---NGDQSYVL-TNWISSGSLNIDWSLNLNLLTASMVIMVNFISSIIHIYSVGYMSK
DSRKSIFMGYLGLFTFFMLFLVSSSNLIQLFVGWEGVGLTSYLLIGFWSYKEEANKASLK
```

```
AFIANRVGDFGLLISLFTIFIVFGTINI---NEILLLVNSHS---QS--YFNFFGFQVHS
ITLIVTTMFIGCMGKSAQFGLHVWLPDAMEGPTPVSALIHAATMVTAGVFLLILMSPLI-
EVSDFGKKLILIVGSITCLFASCVAIFQNDIKRIIAYSTCSQLGYMFMAVGVSAYSVAYF
HLLSHAFFKALLFLAAGSVIHSMSDEQNIKKMGGIYNRIPLTYTSMLIGSFALM-----G
LPF-----LS---GYYSKDLILEFIFLSEMSFKFIAFFIGIFGVLLTTIYSSRLLIYVFH
GQNR------------SDEKVFAH--------------I-HESPLIMVLPLVILAFFSIF
FGMYFYSSFVG---PSLEEIW-SKFMFVNS-ETNQIY-SLGN-IPNIIKKLPLFMIITGL
ILSFLIYFKFSK---SLP------VIKQRLHLIITFFYRKCFIDELYQMILIKPCIYLG-
RGFWKSIDMDLIDNLGPNGISRLVNSFGLMVSKLQSGYLYHYVLSVVIGLTLFISIYIY-
---IF----------
>fig|1913988.3564.peg.1438 LRMED-288_f__TMED127;g__GCA-2691245
----------------------------------------------MVASTFFSILSFIY
LKL---NGDQNIVL-SNWISSGSLNIDWSINLNLLTSSMVLMVNFVSTIIHIYSVEYMSK
DSRKIIFMGYLSLFTFFMLFLVSSSNLIQLFLGWEGVGLTSYLLIGFWSHKNEANKASIK
AFVANRVGDFGLLISLFTIFIVFGTINI---NEILLLVNSHS---QS--YFNFLGFQVHS
ITLIVITMFIGCMGKSAQFGLHVWLPDAMEGPTPVSALIHAATMVTAGVYLLILMSPLI-
ESSLLAQNIILIVGSITCLFASCVAIFQNDIKRIIAYSTCSQLGYMFMAIGVSAYSTAYF
HLLSHAFFKALLFLSAGSVIHSMSDEQDIKKMGGIYNRIPLTYVCMLIGSFALM-----G
LPF-----LS---GYYSKDLILEFIFLSEHPFKLFAFFTGIVGVLLTTIYSSRLIIHVFH
RKNL------------SDEKVFAH--------------I-HESPPIMIYPLIVLAIFSIF
FGMLFHHFFAG---QNLEKIW-TNIMFINL-EVNNSY-YLNE-IPKIIKKLPLIMVLIGI
LISYLIYFKLEV---YNN------ILKKRFKIIINFFYNKCYIDELYDFLIIKPSFYLG-
KGFWRSIDNDLIDNLGPNGISRLIGTLGSYVSRLQSGYLYHYVLSVIIGLTLFISIYIY-
---IF----------
>fig|91750.324.peg.165_AG-911-L17_f__TMED127;g__GCA-2691245;s__GCA-2691245
----------------------------------------------MILSSILSILSLLY
VIL---SGDQNYIL-SNWISSGSLNIDWSLNLNLLTATMVLMVNFVSTIIHIYSIGYMSK
DPRRIIFMGYLSLFTFFMLFLVSSANLIQLFVGWEGVGLTSYLLIGFWSKKEIANKASLK
AFIANRVGDFGLLISLFTIFIVFGTINI---DEILLLVNSHS---QS--YFNFLGFEIHS
ITLIVIFMFIGCMGKSAQFGLHVWLPDAMEGPTPVSALIHAATMVTAGVFLLILMSPLI-
ETSPFAQNFILIIGSITCLFASCVAIFQNDIKRIIAYSTCSQLGYMFMAIGVSAYSMAYF
HLLSHAFFKALLFLGAGSVIHCMSDEQDIKKMGGIYNKIPLTYICMLIGSIALM-----G
LPF-----LS---GYYSKDLILEFIYLSDLNIKFFSFMIGIVGVLLTTIYSSRLIIHVFH
KNNR------------SDEKVFAH--------------I-HESPLIMVFPLVILAFFSIF
FGMIFHNFFAG---SFLEDIW-SGSIFIDL-EANNFY-TLGY-IPTAVKKLPLIMIISGL
FISWMYYYKLAN---KLE------YIKQKSSFIINFFYNKCYIDELYEFLFIRPSIYLG-
KGFWKSIDIDLIDNLGPNGMSRLIGSFGAVVSRLQSGYLYHYVLSVVIGLTLFISIYVY-
---IF----------
>MAJ67247_Rickettsiales_MED617_f__TMED127;g__TMED13;s__TMED13
-------MFLLSIFLPLFSFLVCVTL----NKKLKDKYVCLTSSVFLVFSSICAFISFLN
LET---NQDQAIVL-SSWINSGSFSVDWSIQYNLLSSSMVLMVNFVSTLIHIYSVGYMAK
DDRKVVFMGYLGLFTFFMLFLVSSSNLLQLFLGWEGVGLTSYLLIGFWYKKNQANVAAIK
AFVVNRVGDFGLILALFATFIVFGTLNI---NEVLLLLESQN---TT--KVEFLGFQLHS
ITLIVVLIFIGCMGKSAQFGLHVWLPDAMEGPTPVSALIHAATMVTAGVFLLIILSPLIE
ESSQLSRSLILIIGTITCIFASSVAIFQNDIKRIIAYSTCSQLGYMFMAIGVSAYSVAYF
HLLSHAFFKALLFLGAGSVIHSMSDEQDIKKMGGLYNKIPLTYVMMLIGSFSLM-----G
LPF-----LS---GYYSKDLIIEFIYLGQSDVKTYAYIISIISVLFTTIYSVRLIIYVFH
RENK------------SDEKVFAH--------------I-HESPLIMILPLTILSIFAIF
```

```
FGMLMHNYFAG---ENLFKLW-GEFMFINN-DINDEY-TIGN-VPVLVKKTPLIAIILGI
LFCFLIYFVFKN---YSQ------AMKKKLRLIVNFFENKLYVDEIYNFIFVKSSFYLG-
KGFWKSIDTDLIDNLGPNGISRLIGSFGKVVSRLQSGYLYHYVLSVVVGLTLFLSIYIY-
---IL----------
>fig|1986627.3.peg.661_TMED13_f__TMED127;g__TMED13;s__TMED13
---------------------MCVTL----NKKLKDKYVCLTSSVFLVFSSICAFISFLN
LET---NQDQAIVL-SSWINSGSFSVDWSIQYNLLSSSMVLMVNFVSTLIHIYSVGYMAK
DDRKVVFMGYLGLFTFFMLFLVSSSNLLQLFLGWEGVGLTSYLLIGFWYKKNQANVAAIK
AFVVNRVGDFGLILALFATFIVFGTLNI---NEVLLLLESQN---TT--KVEFLGFQLHS
ITLIVVLIFIGCMGKSAQFGLHVWLPDAMEGPTPVSALIHAATMVTAGVFLLIILSPLIE
ESSQLSRSLILIIGTITCIFASSVAIFQNDIKRIIAYSTCSQLGYMFMAIGVSAYSVAYF
HLLSHAFFKALLFLGAGSVIHSMSDEQDIKKMGGLYNKIPLTYVMMLIGSFSLM-----G
LPF-----LS---GYYSKDLIIEFIYLGQSDVKTYAYIISIISVLFTTIYSVRLIIYVFH
RENK------------SDEKVFAH--------------I-HESPLIMILPLTILSIFAIF
FGMLMHNYFAG---ENLFKLW-GEFMFINN-DINDEY-TIGN-VPVLVKKTPLIAIILGI
LFCFLIYFVFKN---YSQ------AMKKKLRLIVNFFENKLYVDEIYNFIFVKSSFYLG-
KGFWKSIDTDLIDNLGPNGISRLIGSFGKVVSRLQSGYLYHYVLSVVVGLTLFLSIYIY-
---IL----------
>fig|91750.392.peg.642 AG-915-O07 f__TMED127;g__TMED13;s__TMED13
-------MFLLSIFLPLSSFLFCITL----NKRLKDEYICISSSFILIFSSICAFLSFLN
LET---NQDQAIVL-SNWINSGSFSVDWSVQYNLLSSSMVLMVNFVSTLIHIYSVGYMVK
DDRKVIFMGYLGLFTFFMLFLVSSSNLLQLFLGWEGVGLTSYLLIGFWYKKNQANVAAIK
AFVVNRVGDFGLILALFATFIVFGTLNI---NEILLLLESQN---TT--RVEFLGFQFHS
ITLIVVLIFIGCMGKSAQFGLHVWLPDAMEGPTPVSALIHAATMVTAGVFLLIILSPLIE
ESSQISRSIILIIGTITCIFASSVAIFQNDIKRIIAYSTCSQLGYMFMAIGVSAYSVAYF
HLLSHAFFKALLFLGAGSIIHSMSDEQNIKKMGGLYNKIPLTYITMLIGSFSLM-----G
LPF-----LS---GYYSKDLIIEFIYLGQSNIKIYAYIISIVSVLFTTIYSVRLLIYVFH
RENR------------SDEKVFAH--------------I-HESPLIMILPLTILSIFAIF
FGMLMNSYFAG---ESLFKVW-GEFMFINN-DINDEY-TIGN-VPILVKKSPLFAIIFGI
FICFLIYFTFKE---YSQ------ALKKRLMILVNFFENKLYIDEIYNFIFVKTSFYLG-
KGFWKSIDTDLIDNLGPNGISRLVGSFGKVVSRFQSGYLYHYVLSVVVGLTLFLSIYIY-
---IL----------
>fig|1913988.3131.peg.248_139575_S176_rosella_bins.tsv.030 f__TMED127;g__TMED13;s__TMED13
-------MFLLSIFLPLLSFLFCITV----NRRFKDEYICISSSIILIFSSICAFLSFLN
LEI---NQDQAIVL-SSWINSGSLSVDWSVQYNLLSSSMVLMVNFVSTLIHIYSIGYMVK
DDRKVIFMGYLGLFTFFMLFLVSSSNLLQLFLGWEGVGLTSYLLIGFWYKKNQANVAAIK
AFVVNRVGDFGLILALFATFIVFGTLNI---NEILLLLDSQN---TT--RVEFLGFQLHS
ITLIVVLIFIGCMGKSAQFGLHVWLPDAMEGPTPVSALIHAATMVTAGVFLLIILSPLIE
ESSQISRSIILIIGTITCIFASSVAIFQNDIKRIIAYSTCSQLGYMFMAIGVSAYSVAYF
HLLSHAFFKALLFLGAGSVIHSMSDEQNIKKMGGLYNKIPLTYITMLIGSFSLM-----G
LPF-----LS---GYYSKDLIIEFIYLGQSNIKIYAYMISIVSVLFTTIYSVRLLIYVFH
RENR------------SDEKVFAH--------------I-HESPLIMILPLTILSIFAIF
FGMLMNSYFAG---DSLFKVW-GEFMFINN-DINDEY-TIGN-VPNLVKKSPLFAIILGI
FICSLIYFVFKD---YSQ------VLKKKLLILVNFFENKLYIDEIYNFIFVKTSFYLG-
KGFWKSIDTDLIDNLGPNGISRLVGSFGKVVSRFQSGYLYHYVLSVVVGLTLFLSIYIY-
---IL----------
>MBF91430_Rickettsiales_SP91_f__TMED127;g__GCA-2715305;s__GCA-2715305
```

```
-------MFLFAIFLPLLSFFLCISL----NRTIKDSYINILSCSLLVLSAVFSLCSIFF
LGW---DKQQTYII-SNWIVSGSFSVDWSINYNLLTCSMVLMVNFVSSLIHIYSVGYMEH
DSKRTIFLGYLSLFTFFMLILVSSSNLVQLFLGWEGVGLASYLLIGFWNHKNVANSAALK
AFIVNRVGDFGILIGLFTTFMVFGTLNI---NEILILVNSHI---DS--YFNFLGLQIHS
LTLIAILFFVGCMGKSAQFGLHIWLPDAMEGPTPVSALIHAATMVTAGVFLLILVSPII-
EASDFSRKVIIIIGSLTCIFAASVAFFQNDIKRIIAYSTCSQLGYMFMAIGASAYNAAYF
HILSHAFFKALLFLSAGSVIHSMSDEQNIKKMGGLFNKIPMTYSCMLIGSVALV-----G
LPF-----FS---GYFSKDLILELIYLGNSEIRLYAFFVGVLAVLFTSLYSTRLIIHVFH
GKSQ------------ADEKVLAH--------------V-HESPKIMIIPLVILSIFSIF
FGMLSYDLFFG---YNSNILW-SEHFFINT-ELNDKS-LIDN-IPMYIKKSPLIMILFGI
SLSFLIYYPFKQ---VLP------FIKEKLIIVHKFFYNKWYIDEIYNFLIIRPIIYLG-
KGFWKSIDIDLIDTLGPNGISNFVRRFGVIVSSFQSGFLYHYALSVIIGLTLFLSLYVH-
---IF----------
>MBC34057_Rickettsiales_SP101 f__TMED127;g__GCA-2711625;s__GCA-2711625
-----MTLFLFSIFLPLXSFILCLFF----KDLFKDKYLNLTSCFLLSLSTIFSLVSILY
YGM---DNSKTFLI-XNWITSGSXVIDWSVNYTLLTGSMVLMVNFISTLIHIYSIGYMSN
DPKKILFMGYLGLFTFFMLILVTSSNLVQLFLGWEGVGLTSYLLIGFWNHKTIANKAALK
AFVVNRIGDFGILIALFTTFIIFGTLDI---NEILILVNSHL---ET--SFSFLGFDIHV
LTLISMLFFIGCMGKSAQFGLHTWLPDAMEGPTPVSALIHAATMVTAGVFLLIILSPLI-
ETSDFAKDFILIIGSLTCIFAASVGFFQDDIKRIIAYSTCSQLGYMFMAIGASAYSAAYF
HLLSHAFFKALLFLGAGSVIHSMSDEQNIKKMGGLYDKIPVTYLCMLIGSVALI-----G
LPF-----FS---GYYSKDLILELIYLGKNNINFYAAILGVIAVFFTSLYSLRLIIYVFH
GQSR------------SDEKVLAH--------------V-HESPFIMVGPLVLLSIFSIF
FGMATHEYFFG---AKSSILW-SGLLYINN-ELNDYS-LIEG-IPFYIKKIPLLMILIAL
IISLILYYPLKG---ILP------VMKEKLILIFKLFSNKFYIDEIYDFFIVRPIFYLG-
KGFWKSIDNDLIDSLGPNGISNFVKKFGLIVSSFQSGFLYHYALSVIVGLTLFLSLYIY-
---IL----------
>MAI84862_Rickettsiales_MED657_o_TMED127
… f__TMED127;g__GCA-2691145;s__GCA-2691145
-------MFSFAVFLPLLSYFFCIIA----NGKIKDQNIEYFASCLILLSTIISIFCFFA
VSD--ENPQINIQL-AEWLSSGSLYIDWSLSFNRLSSLMVLLVNFISCLVHFYSIGYMSN
DQKKIIFIGYLGLFTFFMLILVTSSNLIQLFLGWEGVGLTSYLLIGFWNFKDVANKAAIK
AFIVNRVGDFSFLIGILVIFILFGTLNF---NEIFILVQSQK---NT--YFELFDINFHA
LTLISVLLFIGCMGKSAQLGLHTWLPDAMEGPTPVSALIHAATMVTAGVFLLVLMSPII-
EESQLAQNMIMIVGALTSFFAATVAITQDDIKRIIAYSTCSQLGYMFVAIGASAYGAAMF
HLVTHAFFKALLFLGAGSVMHAMSDEQNIKKMGGLYKKIPVTYLLMLIGTLSIT-----G
FPF-----FS---GYFSKDLIIEILYFDKSIFREFAFFTSVFVVFLTSFYSFRLLIYVFH
GKNN------------SDEKVLAH--------------V-HESPNVMLIPLIILSIFAIF
SGILLNKYFFG---INSILFW-KDSLYNIS-TTNLPT-LIFE-MPFIIKKIPFFMALLGL
FICILFYRFFPN---ITY------FLKKRFSLVYIFLKNKWYFDEFYDFLFVKPLRYIG-
NGFWKSIDIELIDNVGPNGISKLIKKFGMLISIFHTGYLYHYAMSIIIGLTVFISIYFY-
---IF----------
>fig|91750.153.peg.932 AG-899-E20 f__TMED127;g__GCA-2691145;s__GCA-2691145
-------MFSFAVFLPLLSYFFCIIA----NGKIKDQNIEYFASCLILLSTIISIFCFFA
VSD--ENPQINIQL-AEWLSSGSLYIDWSLSFNRLSSLMVLLVNFISCLVHFYSIGYMSN
DQKKIIFIGYLGLFTFFMLILVTSSNLIQLFLGWEGVGLTSYLLIGFWNFKDVANKAAIK
AFIVNRVGDFSFLIGILVIFILFGTLNF---NEIFILVQSQK---NT--YFELFDINFHA
LTLISVLLFIGCMGKSAQLGLHTWLPDAMEGPTPVSALIHAATMVTAGVFLLVLMSPII-
```

```
EESQLAQNMIMIVGALTSFFAATVAITQDDIKRIIAYSTCSQLGYMFVAIGASAYGAAMF
HLVTHAFFKALLFLGAGSVMHAMSDEQNIKKMGGLYKKIPVTYLLMLIGTLSIT-----G
FPF-----FS---GYFSKDLIIEILYFDKSIFREFAFFTSVFVVFLTSFYSFRLLIYVFH
GKNN------------SDEKVLAH--------------V-HESPNVMLIPLIILSIFAIF
SGILLNKYFFG---INSILFW-KDSLYNIS-TTNLPT-LIFE-MPFIIKKIPFFMALLGL
FICILFYRFFPN---ITY------FLKKRFSLVYIFLKNKWYFDEFYDFLFVKPLRYIG-
NGFWKSIDIELIDNVGPNGISKLIKKFGMLISIFHTGYLYHYAMSIIIGLTVFISIYFY-
---IF----------
>fig|91750.491.peg.245_AG-919-E11_f__TMED127;g__GCA-2691145;s__GCA-2691145
-------MFSFAVFLPLLSYFFCIIA----NGKIKDQNIEYFASCLILLSTIISIFCFFA
VSD--ENPQINIQL-AEWLSSGSLYIDWSLSFNRLSSLMVLLVNFISCLVHFYSIGYMSN
DQKKIIFIGYLGLFTFFMLILVTSSNLIQLFLGWEGVGLTSYLLIGFWNFKDVANKAAIK
AFIVNRVGDFSFLIGILVIFILFGTLNF---NEIFILVQSQK---NT--YFELFDINFHA
LTLISVLLFIGCMGKSAQLGLHTWLPDAMEGPTPVSALIHAATMVTAGVFLLVLMSPII-
EESQLAQNMIMIVGALTSFFAATVAITQDDIKRIIAYSTCSQLGYMFVAIGASAYGAAMF
HLVTHAFFKALLFLGAGSVMHAMSDEQNIKKMGGLYKKIPVTYLLMLIGTLSIT-----G
FPF-----FS---GYFSKDLIIEILYFDKSIFREFAFFTSVFVVFLTSFYSFRLLIYVFH
GKNN------------SDEKVLAH--------------V-HESPNVMLIPLIILSIFAIF
SGILLNKYFFG---INSILFW-KDSLYNIS-TTNLPT-LIFE-MPFIIKKIPFFMALLGL
FICILFYRFFPN---ITY------FLKKRFSLVYIFLKNKWYFDEFYDFLFVKPLRYIG-
NGFWKSIDIELIDNVGPNGISKLIKKFGMLISIFHTGYLYHYAMSIIIGLTVFISIYFY-
---IF----------
>MDC3091444_Rickettsiales_AH-716-M15 f__TMED127;g__TMED211;s__TMED211
-------MFFLSVFLPLISFVFCIFF----NNKIKDTSIELIVSIFLSLSTLSSLYCLFF
ILT--GNPAIIVNL-GEWLVSGGLFIDWSLNFNILSSVMVLVINFISSIVHIYSIGYMEH
DNKKIIFLGYLGLFTFFMLILVTSSNLIQLFLGWEGVGLTSYLLIGFWNYKNSANNAALK
AFVVNRVGDFGFLIGIFAIFILFGTVNL---NEIFLLVDSQR---GT--YFSFLSFDFHA
LTLISILLFIGCMGKSAQFGLHTWLPDAMEGPTPVSALIHAATMVTAGVFLLVLMSPII-
EQSEFAKKLIMIVGALTSFFAASVAITQDDIKKIIAYSTCSQLGYMFVAIGASAYGAAIF
HLVTHAFFKALLFLGAGSVIHAMSDEQNIKKMGGLYNRIPVTYALMLVGTLSLV-----G
FPF-----FS---AYYSKDLILEVLYLSNLSIKNYIYFIGVFVVFLTSFYSFRLLIHVFH
GQCK------------ADEKVRAH--------------I-HESPFVMILPLVVLSFFSIF
AGMIFSSYFFG---INSILFW-KDSIINIA-NTEIIS-TIKD-IPIVIKKLPLVMVLLGF
FTSILLYFFFKN---ITA------FLKEKFNFIYIFLKNKWFVDELYEYTIVSSMKYLG-
NGFWKSIDQELIDNIGPNGISKMIKRVGSFVSYLQTGYLYHYALTVIVGLTIFISIYFY-
---NI----------
>MBH43546_Rickettsiales_SP4073_f__TMED127;g__SP4073;s__SP4073
-------MFFLSVFIPLICYFFCIFF----NKKIAEKKIEFFISSLMTLASLLSIYVFFS
VTE--FNPESQIII-GNWLSSGSFSIDWSLNFNRLSVLMVLVVNIVSTVVHIYSIGYMHN
DPKKVIFLGYLGLFTFFMLILVTSSNLIQLFLGWEGVGLTSYLLIGFWNFKDVANKAALK
AFVVNRIGDFGFLLGIFAIFFIFGTVNF---NEIFILVESQE---NT--FFKFLGFNFHA
LTLISVLLFIGCMGKSAQFGLHTWLPDAMEGPTPVSALIHAATMVTAGVFLLVLMSPII-
EQSEFAQKLIMIIGALTSLFAASVAITQDDIKRIIAYSTCSQLGYMFIAIGCSAYGVAMF
HLVTHAFFKALLFLGAGSVIHAMSDEQNIKKMGALYNKIPVTYILMVIGTLSLV-----G
LPF-----FS---AYYSKDLILEILYLDNFDISIHIYIIGIIVVFFTSFYSFRLLIYVFH
GENK------------SDEKVVAH--------------I-HESPRIMIYPLLILSFFSIF
SGLLFKNYFFG---FESLSFW-GTSIYNIF-ESDISY-AVKE-LPLSIKKLPFLMILLGF
VASLAICRHLIK---LNL------YLKQKLNLIYMFLKNKWYVDEFYDNLVVKPTKFIG-
```

```
YGFWKSIDKELIDNIGPNGVSKLIKKFGIFISSLQTGYLYHYALTIIIGLTIFISIYFY-
---S-----------
>fig|91750.74.peg.862 AG-337-B08 f__TMED127;g__SP4073;s__SP4073
-------MFSLSVFIPLICYFFCIFF----NKRIAEIKIEIVISALMILASLISIYVFFS
VSE--FNPESQIII-GNWLSSGSFLIDWSLNFNSLSVLMVLVVNVVSTIVHIYSIGYMHN
DPKKIIFLGYLGLFTFFMLILVTSSNLIQLFLGWEGVGLTSYLLIGFWNFKDVANKAALK
AFVVNRIGDFGFLLGIFAIFFIFGTVNF---NEIFILVESQE---DT--FFKFLGFNFHA
LTLISVLLFIGCMGKSAQFGLHTWLPDAMEGPTPVSALIHAATMVTAGVFLLVLMSPII-
EQSEFAQKLIMIIGALTSLFAASVAITQDDIKRIIAYSTCSQLGYMFVAIGCSAYGVAMF
HLVTHAFFKALLFLGAGSVIHAMSDEQNIKKMGALYNKIPVTYMLMLIGTFSLV-----G
LPF-----FS---AYYSKDLILEILYLDDFDISIYVYIIGIIVVFFTSFYSFRLLIYVFH
GESK------------SDERVAAH--------------I-HESPGIMIYPLLILSFFSIF
SGLLFKNYFFG---FESLSFW-GNTIYNIF-ESDISY-AVNE-LPLSIKKLPFMMIILGF
LASLAVCRYLTK---LNL------YLKHKLGFLYTFLKNKWYVDEFYDFFVIKPTKFIG-
YGFWKSIDKELIDNIGPNGVSKLIKKFGIFVSSLQTGYLYHYALTIIIGLTIFISIYFY-
---N-----------
>fig|91750.72.peg.1260_AG-337-B03_f__TMED127;g__SP4073;s__SP4073
-------MFSLSVFIPLICYFFCIFF----NKRIAEIKIEIVISALMILASLISIYVFFS
VSE--FNPESQIII-GNWLSSGSFLIDWSLNFNSLSVLMVLVVNVVSTIVHIYSIGYMHN
DPKKVIFLGYLGLFTFFMLILVTSSNLIQLFLGWEGVGLTSYLLIGFWNFKDVANKAALK
AFVVNRIGDFGFMLGIFAIFFIFGTVNF---DEIFILVESQE---NT--FFKFLGFNFHA
LTLISVLLFIGCMGKSAQFGLHTWLPDAMEGPTPVSALIHAATMVTAGVFLLVLMSPII-
EQSEFAQKLIMVIGALTSLFAASVAITQDDIKRIIAYSTCSQLGYMFIAIGCSAYGVAMF
HLVTHAFFKALLFLGAGSVIHAMSDEQNIKKMGALYNKIPVTYMLMLIGTFSLV-----G
LPF-----FS---AYYSKDLILEILYLDDFDISIYVYIIGIIVVFFTSFYSFRLLIYVFH
GESK------------SDERVAAH--------------I-HESPRIMIYPLLILSFFSIF
SGLLFKNYFFG---FESLSFW-GNTIYNIF-ESDISY-AVKE-LPLSIKKLPFMMIILGF
LVSLAVCRYLTK---LNL------YLKHKLSFLYVFLKNKWYVDEFYDFFIIKPTKFIG-
YGFWKSIDKELIDNIGPNGVSKLIKKFGIFVSSLQTGYLYHYALTIIIGLTIFISIYFY-
---S-----------
>fig|91750.84.peg.1475_AG-337-E20_f__TMED127;g__SP4073;s__SP4073
-------MFSLSVFIPLICYFFCIFF----NKRIAEIKIEIVISALMILASLISIYVFFS
VSE--FNPESQIII-GNWLSSGSFLIDWSLNFNSLSVLMVLVVNVVSTIVHIYSIGYMHN
DPKKVIFLGYLGLFTFFMLILVTSSNLIQLFLGWEGVGLTSYLLIGFWNFKDVANKAALK
AFVVNRIGDFGFMLGIFAIFFIFGTVNF---NEIFILVESQE---NT--FFKFLGFNFHA
LTLISVLLFVGCMGKSAQFGLHTWLPDAMEGPTPVSALIHAATMVTAGVFLLVLMSPII-
EQSEFAQKLIMIIGALTSLFAASVAITQNDIKRIIAYSTCSQLGYMFVAIGCSAYGVAMF
HLVTHAFFKALLFLGAGSVIHAMSDEQNIKKMGALYNKIPVTYMLMLIGTFSLV-----G
LPF-----FS---AYYSKDLILEILYLDDFDISIYVYIIGIIVVFFTSFYSFRLLIYVFH
GESK------------SDERVAAH--------------I-HESPRIMIYPLLILSFFSIF
SGLLFKNYFFG---FESLSFW-GNTIYNIF-ESDISY-AVKE-LPLSIKKLPFLMIILGL
LASLAVCRYLTK---LNL------YLKHKLCFLYTFLKNKWYVDEFYDFFIIKPTKFIG-
YGFWKSIDKELIDNIGPNGVSKLIKRFGIFVSSLQTGYLYHYALTIIIGLTIFISIYFY-
---S-----------
>MAH78768_Rickettsiales_MED723_f__TMED127;g__GCA-2710765;s__GCA-2710765
-------MYFLSVFLPLFSFIFCIVS----NKRLSDIKVEFIVCSILLISSLMAVLSFFN
ISQ--IESSEIVIL-GDWLTSGGFYINWSLSFNRLSAIMVVVVNLVSLLVHIYSVGYMAN
DNNKIVFLGYLGLFTFFMLILVTSSNLIQLFLGWEGVGLTSYLLIGFWSYKDSANKAALK
```

```
AFVVNRVGDFGFLLGIFTIFIVFGTVNF---DEIFILVDTQK---ST--FFNFLGMEFNS
LTLISILLFIGCMGKSAQFGLHTWLPDAMEGPTPVSALIHAATMVTAGVFLLVLMSPLI-
EKSDFAQNFIMIIGSLTSLFAASVALAQDDIKKIIAYSTCSQLGYMFIAIGSSAYGIAMF
HLVTHAFFKALLFLGAGSVIHSMSDEQNIKKMGGLYSKIPVTYFLMLIGTLSLV-----G
FPF-----FS---AYYSKDLILEVLYLDNSLFKSYAYIIATIVVFLTSFYSFRLLICVFH
GKNN------------SDEKVLAH--------------V-HESPNIMILPLIVLSFFSIF
SGLIFHDYFFG---VDSILFW-KDTIYNIS-QTETPV-LVND-LPFYIKKLPLLMMLLGL
IFAILLYSFLKN---VRL------FLKKNLRFFYLFLKNKWYIDEIYKFLIISPINNLG-
RGFWKSIDEEFIDNIGPNGISKVVKRLGFLISSLQTGYLYHYALTVIIGLTIFISIYFY-
---VI----------
>MAY89615_Rickettsiales_SAT45_f__TMED127;g__GCA-2710765;s__GCA-2710765
-------MYFLSVFLPLFSFIFCIVS----NKRLSDIKVEFIVCSFLLISTLMAILSFFN
ISE--IESSEIVIL-GDWLTSGGFYINWSLSFNRLSAIMVVVVNLVSLLVHIYSIGYMAN
DDNKIVFLGYLGLFTFFMLILVTSSNLIQLFLGWEGVGLTSYLLIGFWSYKDSANKAALK
AFVVNRVGDFGFLLGIFTIFIVFGTVNF---DEIFILVDTQK---ST--FFNFLGMEFNS
LTLISILLFIGCMGKSAQFGLHTWLPDAMEGPTPVSALIHAATMVTAGVFXLVXMSPLX-
EXSDFAQNXIMIIGXLTSXFAASVALAQXDIKKXIAYSTCSQLGYMFIAIGSSAYGXAMF
HLVTHAFFXALLFLGAGSVIHSMSDEQNIKKMGGLYSKIPVTYFLMLIGTLSLV-----G
FPF-----FS---AYYSKDLILEVLYLDNSLFKSYAYIVAIIVVFLTSFYSFRLLICVFH
GENN------------SDEKVLAH--------------V-HESPNTMILPLIVLSFFSIF
SGLIFHDYFFG---VDSILFW-KDTIYNIS-QTETPV-LVND-LPIYIKKLPLLMMLLGL
TFVILLYSYLKN---VRL------FFKKNLRFFYLFLKNKWYVDEIYKFLIIGPINNLG-
RGFWKSIDEEFIDNIGPNGISKVVKRLGFFISSLQTGYLYHYALTVIIGLTIFISIYFY-
---VI----------
>fig|91750.91.peg.386_AG-896-A16_f__TMED127;g__CACBWF01;s__CACBWF01
-------MHILSVFLPLLSFLIAFFL----SRKISDEIVDYTTCTMMAFSTILSLYCFFL
INE--YNPERIIKV-SEWLNSGSLYIDWSLNFNRLTALMVFLVNFVSTIVHVYSVGYMNN
DDRRPIFMAYLGLFTFFMLMLVTSSNLIQLFLGWEGVGLTSYLLIGYYSYKESANKAAIK
AFIVNRVGDFGFLIGIFAIFIVFGTINL---KEIFLLVDTQE---NT--YFNFLGYNFHT
LTLITMLLFIGAMGKSAQFGFHTWLPDAMEGPTPVSALIHAATMVTAGVFLMVLMSPLL-
EKTEFTRNFIIVIGSLSALFAATVAITQDDIKKIIAYSTCSQLGYMFIAIGVSAYNIAIF
HLFTHAFFKALLFLGAGSVIHAMSDEQNIKKMGGLYKKIPVTYILMLVGTLSLT-----G
FPF-----FS---AYYSKDLIMELVFLDDSYLSNFVFINSVIVVFLTSFYSFRLLFYVFH
GNNR------------SDEKVFSH--------------V-HESPNIMLLPLVVLSFFSVF
SGLAFKKYFFG---IDSIVFW-GDSLINIS-LNVPES-MIYQ-IPFYIKKLPLFNIILGM
MFSAILILYLKK---LKD------FFKEKLKYIVLFLRKSWFFDELYDFLFVKPSKYIG-
NGFWKAIDIELIDNVGPNGMARIVKKMGSIVSSLQTGFLYHYAFMVIIGLTIFISIYFY-
---LF----------
>WP_041471749_Rickettsia_conorii_o_Rickettsiales
--MYQNICIMMIIMLPLASSIINGLF----LRVIDKKLAQVIATGFLSLSALFSLIIFCD
TGL--DGNIIHIKL-LPWIEVGTFKVNWSIYIDQLTSIMFIAVTWVSSIVHIYSLGYMAE
DKGIIRFLSFLSLFTFFMLMLVSSDNFLQLFFGWEGVGVCSYLLIGFWYSKESANKAAIK
AFIINRASDFAFILGVITIIVYCGSANY---KDVLSSAELLS---NI--KI-FL--HFSI
LDIICLLLFIGCMGKSAQIGLHVWLPDAMEGPTPVSALIHAATMVTAGVFLVARCSYLF-
EYSPLILQFITIIGGVTCLFAASIAIMHSDIKKIIAYSTCSQLGYMFMACGVSAYNSGIF
HLVTHAFFKALLFLSAGSVIHAVH-EQDIFKMGDLRNKMPVTYGNFLIGSLALI-----G
IYP-----LA---GFYSKDSILEAAYSS----GSFMFIFGIAAAILTAIYSMKIIMLVFH
GKTK------------LEKDVFEH--------------A-HEPAKVMNNPLILLVVGSFF
```

```
SGMIGYYLLAM---DKPNGYF-HASLFNLH-IYKLL---ISH-PPLYIKLLPMAVGIVGI
VTGIYLYKSSTVM-SFPQ------KILRSSRGMTPLVLNKYYFDEIYNCLIVKPINCLA-
SLFY-LGDQQIIDRFGPNGFSRVVNCFSVLTGKIQTGYVFNYALYIVSFIVVTISYFVW-
---KNIMY-------
>ATY40894_ND5_Picozoan_Picobiliphyte_MS584-11
-------MYLTIVFLPLVGSMIAGLG----GRWVGPIGACLVTTTCLLISFFFSCIVFYE
VGL--CGSPCHIHL-LDWFDSAAFVGAWGFQFDTLTATMLIVVTGVSSCVHIFSVDYMSG
DPHRARFMSYLSFFTFFMLTLITADNFIQLFFGWEGVGLCSYLLINFWYSRITANRAALK
AFIMNRVGDFGFGLGIFGCYMVFQSIEF---ETIFACAPLYA---NQ--TFEFFNFEVDQ
LTCIVCLLFVGAVGKSSQIGLHTWLPDAMEGPTPVSALIHAATMVTAGIYMLLRCSPLL-
EYAPDALSVIAFFGASTAFFAATTGLLQNDLKKVIAYSTCSQLGYMCFAVGLSKYSVSFF
HLANHAFFKALLFLSAGCVIHAMQDEQEMRRMGGLLSSLPFTYSTMTLGSLALM-----G
TPF-----LS---GFYSKDAILEYASIHYFIAGSFTFWLGVLAAFCTAFYSFRLGYMTFL
TNTN------------AYRYCVEH--------------I-HDAPGIVLVALCPLAVGSIW
SGYFGADMFLG---VG-TPFW-GNALFFGW-TTTLTL-EPHF-LPMATKFLPLVFTILGG
FLAFLAFNYLANI-TFTL------TFHPYIKPIFAFLNKKWYFDKIYDEVFLVQSMLFGR
NVTYRLIDGFVFEQLGPMGIQKTIGVLSGSHKRFHNGFIPNYTSIFFFGILLFLVGFVAL
P--KCLVLYPGL---
>NP_062497_ND5_Chondrus_crispus
-------MYLLILFLPLLGSLISGFG----GRWLGCRGTNTFSTLCVVVSSLFSLLAFFE
IGL--TNTTCTIFL-VSWIKSGAFYVSWGFLFDSLTVTMLVVITLVSSLVHIYSIKYMEN
DPHQPRFMSYLEIFTFFMLILVTADNLIQMFLGWEGVGLASYLLINFWYTRLAANQSAIK
ALIVNRVGDFGLSLGIFLIFWVFNSVDY---SVIFSLVPLFD---NQ--FLTFLGFKLHV
LTLISLFLFIGAIGKSAQLGLHTWLPDAMEGPTPVSALIHAATMVTAGVFLMIRFSPLL-
EFSPTILFILITFGSLTAFFAAVTGVFQHDLKRVIAYSTCSQLGYMIFSCGMSCYDVSLF
HLANHAFFKALLFLSAGSVIHAVSNEQDMRRMGSLLKFMPLTYSVMLIGTLALI-----G
FPF-----LT---GFYSKDFILELTSSLQMSYISFACWLGTMSVFFTSFYSFRLIYLTFL
NNTN------------LAKSSLNL--------------V-HESSLLMIFPLIILSIGSIF
AGYLIRDLFVG---SG-SDFW-GAAIFILP-KHSTFI-EAEE-LPIVVKWLPFILSLLGI
FFASFVQIFLKTF-YFKS------NLQNLLSFFTFLINKKWYWDVLYNRLIVLPILNFGY
SISFKILDRGFIELSGPYGFTKFVSFWSQILIKLQTGQITHYLFFMIFTFCSFSIILVY-
---SYINLTFN----
>QKZ95183_NAD5_Pyropia_pulchra
-------MYLLIVVLPLVGTLVTGLG----GRWIGRKGANLFSTTCVILCCCLSIVAFFE
VGL--CGVPCYLSL-SPWISSGALKISWGFLFDSLTTTMLVVITSISSLVHLYSIQYMEY
DPHCPRFCPSWKFFTFFMIVLVTADNFVQMFLGWEGVGLASYLLINFWYTRLCANQAAIK
ALVVNRVGDFGLSLGIFTIFFLFGSVDY---ELVFASASLYT---NY--SIYFLGCSVNF
LTIIGIFLLIGAIGKSAQLGLHTWLPDAMEGPTPVSALIHAATMVTAGVFLIVRCSALI-
NLSSNVLFLITILGSSTAFFASIVGVFQNDIKRVIAYSTCSQLGYMLFVCGLSYYNVGMF
HLVNHAFFKALLFLSAGSVIHALSNEQDMRRMGSLANSLPITYAAMLIGSLSLA-----G
FPF-----LT---GFYSKDLIIEITSSLQITFGIFACWLANISVFFTAFYTFRLLFLTFV
KNSN------------SYEKNIEN--------------I-HESPILILIPLILLSLASIF
VGFLTKDLFVG---VG-TSFW-GNAINILP-TSCNLL-EVEF-MSYLIKWLPFVLSINGA
IFAYTLNIGYYKN-NIQF------AYNHIFRKLAFSLSKKLYWDKLYNFLVVSPLMQFGY
NVSFKNIDRGFIELLGPYGISRTIKKWSTQILKIQTGQVTHYTFFVVSGLCLFFLFAPVW
---SSLEFFIDIR--
>YP_006665880_NAD5_Porphyra_umbilicalis
-------MYLLIIALPLIGTLFTGLG----GRWLGRKGSNIFSTTCVILCFFLSLLAFFE
```

```
VGL--CGVPCYVSI-SPWISSGVLNITWGFLFDSLTTTMLVVITSISSLVHLYSIQYMEY
DPHCPRFMSFLEIFTFFMLLLVTADNFVQMFLGWEGVGLVSYLLINFWYTRLCANQAAVK
ALIVNRVGDFGLSLGILTIFSLFGSVDY---EVVFSLVHTYS---NH--NIYLFGAHLNT
LTLVGIFLLIGAVGKSAQLGLHTWLPDAMEGPTPVSALIHAATMVTAGVFLIVRCSPLI-
DLAPNVLFLITILGSSTAFFASIVGVFQNDIKRVIAYSTCSQLGYMVFVCGLSYYNVGMF
HLVNHAFFKALLFLSAGSVIHALSNEQDMRRMGSLMHNLPITYAAMLIGSLSLA-----G
FPF-----LT---GFYSKDLIIEITSSLYITYGIFACWLANISVFFTSFYTFRLIFLTFI
KNNN------------SYRTYMDG--------------I-HESPKLILIPLILLAIASIF
VGFISKDLFVG---VG-NSFW-GNSISTMP-IACNLL-EVEF-MTTSIKWLPFVLSTLGA
FFAYSVNAGIFKN-NITF------AYSNRFRKLAFSLSKKLYWDKLYNGLIVSFLLNFGY
NISFKNIDRGFIELLGPYGISKIISNLSFKIAKIQTGQITHYTFFVVTGLCLLLLLVPFF
---TFLENLIDIR--
>AIU44693_NAD5_Cyanophora_paradoxa
-------MYLTLIILPLLSALV-GFL----VYFIGNRFAAFICTTLLGFTLALSIKSFYY
IAL--LQNPCYIQT-LPWFLGENLVIFWGFLFDSVTATMLVVVTSISFLVHFYSIEYMGA
DPHLGRFMSYLSFFTFFMLILVTADNFVQMFVGWEGVGLCSYLLINFFFNRIQANKAAIK
AMIMNRIGDFGLSLAIMVIFYTCKSVDY---HTVFACVPFFI---DS--TFMFFNFEVNL
ITCICILLFIGAVGKSAQIGLHTWLPDAMEGPTPVSALIHAATMVTAGVFLIIRCSFLF-
EFSDTALDILTIIGALTAFFAATTGLLQNDAKRVIAYSTCSQLGYMVFVCGFSGYSVSLF
HLANHAFFKALLFLTAGSLIHGMQDEQDFRKMGGLSYLLPYSYSMLLIGSLSLA-----G
FPF-----LT---GFYSKDMILELTFSSYLFKGTFAYTLGVVSAFFTAFYSFRLIYWAFL
AKPN------------GYKYSYAN--------------A-SESRFFITIPLAILAFASIF
IGYILRDMFIG---VG-TSFW-NNSIFILP-ENNFIL-ESEF-IPIHNKLTPVIFSLCGL
FTAFFLYHILFKK-IFIL------NSPYI--KFYTFLNKRWYFDKIYNEFIALPVISSGY
KITFKIIDRGILELFGPAGIVANLLNIARNTSNLQSGFMYHYLFLILLSITGAIVLIIGL
LY-NYFTLPIG----
>YP_009092462_NAD5_Gloeochaete_wittrockiana
-------MYLVIVFLPLVSAIMAGLF----GRFLGNKNSAYLATFCLGFTFFLCLIAFYE
VGL--CHSPCYIVS-FPWIQSELFKVNWSFLFDSITVVMLIVVTSVSFLVHMYSIEYMGQ
DPHLSRFMSYLSFFTFFMLILVTADNFLQMFVGWEGVGLCSYLLINFYFTRVQANKAALK
AMIVNRIGDFGLSLGMMALFFTFKTLDY---NVVFNLAPLVV---NE--NFCFFSFELNK
ITVICLLLFVGAVGKSAQLGLHTWLPDAMEGPTPVSALIHAATMVTAGVFLIIRCSPLF-
ELSAIALDVLMVFGSLTAFFAATTGLLQNDVKRVIAYSTCSQLGYMVFACGASSYSVAMF
HLANHAFFKALLFLTAGAIIHALRDEQDMRKMGGLAMLLPFAYSMIVIGSLSLM-----G
FPF-----LT---GFYSKDSILELVYSSYSSPAGFAYVLGTLAAFCTAFYSFRLIYLVFL
SVPN------------GYKSYFNQ--------------V-HETPYLMALPLVLLAFGSIF
IGYLTKDLMLG---MG-TNFW-GGSIYFLP-ERAGLF-EAEF-IPTNIKILPVCLSLLGA
FTACLFYSSFFK---FLV------YLSVYVKDFYTFLNKKWYFDKIYNEVVGKFFIWFGY
NMSFKLIDRGFVELFGPYGMSKLSFNLAKQFSLLQTGFLYHYIFMLLVGVTFFVGILSFG
---YYLNFWVDTR--
>YP_004222736_NAD5_Glaucocystis_nostochinearum
-------MYLSIVFLPLLSAFVAGLL----GKFLGNKISSYFTTICLGITFILSIIAFYE
VAL--CASPCYLTT-ISWIKSELFHVNWGFLFDTLTVVMLIVVTSVSFLVHMYSIEYMSH
DPHLSRFMCYLSFFTFFMLILVTADNFLQMFVGWEGVGLCSYLLINFYFTRIQANKSAIK
AMIMNRIGDFGLSLGMMAIFLTFKSLNY---DIIFTSVNLYS---HE--FFLFLNFEVNK
ITLICILLFVGAVGKSAQVGLHTWLPDAMEGPTPVSALIHAATMVTAGVFLIARCSPLF-
EYSTLALSILTIFGATTAFFAATTGLLQNDIKRVIAYSTCSQLGYMVFSCGISSYSAAVF
HLANHAFFKALLFLTAGSIIHSFQDEQDMRKMGGLGLILPYSYSMILIGSLSLM-----G
```

```
FPF-----LT---GFYSKDIILELAYSTYSIDSTFAYWLGTISAFFTAFYSFRLIYLVFL
NKPN------------GYKNAYQN--------------A-DDSHFWIALPLSLLAFGSIF
IGYLSKDMMLG---LG-TNFW-GNSLFLLS-NHTTIL-DSEY-IDYYLKLIPVIFSLIGS
VCSYLLYRFSKN---FLF------TLNLRFKTIYIFFNKRWLFDKIYNEFIGLPALSFGY
NISFKLLDRGFFEIFGPYGLAFVLLNFAKNTSRLQTGLLYHYIFIFILGLTFFVGLIKFG
EQ-FFIYAFI----F
>YP_009317212_NAD5_Palpitomonas_bilix
-------MYLLIVALPLISALTSGLF----GRFLGSKGAGFISVCCLIATSFLSFIAFYE
VAL--NGSPCYIKL-ATWVDSEMLHADWGFVFDSLTVIMLVVVTFVSALVHLYSTGYMEG
DPHVPRFMSYLSLFTFFMVMLVTADNFIQMFLGWEGVGLCSYLLITFWFTRVQANKAALK
AFIVNRVGDFGLSLGVFAIFYLFSSLDF---STVFALAPYMV---GS--QLIFCGFEVDS
LTLICILLFVGAVGKSAQLGLHTWLPDAMEGPTPVSALIHAATMVTAGVFMIARCSPLF-
EFAPTALFVVAIVGAMTAFFAATTGMLQNDLKKVIAYSTCSQLGYMVFVCGLSNYSVGVF
HLANHAFFKALLFLSAGSVIHAMSDEQDMRRMGGLAQIIPFTYVMMVIGSLSLM-----G
FPF-----LT---GFYSKDVILEIAYAKYSFAGTFSHWLGCISAFFTAFYSFRLLYLTFI
QNTN------------AYKRVIEH--------------A-HESSFVMLLPLILLSFGSIF
VGYLTKDMIIG---LG-TSFW-GNSIFTLP-TNITMV-DAEF-LPYYIKLIPVILSLSGA
VISLLVYHFMSQI-LFNV------QTSFIGRHLYIFFNKKWFFDKIYNYFFVQHVLRFGY
SITFKSLDKGLIEFLGPYGIANVVSRVSSLTSALHTGYIYHYAFVMFVGVVFILRIIF--
---------------
>YP_007890522_NAD5_Andalucia_godoyi
-------MYLLIVFLPLLGSILAGLF----GRFLGRHGASLVSTTSVGLTCLFSWIAFYE
VGI--CESPVYLTL-LPWIDSGMLTANWGFLFDSLTVVMLIVITTVSFLVHIYSTSYMSE
DPHLPRFMSYLSFFTFFMLMLVTADNFVQMFLGWEGVGLCSYLLINFWFTRLQANKAAIK
AMIMNRIGDFGLSLGMMALFAVFQALDF---STLFAIAPLFA---NQ--DFLFLNFQVDL
LTTICILLFVGSVGKSAQLGLHTWLPDAMEGPTPVSALIHAATMVTAGVFLIVRCSPLF-
EYAPSALLVVTFVGAMTAFFAATTGMLQNDLKRVIAYSTCSQLGYMVFACGLSSYSVSMF
HLMNHAFFKALLFLSAGAVIHALADEQDMRRMGGLVQLLPFTYSMMLIGSLALM-----G
FPF-----LT---GFYSKDVILELAYAKYSLDGTFAHWLGTVSAFFTSFYSFRLLYLTFL
TKTN------------AYKSSVSH--------------A-HDAPFVMAFPLMVLALGSLF
VGYLFRDMMIG---LG-TSFW-GNSLFVLP-EHLILI-DSEF-IPYSIKSIPVLLSFVGA
SLAWLLNHSYGAF-LYEL------KTSSFGRDLYTFLNKRWLFDKVYNDYVGKVLLQFGY
NVSYKTLDKGILEILGPYGIIRLVRHLSDRVSSFHTGYLYHYAFIMLIGVTLLISVIRFW
---DVLSQVVDPR--
>AGH24310_NAD5_Reclinomonas_americana
-------MYLLIVFLPLLGSITAGFF----GRSLGKQGAAIITTSCVALSSLFSMVAFYE
VGL--CGSPCYIRL-FNWIDSEMLHASWGFLFDSLTVVMLIVVTIVSSLVHLYSVGYMSH
DPHLPRFMSYLSLFTFFMLMLVTGDNFVQMFLGWEGVGLCSYLLINFWFTRLQANKSAIK
AMIMNRIGDFGLSLGMMAIFFIFKSVDF---ITVFALSPYMT---DA--TIVFLNYEVHA
LTLICILLFVGAVGKSSQLGLHTWLPDAMEGPTPVSALIHAATMVTAGVFLIARCSPIF-
EYAPTALLVVTIVGAMTAFFAATTGLLQNDIKRVIAYSTCSQLGYMVFACGISGYSVGMF
HLMNHAFFKALLFLSAGCVIHALADEQDMRRMGGIVKIVPFTYGMMLIGSMSLM-----G
FPF-----LT---GFYSKDVILELAFAKYTIDGTFAHWLGTVAAFFTAFYSFRLIYLTFL
GETN------------APRTIINH--------------A-HDAPFIMALPLMILAIGSIF
VGFIMKDMMIG---LG-TDFW-GNSLFTHP-KNLTLI-ESEF-IPTPIKLLPVILSIVGA
SLAIILNNFYATF-LVSL------KTSLLGREIYSFLNKRWYFDIVYNEYVGKTLLWFGY
NISFKSVDKGLIEILGPYGLERLTRRLTSKVSALQTGYIYHYAFIMLLGVTLIITIIGLW
---DYISMWTDYR--
```

```
>YP_009121385_NAD5_Thecamonas_trahens
-------MYSLIIFLPLIGSFAAGFF----GRLIGEKGAAIITTSAMVFSTLFSYVALYN
LVV--TGNPVYIDL-FTWIDSGLFEARWGFTFDSLTVIMLIVVTTVSTLVHLYSTEYMAG
DPHVPRFMSYLSLFTFFMLMLVTSNNLIQLFFGWEGVGLASYLLINFWFTRIPANKAAIK
AMIVNRVGDFGLSLGIMAIFFIFKSVDF---ATIFALIPHFD---GY--TFVFFNDTYNI
FNVICLLLFIGAVGKSAQIGLHTWLPDAMEGPTPVSALIHAATMVTAGVFLLIRCSALF-
EYAPDVLLIVTFVGAVTALFAATAGMLQNDLKKVIAYSTCSQLGYMVFACGISNYSVSMF
HLMNHAFFKALLFLSAGSVIHAMADEQDMRKMGGIINLLPFTYTMILIGSLSLM-----G
FPF-----LT---GFYSKDVILELAFAKYTITSSFSYWFGTLAAFFTAFYSFRLVYLTFI
SNTN------------AYKKVMEQ--------------A-HDAPIRMAIPLIILLFGSIF
VGYLTRDMIIG---LG-TDFW-GNAIFILP-SNLNMI-DSEF-IPWNIKLIPVIFSLIGA
SLAFILNAFYDRF-LVDF------KLSKLGLRLYGFINKKWYVDNIYNEFIVKNFLAFGY
HVSFKTIDRGLIEYIGPYGIVKLAQKFSNTISKLQTGYVYNYTAMTLIGSILFITLIGLW
---DILNVFVDHR--
>NP_054400_NAD5_Marchantia_paleacea
-------MYLLIVILPLIGSFAAGFF----GRFLGSRGVAVVTTTCVSLSSIFSCIAFYE
VAL--CASACYIKI-APWIFSELFDAAWGFLFDSLTVILLLVVTIVSSLVHIYSISYMSE
DPHSPRFFCYLSIFTFFMLMLVTGDNFIQLFLGWEGVGLASYLLINFWFTRIQANKAAIK
AMLINRVGDFGLALGIMGCFTIFQTVDF---STIFACASAFS---EPHHYFLFCNMGFHA
ITVICILVFIGAVGKSAQIGLHTWLPDAMEGPTPVSALIHAATMVTAGVFMIARCSPLF-
EYSPNALIVITFVGAMTSFFAATTGILQNDLKRVIAYSTCSQLGYMIFACGISNYSVSVF
HLMNHACFKALLFLSAGSVIHAMSDEQDMRKMGGLASLLPFTYAMMLIGSLSLI-----G
FPF-----LT---GFYSKDVILELAYTKYTISGNFAFWLGSVSVFFTSYYSFRLLFLTFL
APTN------------SFKRDLSR--------------C-HDAPILMAIPLILLAFGSIF
VGYLAKDMMIG---LG-TNFW-ANSLFILP-KNEILA-ESEFATPTIIKLIPILFSTLGS
FVAYSVNFVVNPL-IFAL------KTSTFGNRLYCFFNKRWFFDKVFNDFLARSFLRFGY
EVSFKALDKGAIEILGPYGISYTIRKMAQQISKIQSGFVYHYAFVMLLGLTIFISVIGLW
---DFISFWVDNR--
>YP_009047585_NAD5_Sphagnum_palustre
-------MYLLIVTLPLLGSCVAGAF----GRFLSSRGTAIVTSTCVSLSFILSLIAFYE
VAL--EASACYIKM-APWIFSEMFDASWGFFFDSLTVIMLIVVTFVSSLVHIYSISYMFE
DPHSPRFMCYLSIFTFSMLMLVTGDNSIQLFLGWEGVGLASYLLINFWFTRLQANKAAIK
AMLVNRVGDFGLALGIMGRFTIFQTVDF---STLFACVSAFS---EPYHYLIFCNMKFHA
ITVICILLFIGAVGKSAQIGLHTWLPDAMEGPTPVSALIHAATMVTAGVFMIARCSPLF-
EYPPNALIVITFVGAMTSFFAATTGILQNDLKRVIAYSTCSQLGYMIFACGISNYSVSVF
HLMNHAFFKALLFLSAGSVIHAMSDEQDMRKMGGLASLLPFTYAMMFIGSLSLI-----G
FPF-----CT---GFYSKDVILELAYTKYTISGNFAFWLGSVSVFFTSYHSFRLLFLTFL
APTN------------SFKQDILQ--------------C-HDAPILMAIPLIFLAFGSIF
AGYLAKDMMIG---LG-TNFW-ANSLFVLP-KNEIIA-ESEFATPTIIKLIPILFSTLGA
FVAYNINFVANQF-IFAL------KTSTLGNRLYCFLNKRWFFDKVFNDFIVRSFLRFGY
EVSFKVLDKGAIEILGPYGISYTFRKLAKQISKLQSGFVYHYAFVMLIGLTIFITIIGLW
---DFISFWVDNR--
>NP_943684_NAD5_Chara_vulgaris
-------MYLLIVVLPLLGSLVAGVF----GRFLGSRGAALVTTTCVSISSGLCFIAFYE
VAL--GASACYIKF-APWILSEMFDASWGFLFDTLTVVMLIVVTFVSSLVHIYSISYMSE
DPFLPRFMCYLSIFTFFMLMLVTGDNLIQMFLGWEGVGLASYLLINFWFTRLQANKAAIK
AMLVNRVGDFGLALGIMGCFAIFQTVDF---STIFACASAFVW--EP--SFIFLNMKIHA
LTAICVLLFIGAVGKSAQIGLHTWLPDAMEGPTPVSALIHAATMVTAGVFMMARCSPLF-
```

```
EYAPKALIVITFVGAMTSFFAATTGILQNDLKRVIAYSTCSQLGYMVFACGISNYSVSVF
HLMNHALFKALLFLSAGSVIHAMSDEQDMRKMGGLASLLPFTYAMMLIGSMSLI-----G
FPF-----LT---GFYSKDVILELAYTKYTISGNLAFWLGSLSVFLTSYYSFRLLFLTFL
APTN------------AFKRDIER--------------C-HDAPTLMAIPLILLAFGSLF
VGYLAKDMMIG---LG-THFW-ANSIFILP-KNEIMF-SSEFATPRMMKLIPILFSAIGA
FLAYRVNFCANKF-IYVL------KTSTLGTKCYCFLNKRWLFDKVFNDFIAKSFLRFGY
EVSLKTLDKGAISILGPYGISTTFRKLAKQISTLQSGFVYHYAFVMLIGLTIFITIIGLW
---DFISFWVDNR--
>YP_009472097_NAD5_Arabidopsis_thaliana
-------MYLLIVFLPLLGSSVAGFF----GRFLGSEGSAIMTTTCVSFSSILSLIAFYE
VAL--GASACYLRI-APWISSEMFDASWGFLFDSLTVVMLIVVTFISSLVHLYSISYMSE
DPHSPRFMCYLSIFTFFMLMLVTGDNFLQLFLGWEGVGLASYLLIHFWFTRLQADKAAIK
AMLVNRVGDFGLALGILGCFTLFQTVDF---STIFACASV------PRNSWIFCNMRLNA
ISLICILLFIGAVGKSAQIGLHTWLPDAMEGPTPVSALIHAATMVTAGVFMIARCSPLF-
EYSPTALIVITFAGAMTSFLAATTGILQNDLKRVIAYSTCSQLGYMIFACGISNYSVSVF
HLMNHAFFKALLFLSAGSVIHAMSDEQDMRKMGGLASSFPLTYAMMLIGSLSLI-----G
FPF-----LT---GFYSKDVILELAYTKYTISGNFAFWLGSISVLFTSYYSFRLLFLTFL
VPTN------------SFGRDISR--------------C-HDAPIPMAIPSILLALGSLF
VGYLAKDMMIG---LG-WNFW-ANSLLVLP-KNEILA-ESEFAAPTIIKLIPILFSTLGA
FVAYNVNLVADQF-QRAF------QTSTFCNRLYSFFNKRWFFDQVLNDFLVRSFLRFGY
EVSFEALDKGAIEILGPYGISYTFRRLAERISQLQSGFVYHYAFAMLLGLTLFVTFFCMW
---DSLSSWVDNR--
>YP_588334_NAD5_Zea_mays
-------MYLLIVFLPLLGSSVAGFF----GRFLGSEGTAIMTTTCVSFSSILSLIAFYE
VAL--GASACYLRI-APWISSEMFDASWGFFFDSLTVVMLIVVTFISSLVHLYSISYMSE
DPHSPRFMCYLSIFTFFMLMLVTGDNFLQLFLGWEGVGLASYLLIHFWFTRLQADKAAIK
AMLVNRVGDFGLALGIFGCFTLFQTVDF---STIFACASA------PRNEWIFCNMRFNA
ITLICILLFIGAVGKSAQIGLHTWLPDAMEGPTPVSALIHAATMVTAGVFMIARCSPLF-
EYSPTALIVITFAGAMTSFLAATTGILQNDLKRVIAYSTCSQLGYMIFACGISNYSVSVF
HLMNHAFFKALLFLSAGSVIHAMSDEQDMRKMGGLASSFPLTYAMMLMGSLSLI-----G
FPF-----LT---GFYSKDVILELAYTKYTISGNFAFWLGSVSVLFTSYYSFRLLFLTFL
VPTN------------SFGRDRLR--------------C-HDAPIPMAIPLILLALGSLF
VGYLAKDMMIG---LG-TNFW-ANSPFVLP-KNEILA-ESEFAAPTITKLIPILFSTSGA
SLAYNVNLVADQF-QRAF------QTSTFCNRLYSFFNKRWFFDQVLNDFLVRSFLRFGY
SVSFEALDKGAIEILGPYGISYTFRRLAERISQLQSGSVYHYAFAMLLGSTPFVTFSRMW
---DSLSSWVDSR--
>AJF36693_NAD5_Klebsormidium_flaccidum
-------MYLLIVTLPLLGCATSALF----GRFLGSRGAAIVTTSCVGTSFLFSLVAFYE
VAL--VGSACTLQF-APWFDSEMFDASWGFLFDSLTVVMLIVVTLVSTLVHTYSISYMSE
DPFLPRFMCYLSIFTFFMLMLVSADNLIQMFLGWEGVGLASYLLINFWFTRLQANKAAIK
AMLVNRVGDFGLALGIMSCFLVFQSVDY---HVLFAVANEWAMKESE--SFVFCNMHVDA
LSAICLLLFVGAVGKSAQIGLHTWLPDAMEGPTPVSALIHAATMVTAGVFLIARCSPLF-
EYAPNALVVVTCMGAMTAFFAATTGILQNDLKRVIAYSTCSQLGYMVFACGISNYAVSIF
HLMNHAFFKALLFLSAGSVIHAMSDEQDMRKMGGLNQVLPFTYAMMLIGSLALI-----G
FPF-----LT---GFYSKDVILELAYTKYTISGHFAFWLGSLSVLFTSYYSFRLLFLTFL
ENTN------------AFKQDIKN--------------A-HDAPPLMAAPLIVLAFGSLF
VGYLCKDMMIG---LG-TDFW-GQSIFVLP-DNSNLS-ESEFSTPQIVKLIPLLLSTLGA
GIAYWINNHKNSF-VYDF------KTSSLGRRFYTFLNKRWLFDKVLNEFLAQSVLRFGY
```

```
SISFVTLDKGVIELLGPYGIATTMRTLSKRTSGLQSGLIYHYAFIMLIGLTVLITFIGLW
---DIVSFWVDSR--
>QIQ59668.1_Nad5_Trebouxia_sp._A1-2
-------MYLSLIFLPLLGSLCAGFF----GRFLGFRGACLISTTSVFSSFIMSTVAFYE
VAL--SGSSCYIKY-SSWFVSEMFDSSWGFYFDTLTVVMLVVVTSVSTLVHLYSISYMSG
DPHLPRFMSYLSIFTFFMLVLVTADNFLQMFFGWEGVGLASYLLINFWFTRLQASKASIK
AMLVNRVGDFGLSLGIMAIFSVFKSVDF---ATVFSCAPHFA---DT--EFLFCNIECTL
LNVVCILLFIGAVGKSAQLGLHTWLPDAMEGPTPVSALIHAATMVTAGVFLIARCSPLF-
EYAPNALMIVTLVGGMTTFFAATTGIVQNDLKRVIAYSTCSQLGYMVFACGISNYSVGVF
HLMNHAFFKALLFLSAGSVIHALSDEQDMRKMGGVVQLLPFTYGMMLIGSLSLT-----G
FPF-----LT---GFYSKDTILELAFASYTVNGNFAYWLGAICVLFTSYYSFRLLFLTFL
SPTN------------LYKSSLKN--------------T-HDANFIMALPLILLAFGSIF
VGYLGKDMMIG---LG-TNFW-GNALFVLP-KNINLL-ESEY-IPQVQKMVPLFFTLLGA
FLAYLVNYSLFRE-TYFI------KTSWIGRKFYYMLSKRWLFDKVYNDFIGQKNLDFGY
RISFKTLDKGCFEILGPYGISFLFQNLTRYLSKLQSGMIYHYAVVMLLGVILLISIVGLW
---EFLEVFVDNR--
>YP_717300.1_nad5_Ostreococcus_tauri
-------MYLLLVFLPFFGFLLVTVF----GRFFGYRGAPLLSTFAVMSSACLSFVALYE
VGI--CGSPCYVQF-LPWFQTDMFVASWGFLFDSLTVIMCVVVTFVSSLVHIYSISYMGE
DPHLPRFMSYLSVFTFFMLMLVTSDNFLQMFFGWEGVGVASYLLINFWYTRLQANKSAIK
AMLVNRVGDFGLALGIMGIFHIFKAVDF---ETVFACASEFA---CH--HFLFFHMEVHT
LTCISLLLFIGAIGKSAQLGLHTWLPDAMEGPTPVSALIHAATMVTAGVFLLARCSPLV-
EYSSQALVVITLVGASTAFFAATTGVVQNDLKRVIAYSTCSQLGYMMFACGISQYALGVF
HLMNHAFFKALLFLSAGSVIHALGDEQDMRKMGGLVRLLPFTYSMMMLGSLALI-----G
FPY-----LT---GFYSKDVILEVAYAKYTLSGNFAYWLGSVSALFTSYYSFRLLYLTFL
ASPQ------------MLKSSVQN--------------V-HDAPFLMGLSLCCLALGSIF
GGFIAKDMMIG---LG-TNFW-GNAVFTLP-TNTLVL-ESEY-IPQAVKFVPIIFSILGA
VFAVNLNGFGSSF-SYSL------KTSVLGKQLYTFLNKRWLFDKVYNDFVARPSLHFGY
TISFKLVDKGFLEMFGPTGIMQQSAFMRHFVKTVQTGSIYHYAFMMFVSLTSLAFFLVAT
---TSVGAFLDTR--
>YP_008994799_NAD5_Bathycoccus_prasinos
-------MYLTILFLPLLGFLLAASG----GRFFGWRGTPLITTFCVFLSSLFSFIAFYE
VGI--AGSPCYIQL-APWFTSEFFDATWGFMFDSLTVVMLVVVTFVSTLVHIYSISYMSE
DPHLPRFMSYLSIFTFFMLMLVTADNFIQLFFGWEGVGLASYLLINFWYTRLQANKSAIK
AMLVNRVGDFGLALGIIATFSLFKSVDF---ATVFACSAHFA---EN--SFIFFHLEWHA
LSLICALLFVGAVGKSAQLGLHTWLPDAMEGPTPVSALIHAATMVTAGVFMIARCSPLF-
EQAPQTLILVTVTGALTAFFAATTGVVQNDLKRVIAYSTCSQLGYMVFACGISQYAVGVF
HLMNHAFFKALLFLSAGAVIHALADEQDMRKMGGTIKVLPFTYSMMFIGSLALI-----G
FPF-----LT---GFYSKDVILEVAYAKYTVAGTFTYWLGAVSALFTSYYSFRLLFLTFI
GPVN------------SLKDSLKH--------------V-HDAPFLMGFPLFLLAFGSIF
VGYLAKDMMIG---MG-TQFW-GNALFMGP-DHGLLV-ESEY-IPQAIKYIPILFSFLGF
LFALNLNLLGASF-SYSL------KTSTFGRTLYIFLNKRWLFDKMYTDFVALPALFFGY
NVSFKTLDKGLLEIVGPAGVIQTVSQGGRQLRKMQSGLIYHYAFIMLCSLTFLIAFVGLW
---DFFSFFVETH--
>YP_009446429_NAD5_Ancoracysta_twista
-------MYLLIVFLPLLSALTVLSF----GRYLGREGSVLLTVSSLFITAAMSTTAFYE
VAL--CGSVCHIKL-APWITSELLEISWGFLFDSLTAVMLVVVTYVSALVHLYSVSYMEE
DPHLPRFMSYLSLFTFFMLMLVTGDNFLQLFLGWEGVGLASYLLINFWYSRIQANKSAIK
```

```
AMVMNRIGDFGLALGLMAIFAVFKSIDY---STVFATAPFFS---NE--NFLFCNLELDQ
LTVICLLLFVGAVGKSAQLGLHTWLPDAMEGPTPVSALIHAATMVTAGVFMICRCSPLF-
EYAPDALFVITLVGASTAFFAATVGVVQNDLKRVIAYSTCSQLGYMVFSCGLSSYSVSMF
HLMNHAFFKALLFLSAGSVIHAMGDEQDMRRMGGLLNLLPFTYTMMLIGSLSLV-----G
FPF-----LT---GFYSKDVILELAYGKYTVSGTFAHWLGSLSVFFTSFYSFRLIYLTFI
NKTN------------SHKSLIEG--------------A-HDAPIIMAIPLMILAVGSIF
VGYVTKDLMIG---LG-TTFW-SHSLFVLP-VNMTSV-ESEF-LDASIKCIPLFFSGMGA
TVAFILCYANRRL-NFRL------TSNPVGRGIYTFLNKRWLVDAVYNVYGAAPLMRFGY
STSFRLLDTGFIQMFGPRGLSTLLLELSSLASRFQSGYVYHYAFIMVCGATILVGFINLW
---PFTSLYLSNK--
>BBD14148_NAD5_Ophirina_amphinema
-------MYLLIVFIPLVTSLVSGLG----GRFIGKLGSQVLTSLGVLMSATLSSVAFYE
VGI--CGSPCYVEF-LPWISSDMLRVYWGFQFDSLTVVMLVVVTYVSTLVHLYSIGYMSE
DPHIPRFMSYLSLFTFFMLMLVTGDNLVQLFFGWEGVGVCSYLLISFWYTRIQANKAALK
AMVMNRVGDFGLSLGIFASFYVFKSLDY---ATIFSLAPYYV---DF--QIPFLYWSFGS
LNLIGILLFVGAVGKSAQLGLHTWLPDAMEGPTPVSALIHAATMVTAGVFLISRCSPLY-
EYAPKALLVVTIIGGTTAFFAATTGLVQNDLKRVIAYSTCSQLGYMIFACGLSGYSIGMF
HLMNHAFFKALLFLSAGSVIHALADEQDMRKMGGMLNIVPLTYTMMLIGSMSLM-----G
FPF-----LT---GFYSKDVILEFSFAKYTIDSSFVFWLGVISAFFTAFYSFRLIYLTFF
GEPN------------SIRSVVSK--------------S-HDAPVIMALPLMLLCLGSVF
VGYIFRDLIIG---LG-TDFW-GSALYVSP-HNSAHI-DSEF-IPHMVKLMPVCLSVLGA
LASLYVYEKLALE-IGSL------YKTAIWLSVYKFLNGRWLFDSVYNYFVAEKLMSFGY
HISFKTLDKGIIEFMGPSGLISLISTWSIRVSRLNTGYLYHSAFLMMLGAVIVLGVTRVC
---QMESDIA-----
>ANA57055_NAD5_Pyramimonas_parkeae
-------MYLLLVVLPLMGSLFAGFF----GRYLGYRGAGLFSSTCVGLSAIGSWVAFYE
VGL--SGSPCYLKF-APWFHSEFFDASWGFLFDSLSVVMLVTVTTASTCIHLYSISYMGE
DPHLPRFMSYLSIFTFFMLVLVTADNFIQMFLGWEGIGLASYLLINFWVTRLQANKAALK
AMLVNKIGDFGLALGILGVFQLVKTVDY---ASLFACVPYIN---ES--SICFFQWELDG
LTLIACLLFLGVVGKSSQIGLHTWLPDAMEGPTPVSALIHAATLVTAGVFLMARISPLL-
EYAPQALSLIAFTGAMTCFLAASTGIMQNDLKRVIAYSTCSQLGYMVFACGLSSYGVSIF
HLMNHAFFKALLFLSAGSLIHALADEQDMRKMGGLGRLLPYTYGMILIGSLSLA-----G
FPF-----LT---GFYSKDVILEMAYAKYSLSGNFAYWLGALSVFFTSYYSFRLISLAFF
APTN------------CIKGSIRT--------------V-HDAPFLLGAPLIPLALGSLF
LGFLGKDMMIG---AG-STFW-GNALFILP-QNSLSI-ESEW-IPQNVKLLPLVLTIGGA
ITAYLINLVWIES-AFKI------KTHWLGREIYLFLNKRWLFDKVYNEFIAKTFMQFGY
NVSFKSLDKGAFEIIGPYGLIQSVPILLKETSRFQSGFLYHYALIMLIGITFLIGLIGLG
---PILNYWIDPR--
>YP_009476712_NAD5_Proteomonas_sulcata
-------MYLTIIFLPLVGCFFSGFF----GRFLSPKGSAFITILGIFIAFIFSLVAFFE
VGL--SLCPCYIQL-LPWIDCELFDASWGFMFDTISVGMCVTVTFISLLVHIYSTSYMSH
DPHQPRFMAYLSLFTFFMLCLVTADNFIQLFLGWEGVGLCSYLLINFWFTRLQANKAAIK
AMIINRIGDVGLALGVFAAFIVFKTTNF---SVIFSLVPYYV---NH--EIFFFTYKCNA
ITLIGLLLFIGSVGKSAQLGLHTWLPDAMEGPTPVSALIHAATMVTAGVFLIIRCSPLY-
EYSESALFIIVIFGALTAFFAATIGLLQNDLKKIIAYSTCSQLGYMVFACGLSSYSVSMF
HLVNHAYFKALLFLSAGSVIHALNNEQDMRRMGGLLQVLPFTYCMFVVGSLALM-----G
FPF-----LT---GFYSKDAILEIAYAKYNFIGTFAHWLGIISAFFTAFYSFRLIYFTFV
ISPN------------GYKTNMQN--------------A-HDAPLLMGLPLFLLSVASIF
```

```
IGYLTRDLYIG---LG-TPFW-NNAIYTAP-QNLSYI-DAEF-IPTSIKWLPVIFSLLGA
ILSIFLYHNCKQI-LYNL------TSISLFRYIYAFLNRRWLFDRLQNDFIGDSFLKIGY
SITYKKLDKGIIEVIGPKGIINLTTNIGKQVISLQSGLIHQYASLIFVGVIFIVIKITNF
---NFFHPFIDIH--
>YP_009050520_NAD5_Sargassum_fusiforme
-------MYLNIVFLPLLGSIIAGFF----GRFVGVYGSCFITVFCVGLTFCLSCLSFYE
VGL--AGSGTYLSL-MTWLESDSFYVEWAFCFDSLTVVMLIVVTFISTLVHIYSTEYMGG
DPHLPRFVSYLSLFTFFMLILVTADNFVQMFVGWEGVGLCSYLLINFWFTRIQANKAAIK
AMIVNRIGDFGLALGIFAIFICFGALDY---ATVFAFSPQLT---SC--TLSFLNIKFKA
LDLIGVLLFIGAVGKSAQLGLHTWLPDAMEGPTPVSALIHAATMVTAGVFLLARCSPLL-
EYCSGALLLISVVGGMTAFFAATTALVQNDLKRVIAYSTCSQLGYMVFACGLSNYSVAVF
HLANHAFFKALLFLSAGSVIHAVGDEQDMRKMGGLRRLLPFTYAMMVIGSLALM-----G
MPF-----LT---GFYSKDVILEIAYATYSIPAHFSYWLGSLAAFCTAFYSIRLIALCFL
VEPN------------GSRKLLLS--------------A-SEGGKAIGFVLGLLAIPSIF
VGYFSRDLFIG---LG-THFW-GSSLFCLP-SNLSLI-DAEF-MSTFSKMLPLCLSIVGG
ALSFTLYRYYNYN-MYFW------KTSLVGRKFYTFLSRKWFFDKIYNGFISQNILDFGY
HFTYKSIDRGLIESLGPFGLSNVLTSQVQQARVWHSGYLYHYLVIFVWGNLLFGLWFLFG
F--PSLRI-------
>YP_009254724_NAD5_Ectocarpus_siliculosus
-------MYLNIVFLPLLGSLVAGFF----GRFLGGRGSGLITISCVGLSFFLSGLAFYE
VGL--GGSSTYLYL-MPWLESDSFHVDWAFCFDSLTVVMLIVVTFISTLVHLYSTEYMGG
DPHLPRFMSYLSLFTFFMLILVTADNFVQMFVGWEGVGLCSYLLINFWFTRIQANKAAIK
AMLVNRVGDFGLALGIFGVFICFGAVDY---ATVFSLAPQLS---NF--TLLFFNFEFNA
LNLIGILLFVGAVGKSAQLGLHTWLPDAMEGPTPVSALIHAATMVTAGVFLLARCSPLL-
EYCPDALFIISIIGGMTAFFAATTALVQNDLKRVIAYSTCSQLGYMIFACGLSNYSVAVF
HLANHAFFKALLFLGAGSVIHAVGDEQDMRKMGGLRRLLPFTYAMMVIGSLALM-----G
MPF-----LT---GFYSKDVILEVGYATYSSASHFAYWLGSFAAFCTAFYSIRLLALCFL
SEPN------------GSRTFLLS--------------A-SEGSWPMGIVLGILAVPSMF
IGYFSRDLFIG---LG-TDFW-GNALFFLP-SNISII-DAEF-MSTGIKILPLCLSILGG
FSSFILYRNYSRN-LFLL------KTSILGRKLYTFLNRKWLFDKIYNDFITQNTLDLGY
HFTYKSIDRGLIESLGPFGFSNVLLDQVNKSRLWHSGYLYHYLVILGWSNLLFGVWFIFG
F--KFLPI-------
>YP_316588_NAD5_Thalassiosira_pseudonana
-------MYLLLIFLPLIGSICAGLF----GRILGPFGSSFVTVICLMVTFCLSLFAFYE
VAL--LDCCVYIKL-APWINSEMLNVDWGFLFDSLTVVMCCVVTFVSSIVHLYSTEYMAY
DPHLPRFMSYLSLFTFFMLILVTADNFIQMFVGWEGVGLCSYLLINFWFTRIQANKAAIK
AMILNRIGDFGLVIGILIIFVEYKAVDY---ATVFALTPIFT---NK--VFHFLNFDFDL
ISLICFFLFIGAVGKSAQLGLHTWLPDAMEGPTPVSALIHAATMVTAGVFLIARTSPLF-
EYTSSILSLVTIVGACTAFFAATVGLLQNDLKRVIAYSTCSQLGYMVFACGLSNYSVGVF
HLVNHAFFKALLFLGAGSIIHAVADEQDMRKMGGLKKLVPFTYSMMVIGSLALI-----G
FPF-----LT---GFYSKDVILEVAYGKYTLEGHFSYILGTLGAFLTAFYSTRLIYLTFL
SKPN------------GYKSVICA--------------A-YDSSYQISASLFFLVIPSVL
IGFYAKDMIIG---FG-SDFW-GNAIFTSI-ENMNRI-DSEF-ITHFYKILPVALSILGV
TSSFLLYLFGSKI-LVKL------KLSTIGKKIYNFFNKKWFFDKVYNEYVSQFFFTISY
TVTYKTIDRGIIEIFGPMGLSSAITKKALYISKLQTGYLYHYTFLILTGLTLILGVRQFW
---VFLGDFIDFK--
>YP_005090357_NAD5_Phaeodactylum_tricornutum
-------MYLLVVFLSAIGSCFAGLF----GKHLGPLGSIFVTTSCLFLSLFISFFIFYE
```

```
VAL--SNSVVYIKL-TTWISSEVLHVDWGFMFDSLTATMCIVVISISFLVHLYSVEYMSH
DPHLPRFMSYLSLFTFFMLILVTADNFIQMFVGWEGVGLCSYLLINFWFTRIQANKAAIK
AMIINRIGDFSLLIGIILIFANYKSVDY---ATVAILSPFFK---NS--SASFLNFNLEL
LTTIGIFLFLGAVGKSAQLGLHTWLPDAMEGPTPVSALIHAATMVTAGVFLIVRSSFIY-
EHTFSVLEFIAILGATTSLFASTTGLLQNDLKRVIAYSTCSQLGYMVFACGLSNYSVSVF
HLSNHAFFKALLFLSAGSVIHAVNDEQDMRKMGGLKNLVPFTYSTMIIGSLTLM-----G
FPF-----LA---GFYSKDLILEVAYGKYNSTGHFCYFLGTTGAFLTAFYSIRLIYLTFL
SKPT------------GFKKTICC--------------A-LDSGQSICIALGCLAILSLF
IGYLTKDLLVG---LG-SDFF-GNAIYVSV-KNLNMF-DAEF-IPSFYKTLPVSLSLCGL
LLGFVLYNFKSKE-LFLL------KISDFGKKYYSFLNRKWFFDKIYNNCLGQVFFRFGY
STSYKFIDRGIFETLGPTGLSLVSLRIASNLHKTQTGYLYHYTLAILIGLTLFISIRQML
---FFMEHLLDYR--
>YP_009245579_NAD5_Diphylleia_rotans
-------MYLSIIVLPGIAALLAGFF----GRFLGPKGSSLITTTLVGITCFLSLTSLYE
VGI--LGSVCEIKL-AKWIESDLLDADWGFLFDSLTVSMLVVITSISTLVHLYSIDYMSE
DPHLPRFMSYLSFFTFLMIILVTADNYLQMFVGWEGVGLCSYLLINFWFTRIAANKSAMK
AIIVNRIGDLGLILGIICLMTTFKSVDY---ETIFSIV-------SG--SDLITGVPSSI
LTLTTLLLFIGAVGKSAQIGLHTWLPDAMEGPTPVSALIHAATMVTAGVFLIIRSSPLF-
IVAEDILVLMTIMGSLTALFAATSGLLQNDLKRVIAYSTCSQLGYMVFACGITSFATSFF
HLANHAFFKALLFLSAGAVIHALGDEQDMRKMGGLIRLLPFTYVMILIGSLSLM-----G
FPF-----LT---GYYSKDVILEFAYSFVSFDSSFAYWLGTISAFFTAFYSIRLIYLTFI
TETN------------AYHSIIKN--------------V-HEVPLKMAIPLMILAFGSIF
IGYLTKDLIIG---FG-TPFW-NNAIVLLP-N------DAEF-LPFFIKLIPVIFSLFGA
FLGYTLYSSNILW-NMP-------EQGSVLRFVYSFFNQKWYFDQLYNRFCVQPLLKIGY
HTTFKLLDKGIIEILGPYGIASIIRYTVDRARYLQSGLIYHYTFIMVSGLILLLLVISQK
EF-SIITPFPI----
>ADU04611_NAD5_Mesostigma_viride
-------MYGTIIALPLIGSFCAGAL----GRWLGFRGAAFITTLFVTISTIFSWIAFYE
VAL--SGSFCYIPV-GAWILSEMFDVSWLFLFDPLTVVMLIVVTSVSTLVHLYSISYMWS
DPHLPRFMSNLSIFTFFMLILVTAGNLVQMFLGWEGVGLASYLLINFWFTRLQANKSAIK
AMLMNRVGDVGLALAIMGIFYTFQSVDF---SVLLALSASEH---SS--------SSENF
FTFVSLLLFVGSIGKSAQLGLHTWLPDAMEGPTPVSALIHAATMVTAGVFLIARCSALF-
EQSVHASFLVALVGAMTAFFAATTGAAQNDLKRVIAYSTCSQLGYMVFASGLSHYDLSVF
HLMNHAFFKALLFLGAGSVIHALSDEQDLRRMGGLVQLLPFTYTMMLIGSLSLV-----G
IPF-----LT---GFYSKDAILEVAYAKYTVSAHFAYWLGSLSVLFTSTYSFQLLQSAFL
TSTN------------AKRFALAK--------------I-HESDLLLSIPLVLLALGSIF
VGYLAKDMMIG---LG-TDFW-SQSLFVQP-YNVRLL-EAEFATPISIKMVPLIFSFLGA
LIAHLLQNNKNVQ-TFIFSLLLFNPATDFLRSIYSFWLKRWLFDKVYNDFIVRPCMKFGY
YVTFKTIDKGFLEWLGPLGISYTIQKFGSTLSRFQSGYIYHYAFVMLIAFTTFISMQTPF
---AESTFQLQST--
>YP_009092497_NAD5_Cyanoptyche_gloeocystis
-------MHLTVIILPLLSFVINTLF----GRSLGNSKIAIISTFNMAVTFIISVILFYE
TSI--IGQVLNIDV-SQWIRSEYLIVNWGFLFDQLTTTMLIIVNSISFLVHLYSTEYMRN
DPHLGRFMSYLSFFTFFMLILVTANNYLQMFVGWEGVGLCSYLLINFFSARIQANKAAIK
AMVLNRIGDFGLTIGMLMLLFVFKSLDY---SIIFSLAPYYT---EE--YIIIMNKSFSV
LDLTTFFLLIGAIGKSAQLGLHTWLPDAMEGPTPVSALIHAATMVTAGVFMIIRSSPLF-
EFAPTTLFTISIVGCLTAFFAASTGLLQNDLKRVIAYSTCSQLGYMVFACGVSAYNVAIA
HLFNHAFFKALLFLTAGSIIHSLADEQDLRKMGGLTQLLPLAYVMMLIGSLSLM-----G
```

```
FPF-----LT---GFYSKDFILEVTLSKYTINSAFCYWLGTLSAFFTAYYSFRLLYYTFL
SKPN------------GYKSTYNN--------------I-HESNITMVLPLCALGFLSIF
VGFAFKDMMVG---IG-TNFW-DSSIFILP-SIMNHINDAEN-IQLFLKWLPVIFSILGA
MLALLVYNYLKN---ILL------SNLIMYKALYSFFNKKWYFDKIYNEIIAQYISMMGY
HYTFKTLDRGIIEIIGPFGIADTFIKISKSTSFLQSGYLHHYIYTIFFGAITLVILTSNY
---IKLQFWTDLR--
>YP_010007659_NAD5_Cyanidium_caldarium
-------MYLIILFLPFCGSLFSILF----GRWVGGFGSSVITCLCLILAIFLSCIAFYE
VGI--LNYPCYIKL-TSWIDSGIFHVSWGFLFDSLTVTMIVIVSVISLLVHIYAIEYIRL
DPHLPRFISYLSVFTFFILILVTANNFIQIFLGWEGVGFASYLLINFWFTRLSANKAALK
AIVINRIGDVGLSLGILVIFLKFHSVDY---LTVFNLVPNVF---FE--SFNFFNIKVNF
CTIICLLLFIGVIGKSAQIGLHTWLPDAMEGPTPVSALIHAATIVTAGVFLIIRCSPLF-
EFSDTALFIVAIIGGITAFFSASIGLFQYDLKKIIAYSTCSQLGYIVFSCGLSNYSVGFF
HLFNHAFFKALLFLSAGSVIHAFLNEQDIRFIGGLKNLLPFTYAIMIVGSFSLL-----G
FPF-----IT---GFYSKDLILESAYNNFNIGSDFIYWLGLITVLLTAFYSFRLLFLVFL
NYPN------------FSTLIFNK--------------V-NDCSFYIKTSLTFLAIGSIF
LGYLTKEMLVG---MG-TDFW-DISLFFKL-NHIDFV-ENEY-LYFLIKLIPIFFSFFGI
LFCYLFNYFINL---KFL------EVQLRFRYFLFFLNKKWGWDYIYT-YIAYQFMNFGY
YISFKLIDKGILEFFGPQSLMKLITNMLNKLKFIQTGQITHYVFIIILGALVLFTLVELL
---DVLVYFFNIK--
>BBU60045_NAD5_Cyanidioschyzon_merolae
-------MYLTTVFAPLLGFLTCMLF----GRFLGKIGACFIVCFCTAISLLFSIIIFYE
VCI--ADYTNNIFF-TLWLSTGLLEIPWGFLFDPLTAVILLTINFVSLLVNVYSVEYIGE
DPHQVRFIAYLNIFAFFITILVTANNFLQIFLGWEGVGLASYLLINFWFTRLLASKSAIK
AIIINRVGDFFLSLAIFLIFLKYKSLNY---DIVFSITPFMS----N--YIECLGIKFSY
LDMISLFLFLGAIGKSAQIGMHIWLPDAIEGPTPVSALIHAATMVTAGVFLIVRCSPIL-
EYSSNVLIFISVIGASTSLFAGSVAIFQNDLKKVIAYSTCSQLGYMIFACGISNYIPSIF
HLVNHAFFKALLFLSAGYIIHSLFNEQDIRRIGSLLRILPIAYLMIMIGSLSLI-----G
IPF-----LT---GFYSKDLILELSFTSYFIINDFVYYLAIVSACLTAIYSTRLILLTFI
TKPN------------FSKISLIK--------------L-HYSSNWITLPLLILAFASIC
FGFILKDIFVG---LG-TNFW-GSSILINY-NHIYFI-EAEN--PIIIKWIPFFASIIGF
FIIIIMQNNYA---IFSF------SFVRYMHSFFYFFNKKWYLDKIII-IIANSVLNFGY
YVSLKIFDKGVIEYFGPFGLIKIIPYMSKQSRNLQSGQIAHYVFIILIGLLVLIIVIEFN
---NLYFIINYIK--
>AIA61058_ND5_Cyanidiaceae_sp._MX-AZ01
-------MHLTVILAPLIGFLTCGLF----GRFIGKKGANFISCFCICVSLLFSIIIFYE
ACF--SNYKNHILL-SSWIDSAFLEVSWGFIFDSLTAVILLIVNFISLLVIIYSIEYIQE
DPHQIRFISYLTIFVFFITILVTANNFMQLFLGWEGVGLASYLLINFWFTRLQANKSAIK
AIIINRVGDFFLSLGILFIFIKFQSLNY---NVVFSIVPFIT---DN--FLECYGIKFSY
LDIITLLLFLGAMGKSAQLGIHIWLPDAIEGPTPVSALIHAATIVTAGVFLVVRCSPIF-
EFSNFTLVFVSVVGALTSFFAGSVAIFQNDLKKVIAYSTCSQLGYMIFVCGISNYTSSIF
HMVNHAFFKALLFLSAGYIIHSLINEQDIRRIGALMRILPIAYIIMLIGSLSLI-----G
IPF-----IT---GFYSKDLILELIFESYFVIHDFVYWLAIVSVFLTAMYSIRVILLVFV
KNSN------------TARINLIN--------------L-HYSSFFITSPLFILAFASIC
FGFLLKDIFVG---LG-TNFW-GSSVIIYY-SQVNFL-EVEN-LSTVIKWLPFIISIIGV
FLILLSINYD----LFRL------FPLEKIHTIFYFFNKKWYFDKIIT-TIANFILSFGY
DISLRIFDKGIIEYFGPFGLIKIVPYFSIRSKNLQTGQIAHYIFVILIGFLSLVLFTEFL
---NIINLIHFEK--
```

```
>YP_009968183_ND5_Cyanidiococcus_yangmingshanensis
-------MYLITIFAPLLGFIICGLF----GRFLGKKGACFITYFCSSISLVFSIIIFYE
VSV--AAYDNHILL-AKWVESSIFFVSWGFNFDSLTALILVMVSFISLLVNIYSIEYIEE
DPHQVRFISYLNIFVFFMFMLVTANNFIQLFLGWEGVGLSSYLLINFWFTRLIANKAAIK
AIIINRVGDFFLSLGIFFIFFKFKSLDY---NVVFSIAPSII---DN--YLECLGIKFLY
LDVVTLFLFLGAIGKSAQIGIHMWLPDAIEGPTPVSALIHAATMVTAGVFLIVRCSPIF-
EFSHFTLVIISVVGALTSLFAGSVAIFQNDLKRVIAYSTCSQLGYIMFACGISNYMASIF
HLLNHAFFKALLFLSAGYIIHSLANEQDIRRIGSLIRIIPIAYIAIIIGSLSLI-----G
IPF-----LT---GFYSKDLILELTFGSYFIIHDFTYWLAIISASMTAIYSMRLVLLTFI
KSTN------------VIKINLIH--------------I-HYSSSWMIVPLVILIIGSIY
FGFILKDIIIG---LG-TDFW-GSSILTHY-SRANFV-EVEN-LHVIIKWLPFFVSILGV
VIILFIYNYDFK--IFST------IFLKYTHSFFYFFNKKWYFDKIAN-IIAIYLLSFGY
NVCLKVFDKGIIEYFGPFGLMKIVPYLSTESKSLQTGQIAHYIFVILMGLLFLILSTEFI
---NIFYIINYIK--
>YP_008999866_NAD5_Prasinoderma_coloniale
-------MYLVLLLLPTLGTLFGCFF----GRAMGGRGTAILTTTAVALSALLSFVGLYE
VGY--AGSPCALSL-GRWVHSEAFDAEWGFLFDTLTMVMLCMITGVSSLVHLYSIGYMSA
DPHLPRFMTYLSAFTIFMMLLVTGNNFLILFLGWEGVGLASYLLINFWFTRAQANKASIK
AMIMNRVGDVGLALGIFGIYALFKSLDY---ATVFSTAHLHA---ED--VLIFFGYEVHA
LTAIGCLLFIGAIGKSAQVGLHTWLPDAMEGPTPVSALIHAATMVTAGVFLLARVSPLL-
AYAPGALQVVTVVGATTALLGAVTGLTQNDMKRVIAYSTCSQLGYMVFAAGVGAYAVALY
HMIIHAFFKALLFLCAGSVIHAIGGEQDMRKMGGLARLLPLTYACMVVGSLALV-----G
FPY-----LG---AFYSKDLVLEAAYSRGMATGLLAYLLGIFAAFCTSYYSFRLLFLVFW
GESR------------TPKAGVRL--------------A-HEGSWTMLIPLLVLTTCSIF
GAEGLKEAFTG---LG-TPFW-GGSLVAAG-GLTQSF-EAEL-VPLSAKLAPLVATGLGA
TLAYLVHLGPLRKLALKA-----LTESQLGRTFYIFTNKRWFIDKIYAEVFGWSALFFGY
HVTFKTIDKGLLEVFGPTGAKFSAASLIPALRLRSWTSLPHYASVLAFCLILLLSILGPA
S--GMLSLGIGP---
>BAA78087_NAD5_Dictyostelium_discoideum_3D
-------MYIVNLILPLIGSIITGIF----GHKLGNRISIKIAVGCMMLTAISSLYIGYE
ILL--CNSVVHFKL-GTWMQVGSLNVEYGLLYDSLTSIMIIVITCISSMVHLYSMDYMKE
DPHKTRFFSYLSLFTFFMMLLVTADNFVQLFFGWEGVGIMSYLLINFWYTRLQANKSALK
AVILNRFGDFGLFFGILLVFLVFKSVDF---SVIFTIAPFIT---EY--TINLLGYEVNA
ITLIGSFIVIGVVGKSAQLGLHMWLPDAMEGPTPVSALLHAATMVTAGVFLVLRTSPLL-
SYSITILNILTIIGALTTLFATTIGIVQNDIKRVIAYSTCSQLGYMIFACGLLNYNASIY
HLTTHAFFKALLFLSAGSVIHGLNDEQDMRKMGGLVNLMPLTYQCMLIGTLALT-----G
FPF-----LS---GYYSKDIILETSYATYYWEGTFAAIIGYVAAFGTTFYSFRLLILTFF
NKPR------------MQYKTIAG--------------V-HEASTNMVIPLVILALCSIF
IGYVTKDFFVG---LG-TPVW-NNSFFAYP-YNNLIL-ESEV-LQRELKLLPLFAFIYGV
ITPVLFYFNIKE---IDR--MINVKQNLMVKESYFFFVKKWYFDFLSRVLIVVPFFHLSY
DVMNKNLDKGLWEKIGVTGVAKNEITLSTYISYI----VQTIILIIVVGIFSFMTGFIYM
ELCIIIGILYICLPS
```
